# Full-length transcript sequencing of human and mouse identifies widespread isoform diversity and alternative splicing in the cerebral cortex

**DOI:** 10.1101/2020.10.14.339200

**Authors:** A.R. Jeffries, SK. Leung, I. Castanho, K. Moore, J.P. Davies, E.L. Dempster, N.J. Bray, P. O‘Neill, E. Tseng, Z. Ahmed, D. Collier, S. Prabhakar, L. Schalkwyk, M.J Gandal, E. Hannon, J. Mill

## Abstract

Alternative splicing is a post-transcriptional regulatory mechanism producing multiple distinct mRNA molecules from a single pre-mRNA. Alternative splicing has a prominent role in the central nervous system, impacting neurodevelopment and various neuronal functions as well as being increasingly implicated in brain disorders including autism, schizophrenia and Alzheimer’s disease. Standard short-read RNA-Seq approaches only sequence fragments of the mRNA molecule, making it difficult to accurately characterize the true nature of RNA isoform diversity. In this study, we used long-read isoform sequencing (Iso-Seq) to generate full-length cDNA sequences and map transcript diversity in the human and mouse cerebral cortex. We identify widespread RNA isoform diversity amongst expressed genes in the cortex, including many novel transcripts not present in existing genome annotations. Alternative splicing events were found to make a major contribution to RNA isoform diversity in the cortex, with intron retention being a relatively common event associated with nonsense-mediated decay and reduced transcript expression. Of note, we found evidence for transcription from novel (unannotated genes) and fusion events between neighbouring genes. Although global patterns of RNA isoform diversity were found to be generally similar between human and mouse cortex, we identified some notable exceptions. We also identified striking developmental changes in transcript diversity, with differential transcript usage between human adult and fetal cerebral cortex. Finally, we found evidence for extensive isoform diversity in genes associated with autism, schizophrenia and Alzheimer’s disease. Our data confirm the importance of alternative splicing in the cerebral cortex, dramatically increasing transcriptional diversity and representing an important mechanism underpinning gene regulation in the brain. We provide this transcript level data as a resource to the scientific community.

## Introduction

Alternative splicing is a post-transcriptional regulatory mechanism producing multiple distinct molecules, or RNA isoforms, from a single mRNA precursor. In eukaryotes, alternative splicing dramatically increases transcriptomic and proteomic diversity from the coding genome and represents an important mechanism in the developmental and cell-type specific control of gene expression. The mechanisms involved in alternative splicing include the use of alternative first and last exons, exon skipping, alternative 5’ and 3’ splice sites, mutually exclusive exons and intron retention^1^. These phenomena are relatively common, influencing the transcription of >95% of human genes^2^. Importantly, because alternatively spliced transcripts from a single gene can produce proteins with very different, often antagonistic, functions^3,4^, there is increasing interest in the role of RNA isoform diversity in health and disease^5^; the correction of alternative splicing deficits has been shown to have dramatic therapeutic benefit in spinal muscular atrophy^6^. Alternative splicing appears to be particularly important and prevalent in the central nervous system^7^, where it impacts upon neurodevelopment^8^, aging^9^ and key neuronal functions^10^. Of note, mis-splicing is a common feature of many neuropsychiatric and neurodegenerative diseases^11^ with recent studies highlighting splicing differences associated with autism^12^, schizophrenia (SZ)^13^ and Alzheimer’s disease (AD)^14^.

A systematic analysis of the full complement of transcripts across tissues and development is an important step in understanding the functional biology of the genome. For example, transcript-level annotation can be used to improve the functional consequences of rare genetic variants^15^. Current efforts to characterize RNA isoform diversity are constrained by the fact that standard short-read RNA-Seq approaches cannot span full-length transcripts, making it difficult to characterize the diverse landscape of alternatively spliced transcripts^16^. Recent advances in long-read sequencing technology have enabled the full complement of RNA isoforms to be quantified^17^. Pacific Biosciences’ (PacBio) single-molecule real-time (SMRT) sequencing and Oxford Nanopore Technologies (ONT) nanopore sequencing, for example, both generate reads >10Kb, enabling the direct assessment of alternatively-spliced transcripts^17^.

In this study, we systematically characterize RNA isoform diversity in the cerebral cortex, a key region of the brain involved in perception, cognition and consciousness. We used the Pacific Biosciences long-read isoform sequencing (Iso-Seq) approach^18^ to generate full-length cDNA sequences from the human and mouse cortex. We identify widespread transcript diversity including alternatively spliced transcripts not previously described in any existing genomic annotations, and evidence for novel transcripts of genes robustly associated with neuropsychiatric and neurodegenerative disease. We subsequently used short-read RNA-Seq and nanopore sequencing from the same samples to validate and complement our Iso-Seq data. Importantly, we find widespread evidence of different alternative splicing events, and examples of fusion genes representing read-through transcription between adjacent genes. A comparison of human and mouse cortex identified evidence for species-specific transcript diversity at several genes, and a comparison of fetal and adult human cortex highlights developmental changes in splicing and transcript expression. Our data confirm the importance of alternative splicing in the cerebral cortex, dramatically increasing transcriptional diversity and representing an important mechanism underpinning gene regulation in the brain. Our transcript annotations and data are available as a resource to the research community (see **Web Resources**).

## Results

### Harnessing long-read RNA isoform sequencing to characterize the transcriptional landscape of the human and mouse cortex: methodological overview

We first generated Pacific Biosciences SMRT Iso-Seq datasets using high-quality RNA isolated from i) human cortex (n = 7) dissected from fetal (n = 3, mean age = 16 weeks post-conception (WPC), range = 14 – 17 WPC) and adult (n = 4, mean age = 61.8 years, range = 24 – 89 years) donors and ii) mouse cortex (n = 8, mean age = 6 months, range = 2 – 12 months) (see **Supplementary Table 1**). Following library preparation and SMRT sequencing (see **Methods**), a total of 5.14M circular consensus sequence (CCS) reads were generated from the human cortex samples (human fetal cortex: 2.70M CCS reads; human adult cortex: 2.44M CCS reads) and 4.32M reads were generated from the mouse cortex samples (**Supplementary Table 2**). There was a similar distribution of CCS read-lengths in both human and mouse cortex samples with the majority of reads sized 2 - 3kb in length (human adult cortex: mean length = 2.3kb; human fetal cortex: mean length = 3.1kb; mouse cortex: mean length = 2.6kb) (**Figure 1**, **Supplementary Figure 1**), corresponding to the mean length of mRNA in the human^19^ and mouse^20^ reference genome. Raw reads were processed using the *Iso-Seq* (v3.1.2) pipeline^18^ (see **Methods**). Briefly, high-quality polished reads were filtered, mapped to the genome (human: hg38; mouse: mm10), and clustered using *cDNA Cupcake* scripts^21^ followed by *SQANTI2*^22^ annotation and subsequent filtering to stringently remove potential artifacts (see **Methods** and **Supplementary Table 3**). Rarefaction curves confirmed that the Iso-Seq datasets approached saturation, indicating that our coverage of RNA isoform diversity was representative of the true population of transcripts (**Figure 1**, **Supplementary Figure 2**). All downstream analyses and statistics reported in this paper were based on the subset of *SQANTI2*-filtered transcripts unless otherwise indicated, although both unfiltered and filtered datasets are provided as a resource and genome browser tracks to the community (see **Web Resources**). We subsequently complemented our whole-transcriptome Iso-Seq data with short-read RNA-Seq (Illumina) data and additional full-length transcriptome data generated from nanopore sequencing (ONT) on an overlapping set of human cortex samples. An overview of the methods and datasets used in our analysis is given in **Supplementary Figure 3**. Taken together, our analysis represents the most comprehensive characterization of full-length transcripts and transcript diversity in the human and mouse cortex yet undertaken.

**Figure 1:**
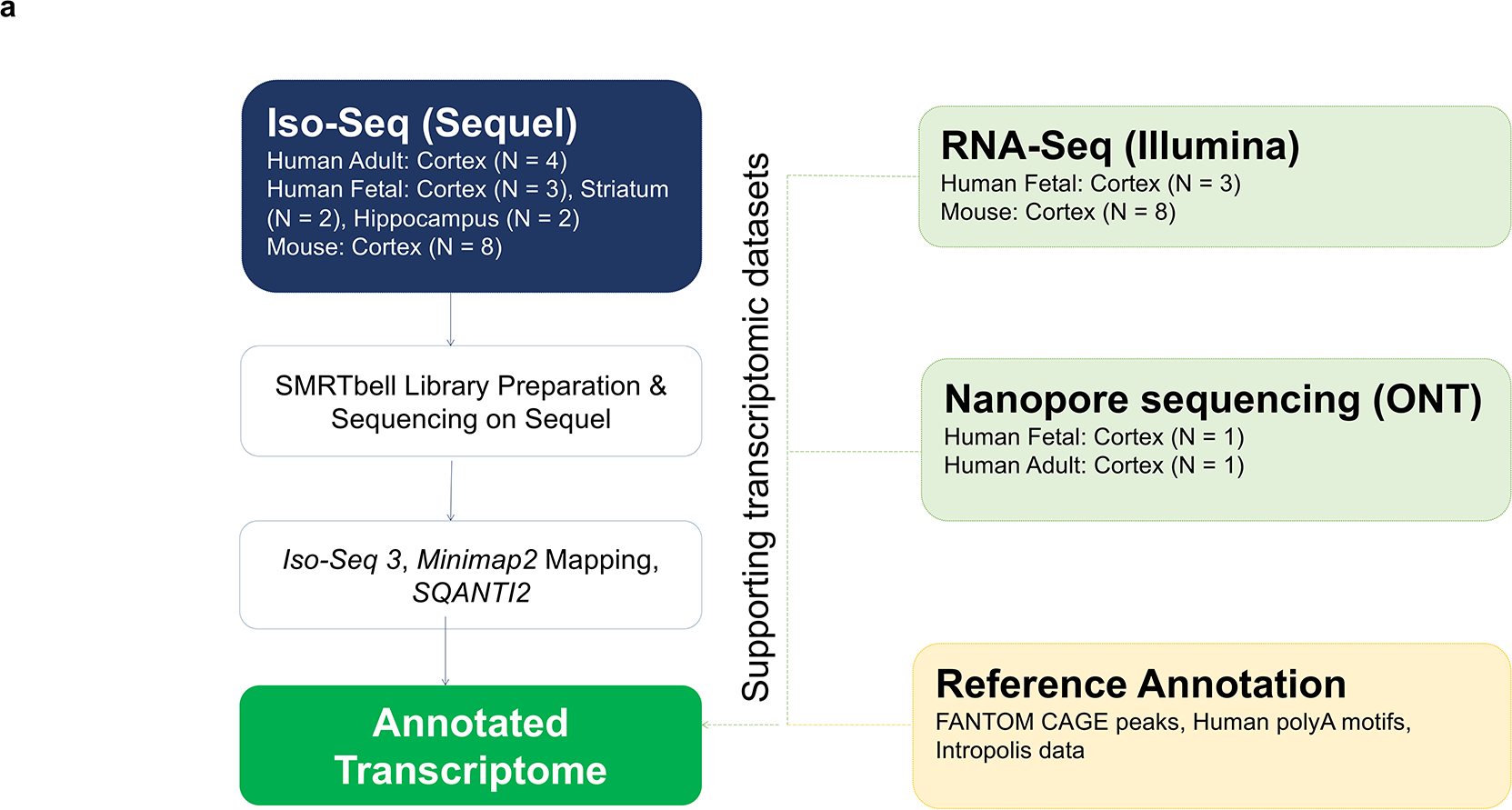
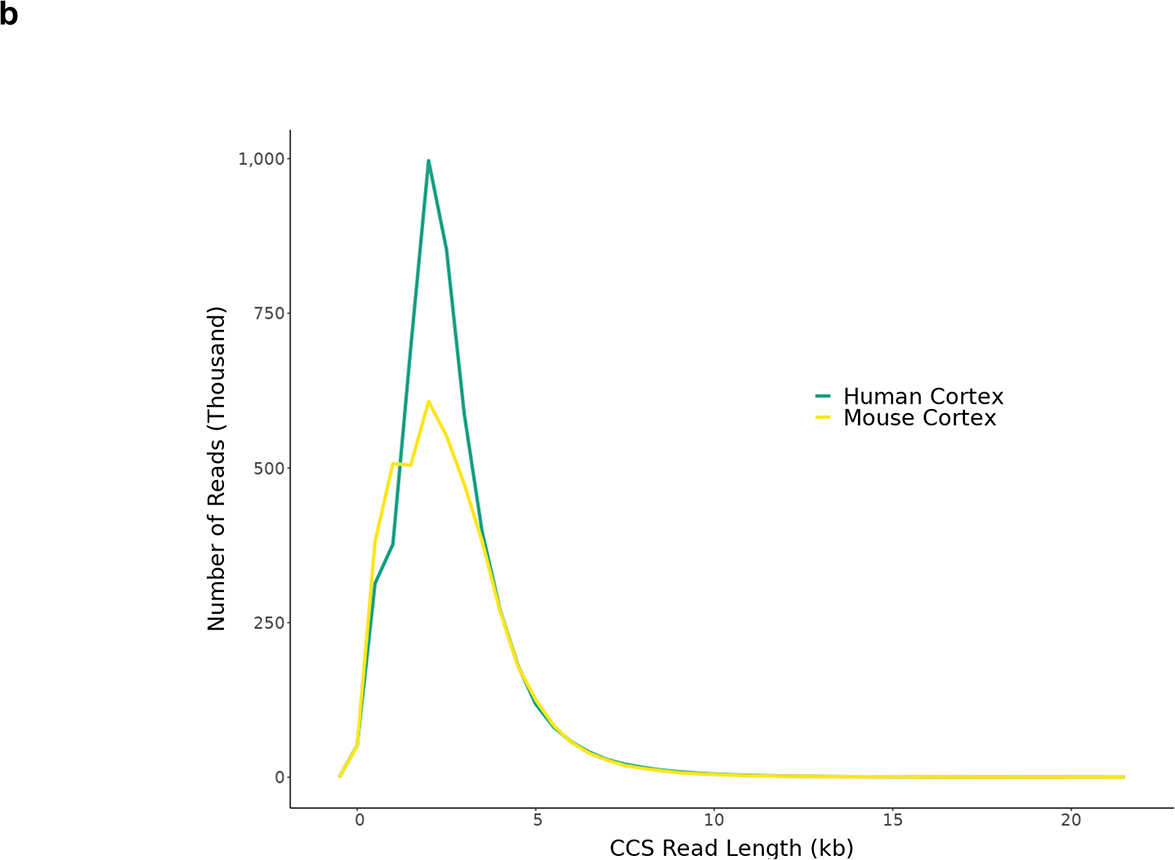
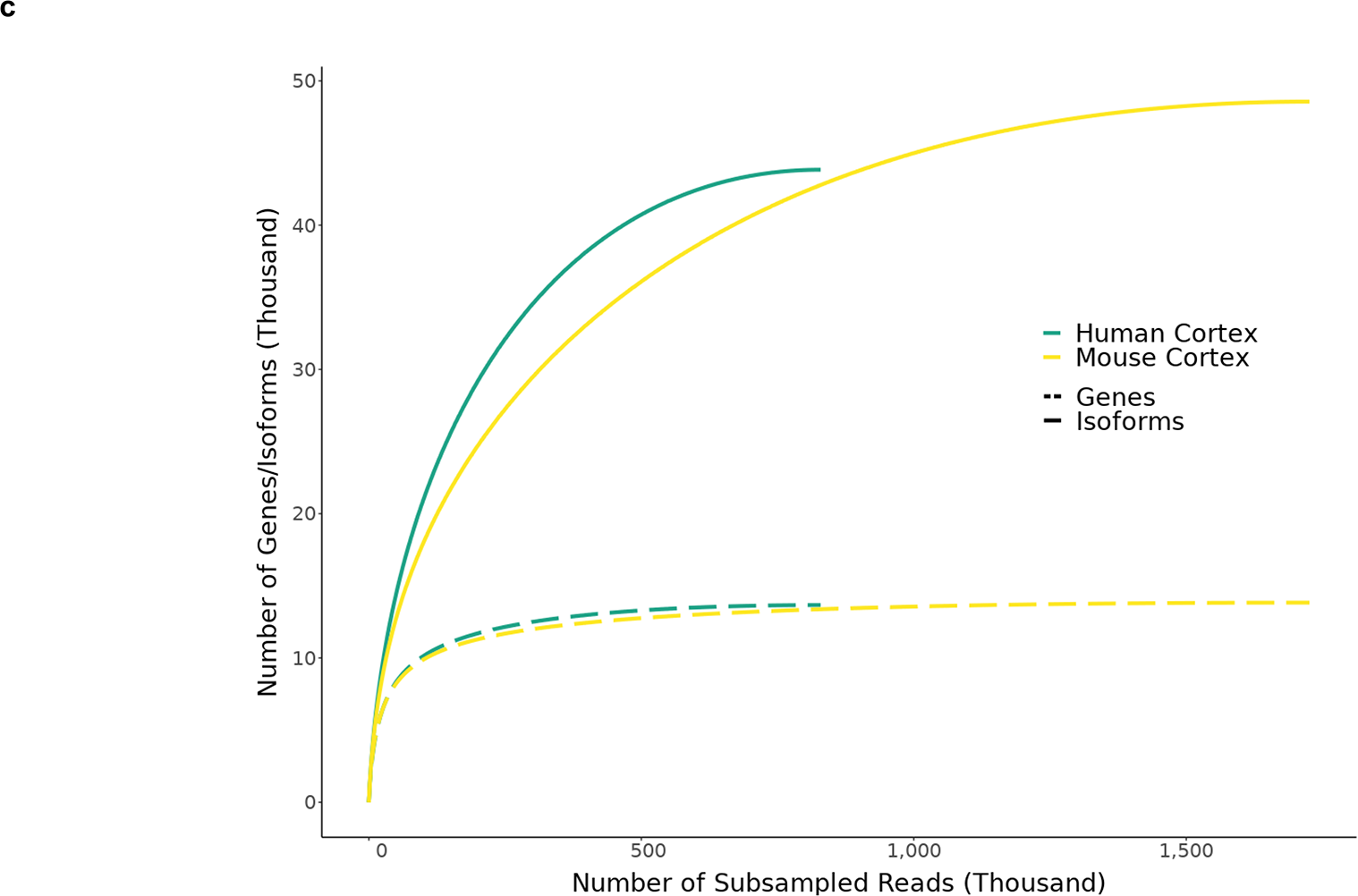
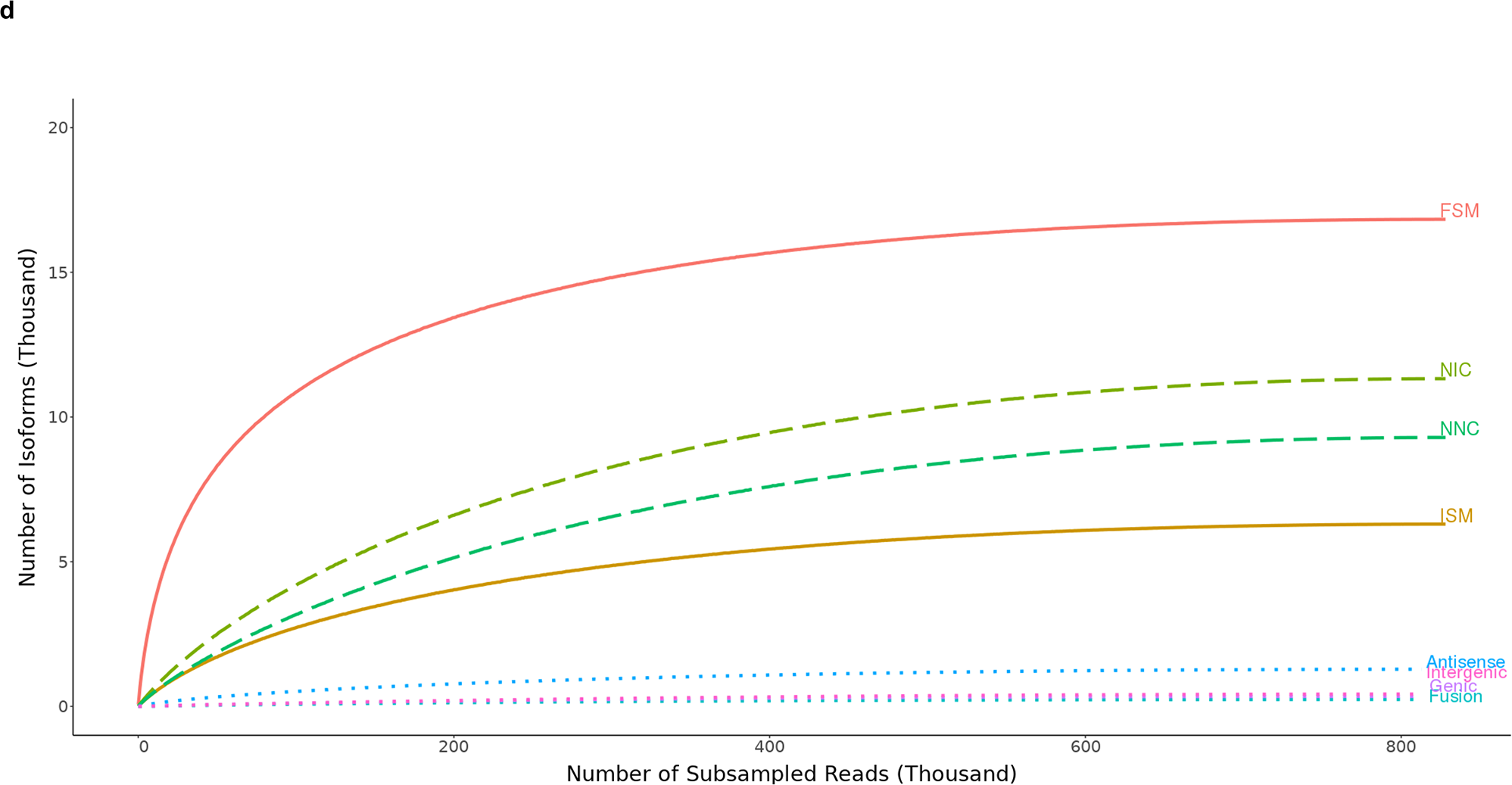
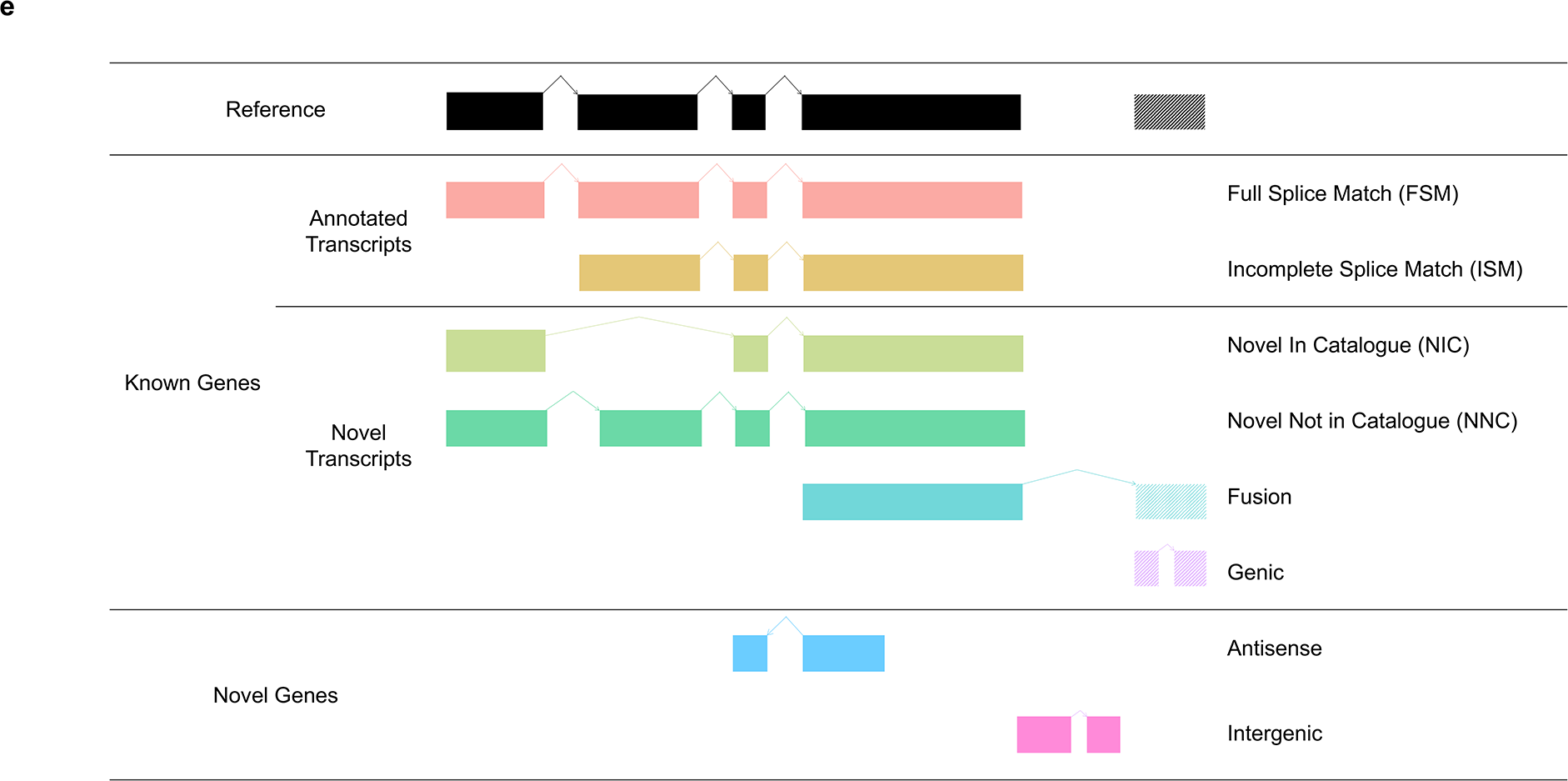
Generation of high-quality long-read transcriptome datasets for human and mouse cerebral cortex. **a)** An overview of the analysis pipeline used to comprehensively annotate the human and mouse cortical transcriptome using the Pacific Biosciences Iso-Seq approach. Iso-Seq data was complemented with short-read RNA-Seq (Illumina) and nanopore sequencing (ONT) for a subset of the same samples, and compared to existing genomic annotation data. Further details on the pipeline can be found in **Supplementary Figure 3** and details on the samples in **Supplementary Table 1**. **b)** The distribution of CCS read lengths in our human and mouse cortex datasets. The distribution of CCS read lengths for individual samples can be found in **Supplementary Figure 1**. **c)** Iso-Seq rarefaction curves at the gene (dotted line) and isoform (solid line, minimum two FL reads) level for human (n = 7 biologically independent samples) and mouse (n = 8 biologically independent samples) cortex. **d)** Iso-Seq rarefaction curves for SQANTI2 classification categories (see **Figure 1e**) in the human cortex dataset. **e)** An isoform was classified as ‘FSM’ if it aligned with reference genome with the same splice junctions and contained the same number of exons, ‘ISM’ if it contained fewer 5’ exons than the reference genome, ‘NIC’ if it represented a novel isoform containing a combination of known donor or acceptor sites, or ‘NNC’ if it represented a novel isoform with at least one novel donor or acceptor site. Rarefaction curves for these isoform categories for mouse cortex can be seen in **Supplementary Figure 2**. CCS – Circular consensus sequence, FL - Full length, FSM – Full Splice Match, ISM – Incomplete Splice Match, NIC – Novel In Catalogue, NNC – Novel Not in Catalogue. ONT - Oxford Nanopore Technology, SMRT - Single molecule real-time

### SMRT sequencing can accurately quantify patterns of gene expression in the human and mouse cortex

Following stringent quality control (QC) of our data, SMRT sequencing reads mapped to 12,832 (human cortex) and 13,450 (mouse cortex) ‘annotated’ genes already present in existing genomic databases (**Table 1**, **Supplementary Figure 4**). Gene expression patterns from Iso-Seq reflected expected transcriptional profiles for the brain regions profiled. Using the Human Gene Atlas database^23^, for example, we found that the most abundantly-expressed genes (top 500, ranked by transcripts per million (TPM)) in the human cortex Iso-Seq dataset were most significantly enriched for ‘prefrontal cortex’ genes (odds ratio = 5.91, adjusted P = 1.93 ×◻10^−35^) (**Supplementary Figure 5** and **Supplementary Table 4**). Likewise, using the Mouse Gene Atlas database^23^, we found that the most abundantly-expressed genes in the mouse cortex Iso-Seq dataset were most significantly enriched for ‘prefrontal cerebral cortex’ (odds ratio = 5.47, adjusted P = 1.58 ×◻10^−19^) (**Supplementary Figure 5** and **Supplementary Table 4**). Although the Iso-Seq method has been shown to be accurate at characterizing the diversity of RNA molecules present in a sample^24^, its sensitivity for quantifying levels of gene expression has not been systematically explored. We therefore generated highly-parallel short-read RNA-seq (Illumina) data on human fetal cortex samples (n = 3, total 153M mapped reads), which represent a subset of the human cortex Iso-Seq data, and mouse cortex (n = 8, total 128M mapped reads, **Supplementary Table 5**) samples, finding a strong correlation between gene level expression levels quantified using the two methods in both datasets despite the relatively low number of sample comparisons (human fetal cortex: n = 9,223 genes, corr = 0.58, P < 2.23 ×◻10^−308^; mouse cortex: n = 12,978 genes, corr = 0.80; P < 2.23 ×◻10^−308^, **Figure 2**, **Supplementary Figure 6**).

**Table 1:** An overview of the whole transcriptome Iso-Seq datasets generated on human and mouse cerebral cortex. FSM – Full Splice Match, ISM – Incomplete Splice Match, NIC – Novel In Catalogue, NNC - Novel Not in Catalogue

|  | Human Cortex | Mouse Cortex | Adult Cortex | Fetal Cortex |
| --- | --- | --- | --- | --- |
| Unique Genes | 12881 | 13581 | 10978 | 9664 |
| Annotated Genes | 12832 (99.62%) | 13450 (99.04%) | 10949 (99.74%) | 9647 (99.82%) |
| Novel Genes | 49 (0.38%) | 131 (0.96%) | 29 (0.26%) | 17 (0.18%) |
| Isoforms | 42645 | 51159 | 27873 | 23213 |
| Genes with > 1 Isoform | 8599 (66.74%) | 9612 (70.73%) | 6386 (58.17%) | 5409 (55.95%) |
| Genes with > 10 Isoforms | 443 (3.44%) | 670 (4.93%) | 153 (1.39%) | 105 (1.09%) |
| Protein-coding Transcripts | 39372 (92.33%) | 47827 (93.49%) | 25702 (92.21%) | 21770 (93.78%) |
| Non Protein-coding Transcripts | 3273 (7.67%) | 3332 (6.51%) | 2171 (7.79%) | 1443 (6.22%) |
| Known Transcripts (FSM, ISM) | 28654 (67.19%) | 32664 (63.85%) | 20451 (73.37%) | 16947 (73.01%) |
| Novel Transcripts | 13991 (32.81%) | 18495 (36.15%) | 7422 (26.63%) | 6266 (26.99%) |
| *FSM* | 23629 (55.41%) | 27475 (53.71%) | 16971 (60.89%) | 14796 (63.74%) |
| *ISM* | 5025 | 5189 | 3480 | 2151 |
| *NIC* | 9724 | 10649 | 4903 | 4730 |
| *NNC* | 4013 | 7414 | 2380 | 1447 |
| *Genic Genomic* | 41 | 57 | 23 | 13 |
| *Antisense* | 35 | 78 | 20 | 9 |
| *Fusion* | 153 | 219 | 85 | 54 |
| *Intergenic* | 25 | 78 | 11 | 13 |
| *Genic Intron* | 0 | 0 | 0 | 0 |

**Figure 2:**
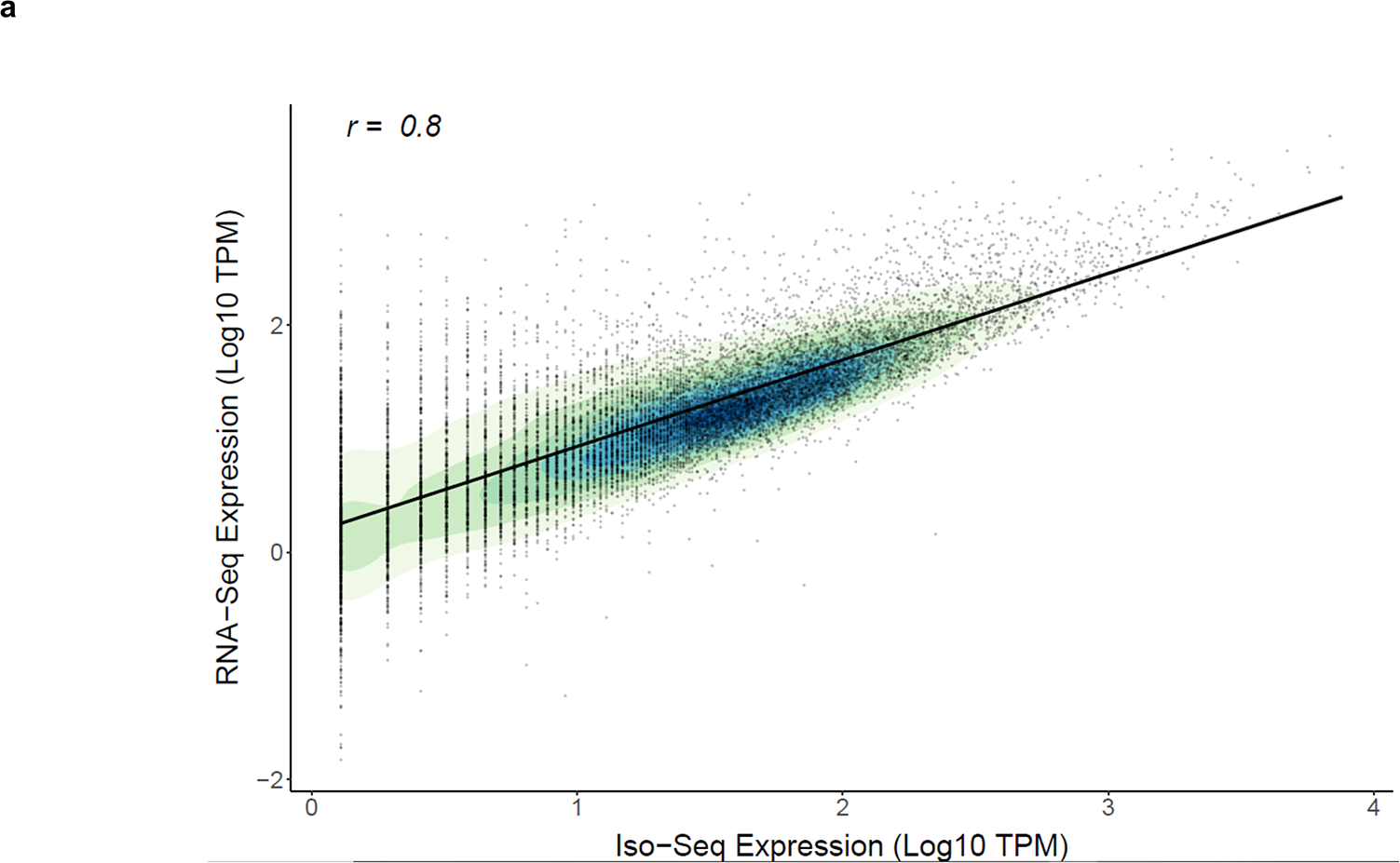
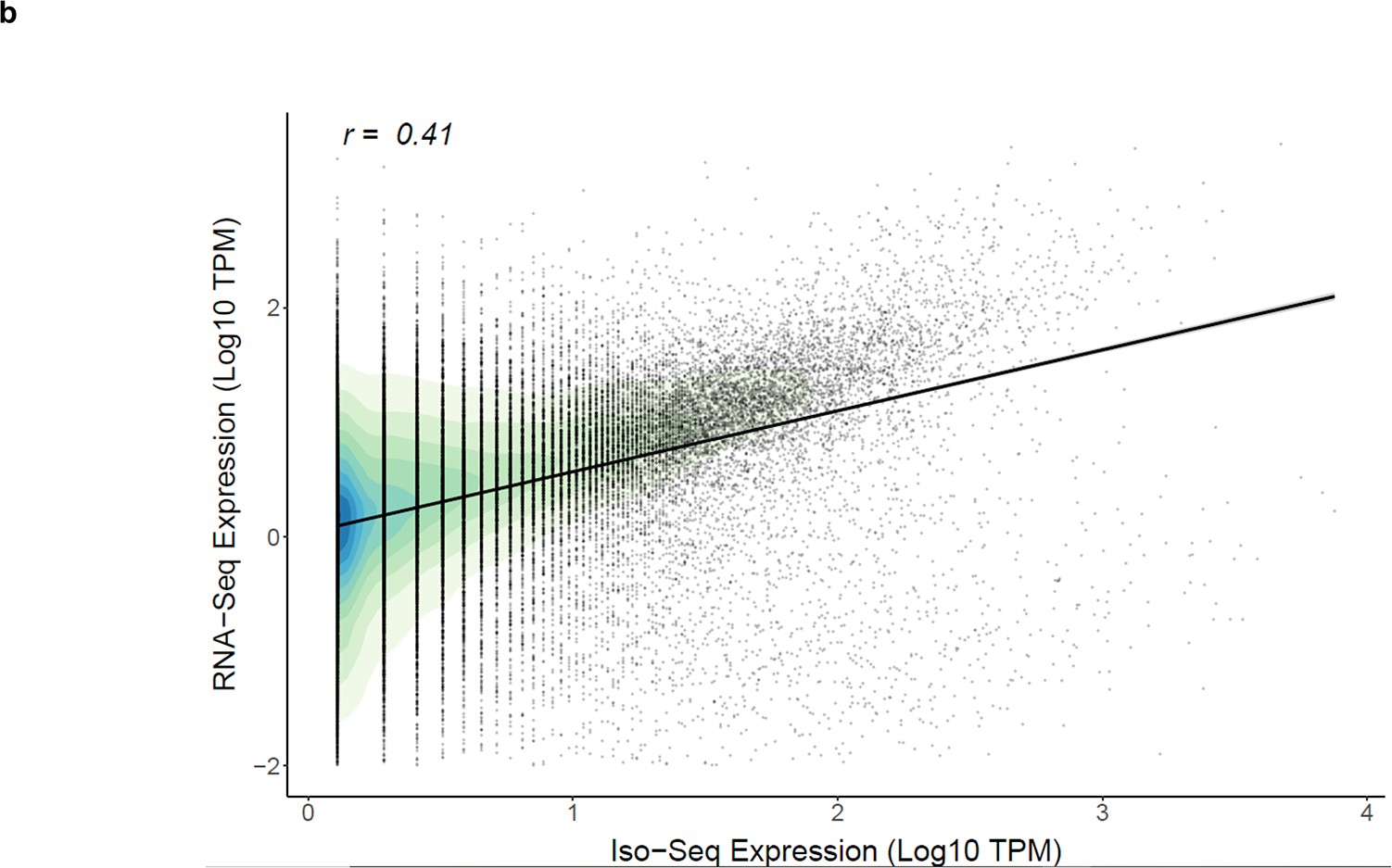
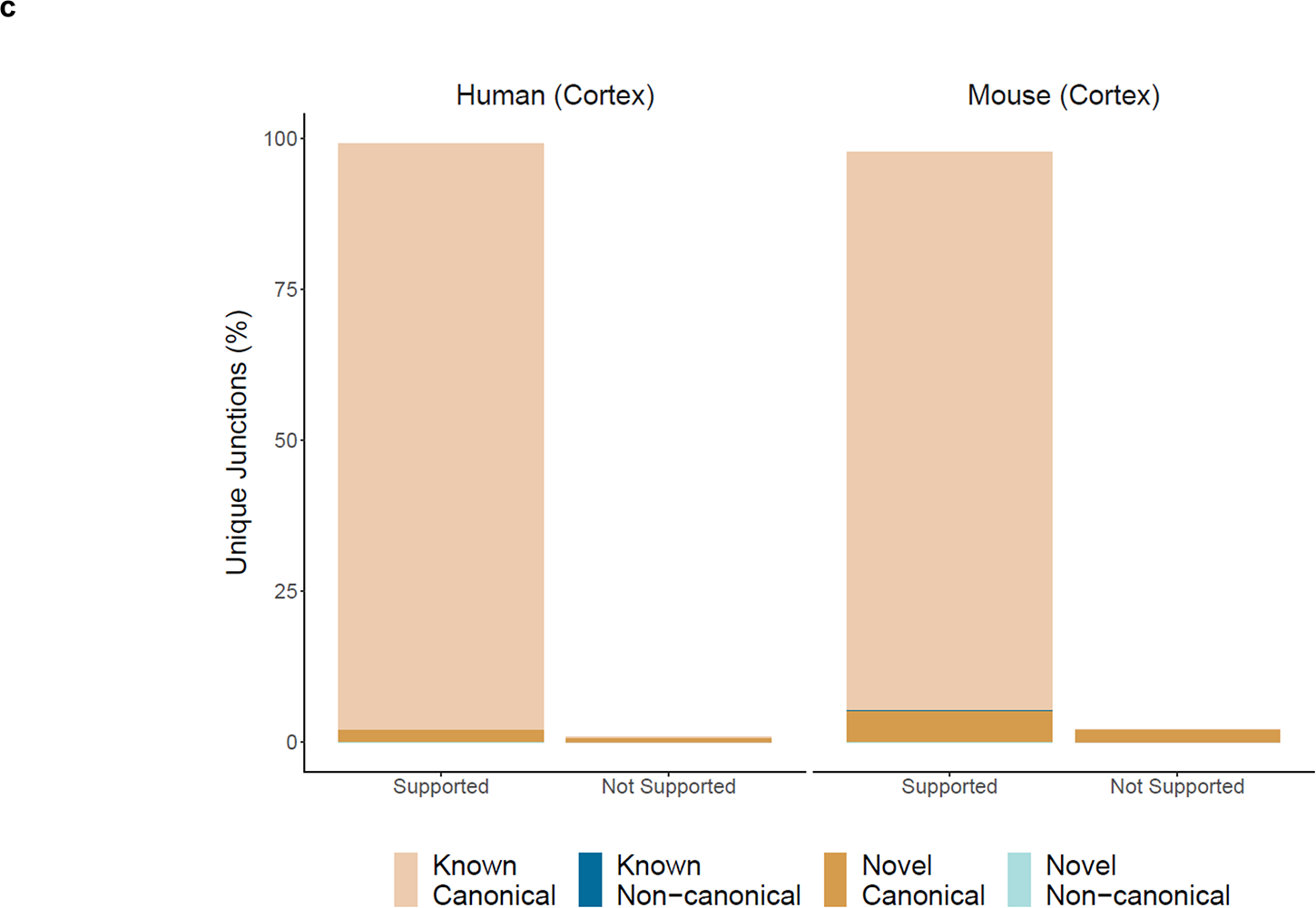
Long-read Iso-Seq data can accurately quantify levels of gene expression in the cortex. Shown is the relationship between expression estimated using RNA-Seq and Iso-Seq at the **a)** gene level (n = 12,978 genes, Pearson‘s correlation = 0.81, P < 2.23 ×◻10^−308^) and **b)** transcript level (n = 44,239 multi-exonic transcripts, Pearson‘s correlation = 0.41, P < 2.23 ×◻10^−308^, RNASeq threshold of 0.01TPM) in the mouse cortex (data derived from 8 biologically independent samples). The density of values is represented in increasing scale from light green to dark blue. Equivalent plots for the human fetal cortex are presented in **Supplementary Figure 6**. **c)** The proportion of unique splice junctions that were identified from Iso-Seq human (n = 7 biologically independent samples) and mouse (n = 8 biologically independent samples) cortical datasets and supported by RNA-Seq. Splice junctions were classified as novel if the junction is not present in the reference isoform, and non-canonical if it differs from GT/AG, GC/AG, or AT/AC. TPM – Transcripts per Million.

### Transcript-level analysis of gene expression identifies widespread RNA isoform diversity amongst expressed genes in the cerebral cortex

In total, we identified 42,645 unique transcripts (mean length = 2.6kb, s.d = 1.3kb, range = 0.082 – 11.8kb) in the human cortex and 51,159 unique transcripts (mean length = 2.9kb, s.d = 1.6kb, range = 0.08 – 15.9kb) in the mouse cortex (**Table 1**). As expected, transcripts were enriched near to annotated Cap Analysis Gene Expression (CAGE) peaks derived from the FANTOM5^25^ dataset, which facilitates the mapping of transcripts, transcription factors, transcriptional promoters and enhancers (human cortex: mean distance from CAGE peak = 542bp downstream, 30,978 (72.6%) transcripts located within 50bp of a CAGE peak, **Figure 3**; mouse cortex: mean distance from CAGE peak = 247bp downstream, 35,781 (69.9%) transcripts located within 50bp of a CAGE peak, **Figure 3**), and were also located proximal to annotated transcription start sites and transcription termination sites (**Figure 3**). For instances where transcripts could be confirmed using the available matched short-read RNA-Seq data using *Kallisto* (see **Methods**), there was a significant correlation between transcript expression levels quantified using both sequencing approaches (human cortex: n = 21,144 transcripts, corr = 0.36, P < 2.23 × 10^−308^; mouse cortex: n = 44,239 transcripts, corr = 0.41, P < 2.23 × 10^−308^, **Figure 2**, **Supplementary Figure 6**). The vast majority of unique splice junctions were supported by RNA-Seq in both human (n = 89,981 (99.2%) junctions) and mouse cortex (n = 138,032 (97.8%) junctions), and the majority of junctions that were not supported were novel and canonical (human cortex: GT/AG = 422, GC/AG = 90; mouse cortex: GT/AG = 2,390, GC/AG = 388, AT/AC = 3, **Figure 2**). As a resource to the community, our human and mouse cortical RNA isoform maps (including both filtered and unfiltered transcripts) are available to browse and download from http://genome.exeter.ac.uk/BrainIsoforms.html. There was a wide range in the number of discrete multi-exonic RNA isoforms identified per gene (human cortex: 1 - 87, mouse cortex: 1 - 79; **Supplementary Figure 7** and **Supplementary Table 6**), with the majority of genes (human cortex: n = 8,599 (66.7%), mouse cortex: n = 9,612 (70.7%)) characterized by more than one RNA isoform, and a notable proportion of genes characterized by more than ten isoforms (human: n = 443 (3.4%), mouse: n = 670 (4.9%), **Figure 3**). The gene displaying greatest RNA isoform diversity in human cortex was *MEG3*, a maternally expressed imprinted long non-coding RNA (lncRNA) gene involved in synaptic plasticity^26^ (87 isoforms, **Supplementary Figure 8**), and the most diverse gene in mouse cortex was *Tcf4*, a neurodevelopmental gene implicated in schizophrenia^27^ (79 isoforms, **Supplementary Figure 9**). Gene ontology (GO) analysis showed that the most enriched molecular function amongst the 100 most transcriptionally diverse genes in both human and mouse cortex was ‘RNA binding’ (human cortex: odds ratio = 3.17, adjusted P = 1.21 × 10^−3^; mouse cortex: odds ratio = 2.88, adjusted P = 1.69 × 10^−2^) (**Supplementary Table 4**), an interesting observation given the role that RNA-binding proteins (RBPs) themselves play in regulating tissue-specific patterns of alternative splicing^28^. In both human and mouse cortex, the number of detected RNA isoforms was correlated with gene length (human cortex: corr = 0.28, P = 1.71 × 10^−223^; mouse cortex: corr = 0.32, P = 3.17 × 10^−301^, **Supplementary Figure 10**) and the number of exons (human cortex: corr = 0.24, P = 2.42 × 10^−153^; mouse cortex: corr = 0.22, P = 2.44 × 10^−143^, **Supplementary Figure 11**). Of note, amongst ‘highly-expressed’ genes (> 2.5 Log_10_ TPM) there was a stronger relationship between isoform number and both gene length (human cortex: corr = 0.43, p = 2.62 × 10^−27^; mouse cortex: corr = 0.45, P = 2.42 × 10^−31^, **Supplementary Figure 10**) and gene exon number (human cortex: corr = 0.42, P = 3.75 × 10^−25^; mouse cortex: corr = 0.43, P = 1.45 × 10^−28^, **Supplementary Figure 11**), consistent with the additional sensitivity for detecting transcripts in highly expressed genes.

**Figure 3:**
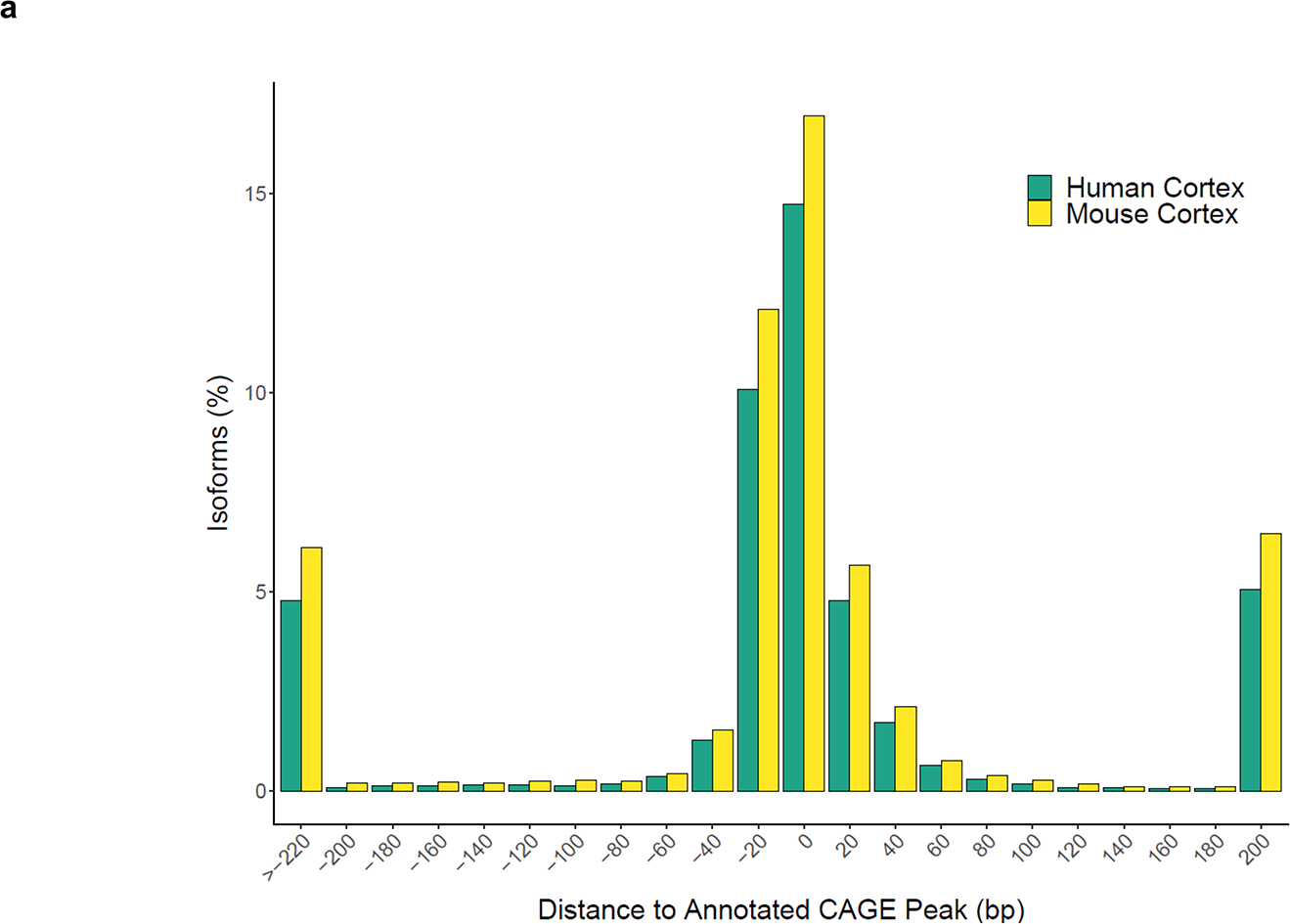
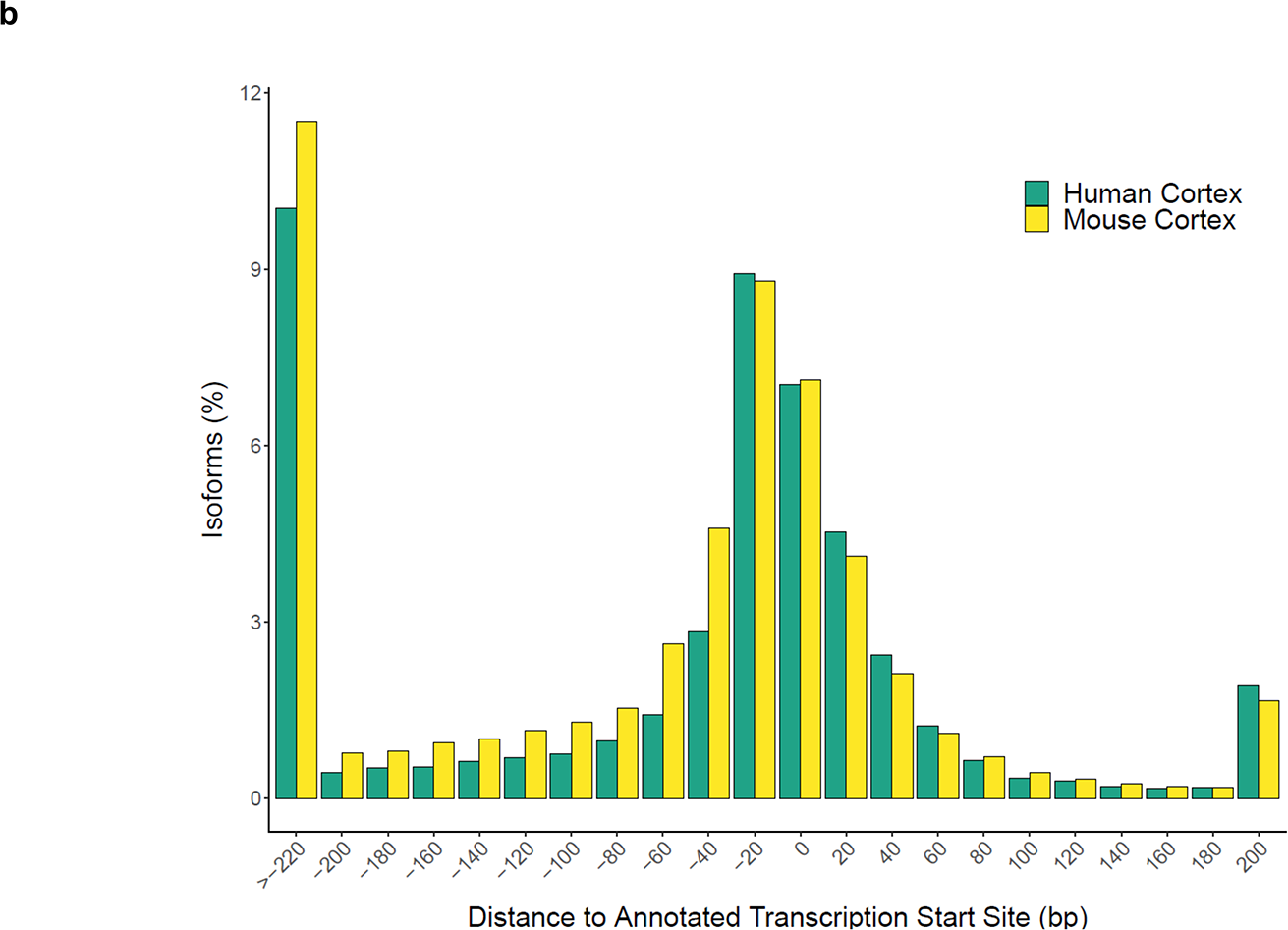
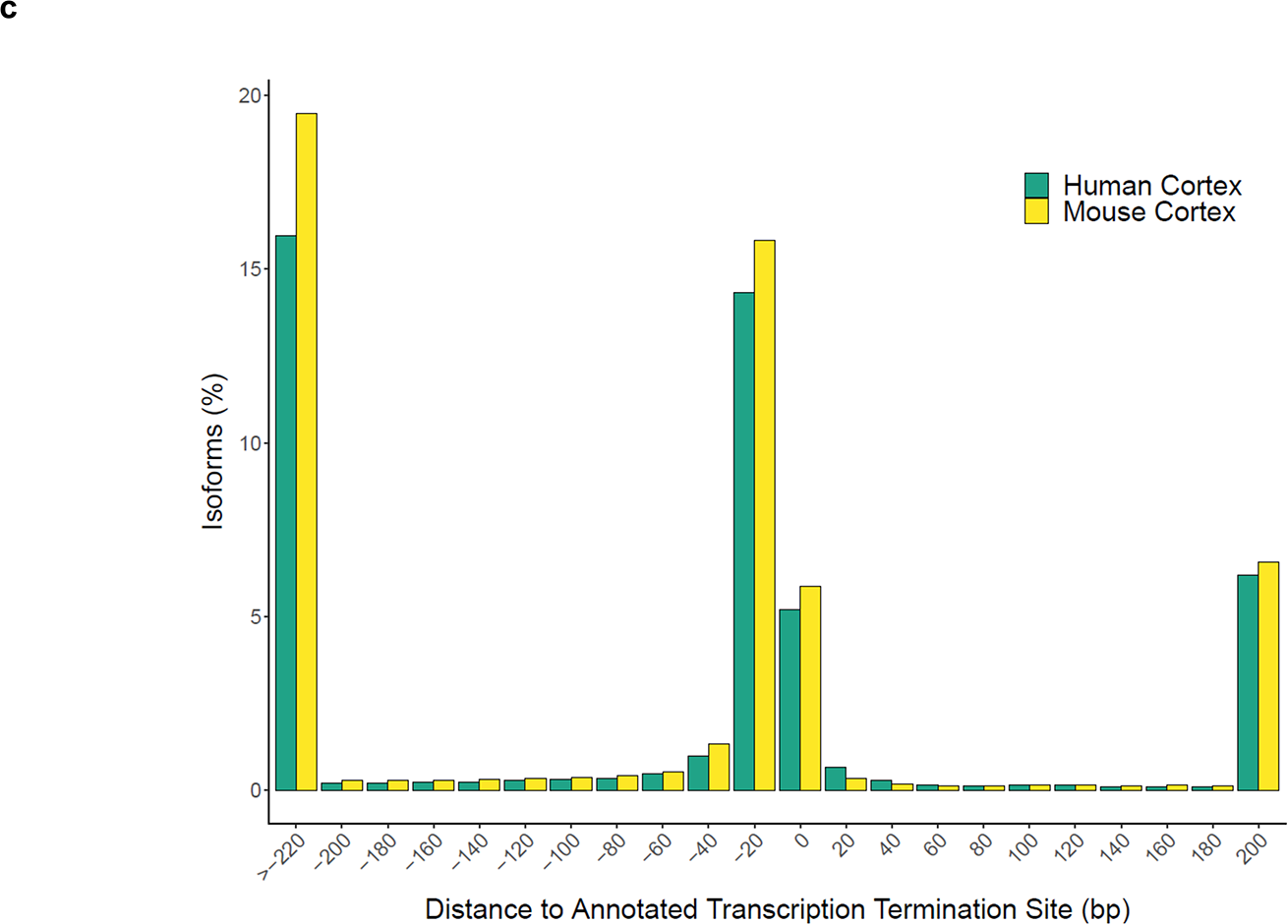
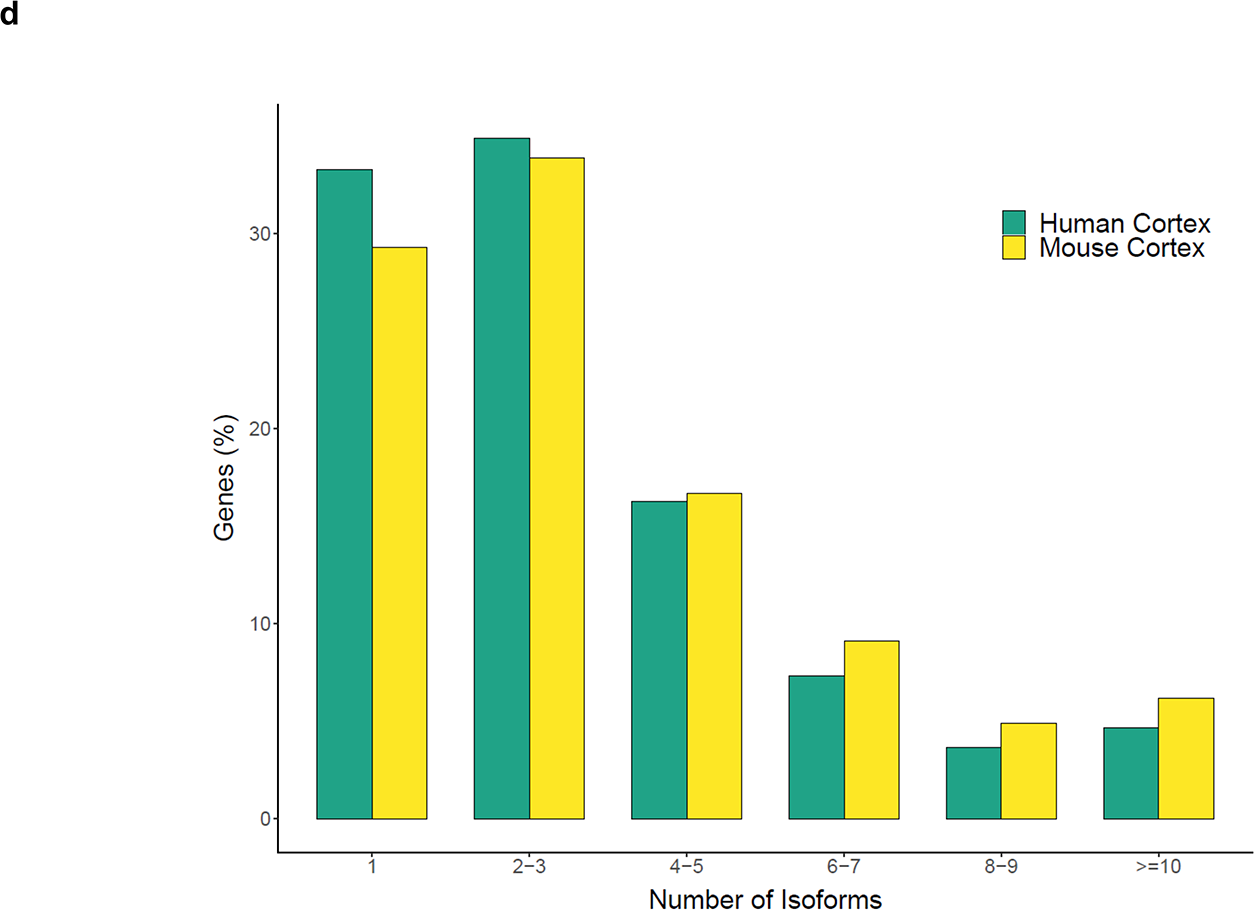
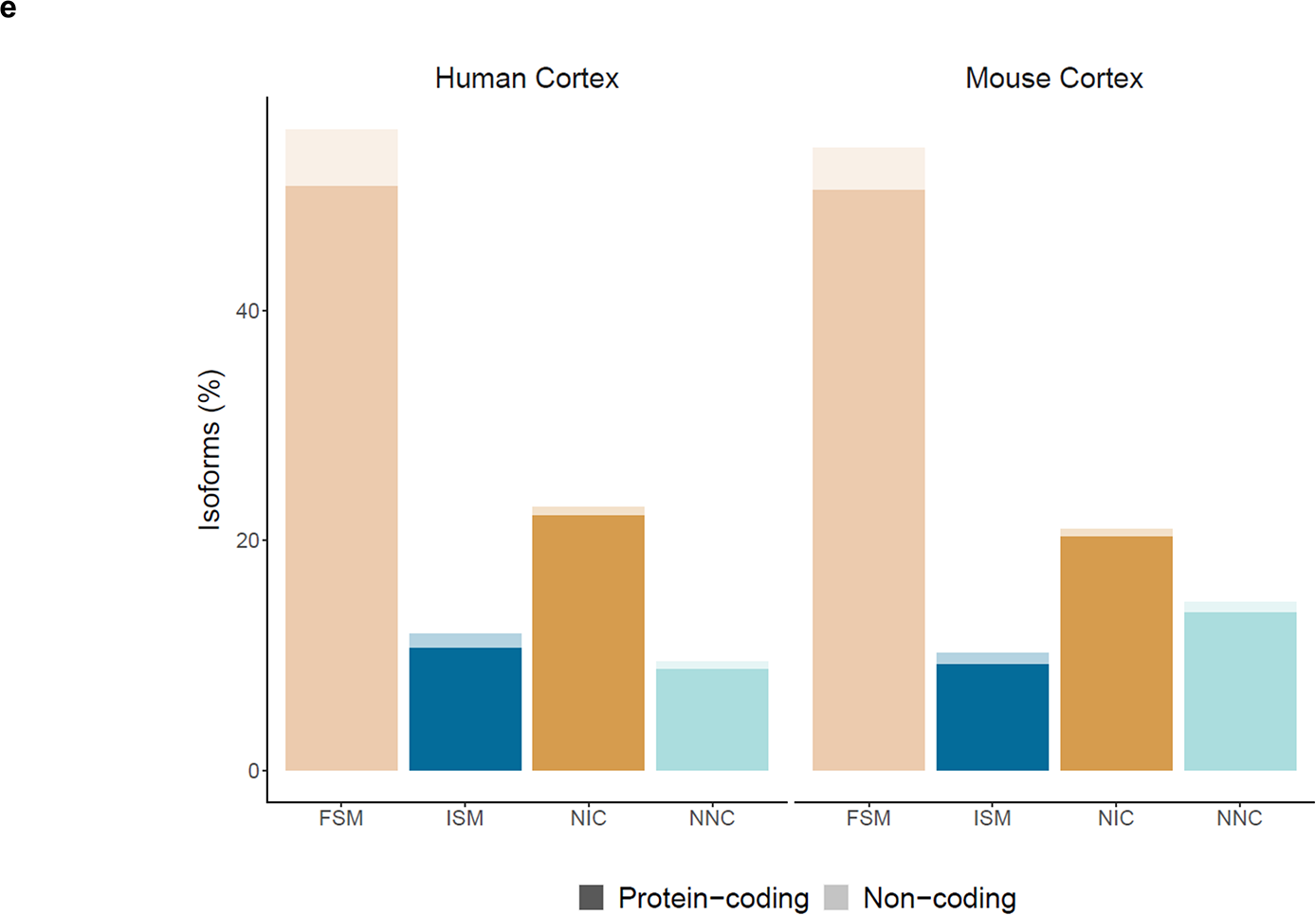
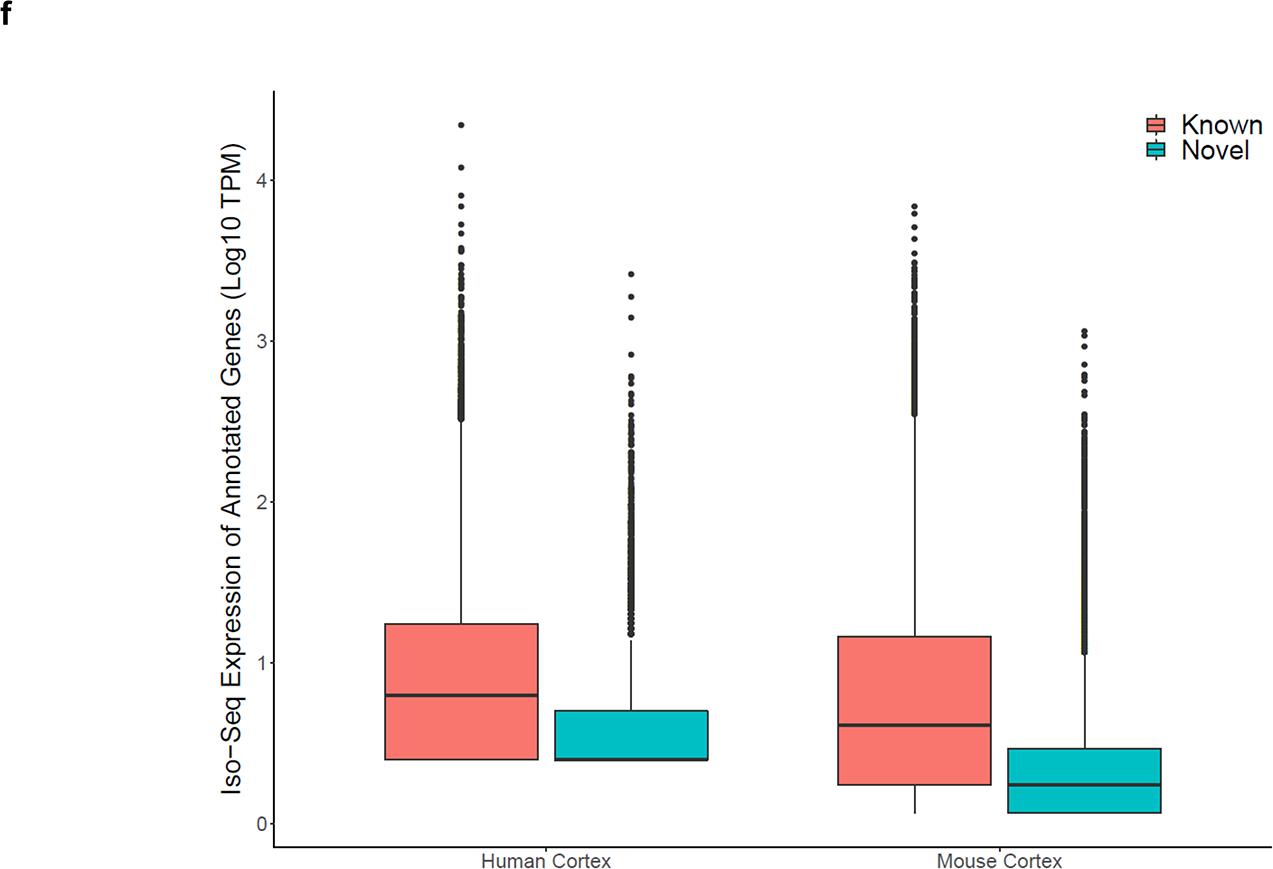
A large proportion of cortical transcripts represent novel transcripts not in existing annotations. **a)** The location of multi-exonic transcripts relative to annotated CAGE peaks in the human and mouse cortex. **b)** The location of multi-exonic transcripts relative to annotated TSSs in the human and mouse cortex. **c)** The location of multi-exonic transcripts relative to annotated TTSs in the human and mouse cortex. **d)** The distribution of isoform numbers identified for each detected gene in the human (n = 7 biologically independent samples) and mouse (n = 8 biologically independent samples) cortex. The distribution of isoform numbers in the adult and fetal cortex can be found in **Supplementary Figure 8. e)** The proportion of multi-exonic transcripts across different *SQANTI2* classification categories (**Figure 1e**). **f)** Iso-Seq transcript expression of known and novel multi-exonic transcripts of genes annotated to reference genome (human: hg38, mouse: mm10). Known transcripts were classified as FSM and ISM, and novel transcripts were classified as NIC, NNC, fusion and antisense. Novel transcripts were characterized by a significantly lower transcript expression than annotated transcripts in both the human cortex (Mann-Whitney-Wilcoxon test, W = 2.59 × 10^8^, P < 2.23 × 10^−308^) and mouse cortex (Mann-Whitney-Wilcoxon test, W = 3.98 × 10^8^, P < 2.23 × 10^−308^). FSM – Full Splice Match, ISM – Incomplete Splice Match, NIC - Novel In Catalogue, NNC - Novel Not in Catalogue, TSS – Transcription Start Site, TTS – Transcription Termination Site.

### Identification of novel transcripts for a large proportion of expressed genes in the cerebral cortex

Amongst full-length transcript reads aligning to annotated genes, the majority of transcripts were characterized by a complete full splice match (FSM) (human cortex: n = 23,629 (55.5%) transcripts; mouse cortex: n = 27,475 (53.9%)) or incomplete splice match (ISM) (human cortex: n = 5.025 (11.8%); mouse cortex: 5,189 (10.2%)) to existing annotations (**Figure 3**, **Supplementary Table 7**). A significant proportion of transcripts, however, represented novel transcripts not present in existing annotation databases. In the human cortex, 13,931 (32.7%) of detected transcripts associated with 5,622 (43.8%) genes were classified as novel (mean size = 2.8kb, s.d = 1.2kb, range = 0.21 – 11.3kb, mean number of exons = 11.1). Similarly, in the mouse cortex, 18,339 (36.0%) of detected transcripts associated with 6,694 (49.8%) genes were classified as novel (mean size = 3.1kb, s.d = 1.5kb, range = 0.16 – 11.7kb, mean number of exons = 12.3). The majority of these novel transcripts contained a combination of known donor and acceptor splice sites and were classified as ‘novel in catalogue’ (NIC) (human cortex: 9,724 (69.8% of all novel transcripts), mouse cortex: 10,649 (58.1% of all novel transcripts), **Supplementary Figure 12**).The remaining novel transcripts were predominantly classified as ‘novel not in catalogue’ (NNC), with at least one novel donor or acceptor site (human cortex: 4,013 (28.9% of all novel transcripts), mouse cortex: 7,414 (40.4% of all novel transcripts)). Amongst genes with novel detected transcripts, there was considerable variation in the number of novel multi-exonic RNA isoforms identified (human cortex: 1 - 30, mouse cortex: 1 - 54, **Supplementary Table 8**); *SEPT4* was found to have the highest number of novel isoforms in human cortex and *Sorbs1* was found to have the highest number of novel isoforms in mouse cortex. Overall, novel transcripts were less abundant than known ones (human cortex: Mann-Whitney-Wilcoxon test W = 2.59 × 10^8^, P < 2.23 × 10^−308^, mouse cortex: Mann-Whitney-Wilcoxon test W = 3.98 × 10^8^, P < 2.23 × 10^−308^, **Figure 3**, **Supplementary Figure 13**), suggesting that they would have been hard to detect using traditional short-read RNA-Seq because of the difficulty in assembling transcripts with limited read coverage^29^. Of note, novel transcripts were both longer (human cortex: W = 1.61 × 10^8^, P = 1.03 × 10^−133^; mouse cortex: W = 2.38 × 10^8^, P = 9.75 × 10^−203^, **Supplementary Figure 14**) and had more exons (human cortex: W = 1.43 × 10^8^, P < 2.23 × 10^−308^; mouse cortex: W = 2.22 × 10^8^, P < 2.23 × 10^−308^, **Supplementary Figure 14**) than already known transcripts. We used the FANTOM5 CAGE^25^ database to show that the majority (human cortex: 10,715 (76.9% of all novel transcripts), mouse cortex: 13,054 (71.2% of all novel transcripts)) of novel transcripts identified using Iso-Seq were within 50bp of an annotated CAGE peak (**Supplementary Figure 15**).

### Validation of novel transcripts identified by Iso-Seq using short-read RNA-seq and nanopore sequencing

To validate the novel multi-exonic transcripts identified in our Iso-Seq datasets, we first used exon-spanning reads obtained from the highly-parallel short-read RNA-Seq data generated for an overlapping subset of samples. Strikingly, these data supported all splice junctions for the majority of novel transcripts (human fetal cortex: 5,618 (89.2%) junctions, mouse cortex: 15,195 (82.9%) junctions), with less than 1% of transcripts having no support from short-read RNA-Seq data (**Supplementary Figure 16**). Next, we interrogated publicly-available Iso-Seq data^30^ from an Alzheimer’s disease (AD) brain sample, processing the data through the same analytical pipeline (see **Methods**). Of the 13,931 novel transcripts identified in our human cortex dataset and mapped to annotated genes, 5,817 (41.76%) were also detected in this single AD dataset. Finally, we used an alternative long-read sequencing method (nanopore sequencing, see **Methods**) to generate additional long-read transcript sequences for a subset of human cerebral cortex (n = 2, 40.7 million reads, 23,609 polished transcripts (mean length = 1.39kb, s.d = 0.97kb, range = 0.085 - 7.47kb) mapping to 9,762 genes). Overall, transcriptional patterns were very similar between the PacBio and ONT datasets (**Supplementary Figure 17**) with a large proportion of novel transcripts of annotated genes from the Iso-Seq dataset also detected in the ONT dataset (human cortex: 7,081 (50.83%) of novel transcripts).

### A subset of cortex-expressed transcripts represent fusion events between neighbouring genes

Transcriptional read-through between two (or more) adjacent genes can produce ‘fusion transcripts’^31^ that represent an important class of mutation in several types of cancer^32^. Although fusion events are thought to be rare^33^, we found that ~0.4% of transcripts included exons from two or more adjacent genes (human cortex: n = 153 fusion transcripts associated with 114 genes (0.89%); mouse cortex: n = 219 fusion transcripts associated with 160 genes (1.19%)). A number of genes were associated with more than one fusion transcript (human cortex: n = 23 (20.2%) genes; mouse cortex: n = 40 (25%) genes), and we identified examples of fusion transcripts encompassing more than two genes - e.g. a fusion transcript incorporating exons from three adjacent pseudogenes in the human cortex *AC138649.4_AC138649.1_PDCD6IPP1* (**Supplementary Figure 18**). The majority of fusion transcripts identified in our Iso-Seq data were supported by short-read RNA-Seq generated on both mouse cortex (n = 212 (96.8%) transcripts) and human cortex (n = 53 (98.1%) transcripts associated with 47 genes (0.49%)). We also confirmed many specific fusion events using our human cortex ONT nanopore sequencing dataset (n = 54 (35.29%) transcripts). Furthermore, several of the fusion genes identified in the human cortex (n = 5 (4.4%) genes) and mouse cortex (n = 7 (4.4%) genes) were predicted as potential ‘conjoined-genes’ in the ConjoinG database^34^. Although the majority of fusion events were specific to the human or mouse cortex datasets, we found evidence of fusion, protein-coding transcripts incorporating exons from *TMEM107* and *VAMP2* in both species (**Figure 4**). Of note, both of these genes are known to be associated with rare neurodevelopmental disorders^35^, and these fusions were supported by RNA-Seq junction-spanning reads in both species and ONT sequencing reads from the human cortex.

**Figure 4:**
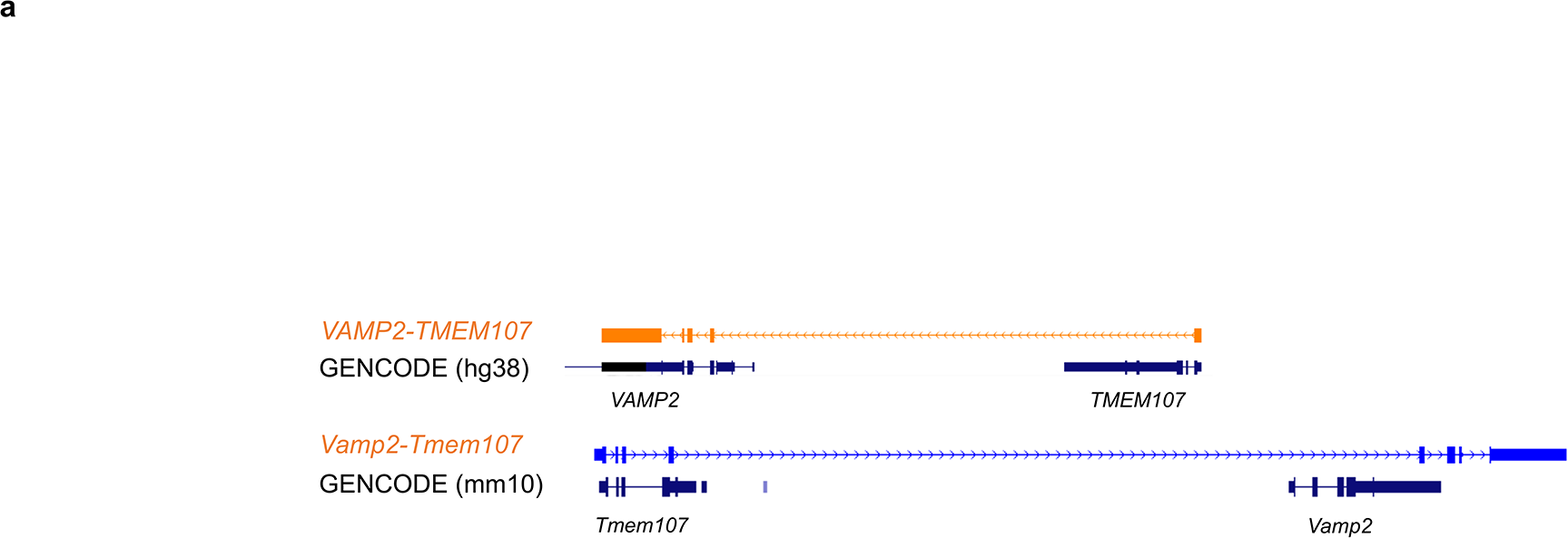
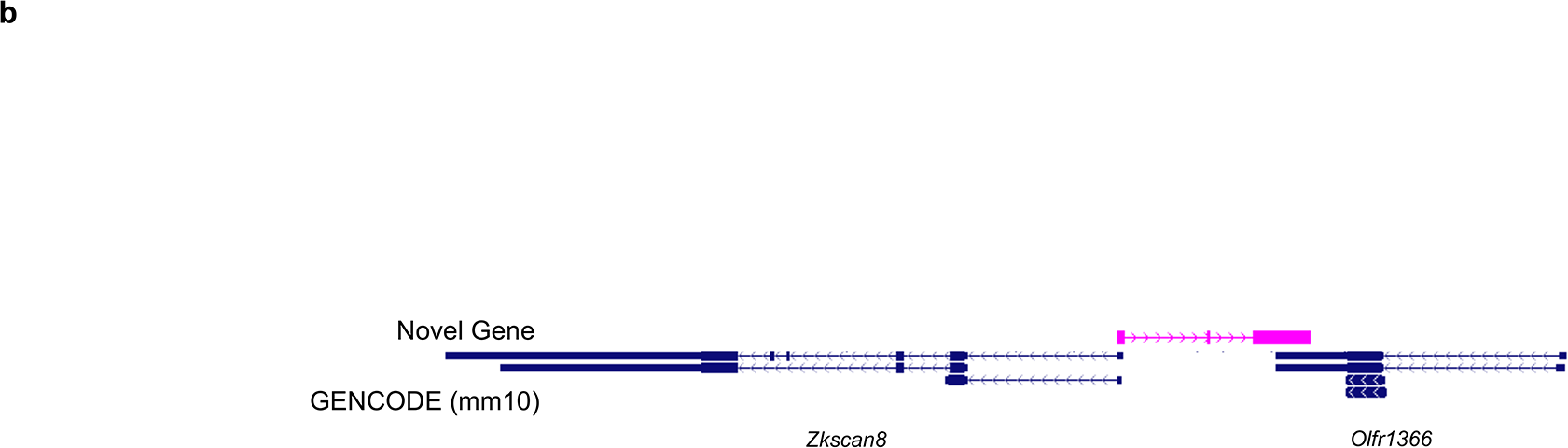
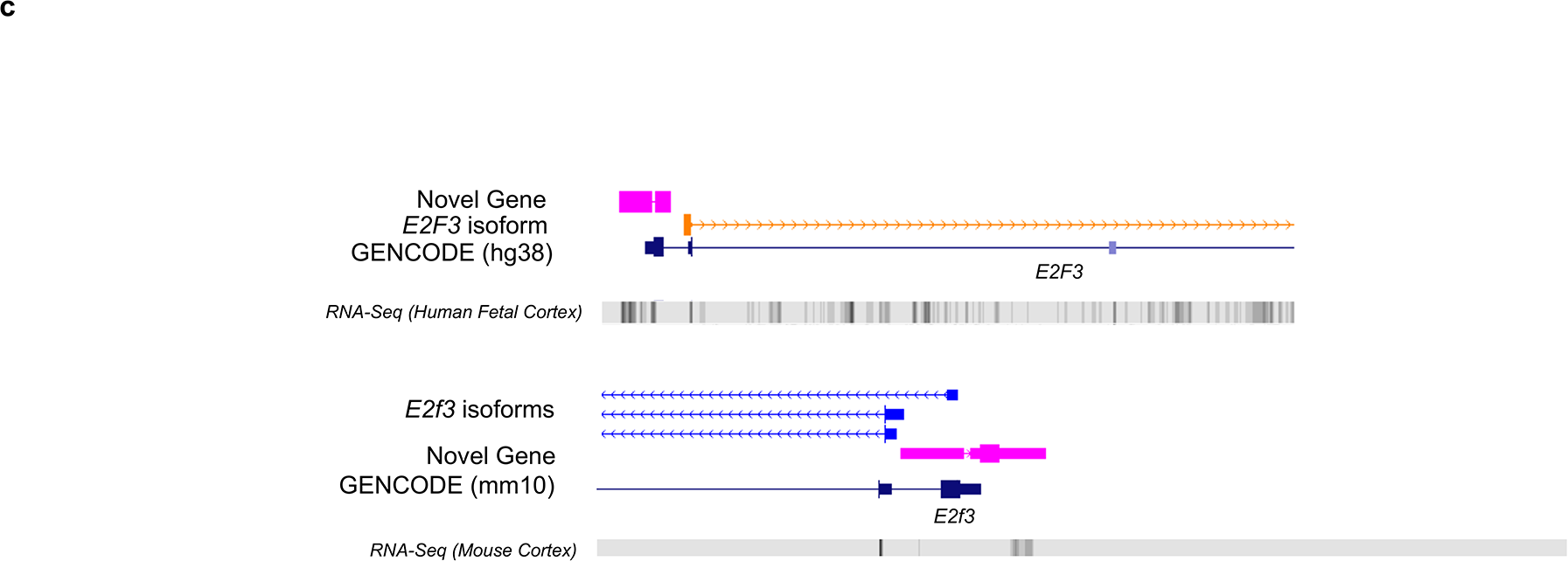
Examples of fusion transcripts and novel genes were identified in the human and mouse cortex. **a)** A common fusion transcript incorporating exons from *TMEM107* and *VAMP2* in both human (n = 7 biologically independent samples) and mouse (n = 8 biologically independent samples) cortex. **b)** An example of a novel gene that shared exons across *Zkscan8* and *Olfr1366*, and is antisense to both genes, in the mouse cortex. Additional examples of fusion transcripts are shown in **Supplementary Figure 20. c)** A common novel gene was identified in both human and mouse cortex, antisense and upstream to *E2F3*. Isoforms are coloured based on *SQANTI2* classification categories (blue = FSM, cyan = ISM, red = NIC, orange = NNC, purple = antisense). FSM – Full Splice Match, ISM – Incomplete Splice Match, NIC – Novel In Catalogue, NNC – Novel Not in Catalogue.

### Identification of novel cortex-expressed genes using long-read sequencing

Although the vast majority of transcripts identified in both the human and mouse cortex were assigned to annotated genes (human cortex: 99.9% of total transcripts; mouse cortex: 99.7% of total transcripts), a small number did not and potentially represented novel genes (human: n = 60 novel transcripts mapping to 49 novel genes; mouse: n = 156 novel transcripts mapping to 131 novel genes) (**Supplementary Table 9**). These novel genes were all multiexonic (human: mean length = 2.0kb, s.d = 0.9kb, range = 0.4 - 4.9kb, mean number of exons = 2.9; mouse: mean length = 1.7kb, s.d = 1.1kb, range = 0.3 - 6.9kb, mean number of exons = 2.5), with over half the identified transcripts from these genes predicted to be non-coding (human: n = 40 (66.7%) novel-gene transcripts; mouse: n = 86 (55.1%) novel-gene transcripts), and shorter than annotated genes (human cortex: W = 1.6 × 10^6^, P = 1.5 × 10^−4^, mouse cortex: W = 5.7 × 10^6^, P = 2.8 × 10^−25^, **Supplementary Figure 19**). The overall expression of these novel genes was lower than that of annotated genes (human cortex: W = 5.1 × 10^5^, P = 9.8 × 10^−18^; mouse cortex: W = 1.5 × 10^6^, P = 7.7 × 10^−59^), although a number were characterized by relatively high expression (**Supplementary Figure 19**). Although the majority of these novel genes did not show high homology with other genomic regions, BLAST analysis identified 18 (21.7%) homologous (greater than 500bp, more than 90% identity) novel-gene transcripts in human cortex and 26 (16.0%) homologous novel-gene transcripts in mouse cortex (**Supplementary Table 10**). Of the 60 novel-gene transcripts identified in the human cerebral cortex, 28 (46.7%) were also identified as novel by the GTEx consortium (CHESS v2.2 annotation)^36^. Furthermore, our matched short-read RNA-Seq data fully supported novel genes identified in a subset of human cerebral cortex samples (human fetal cortex: n = 22 novel transcripts mapping to 20 novel genes). Ten (16.67%) of the putative novel-gene transcripts identified in the human cortex were also supported by transcripts present in a publicly-available Iso-Seq dataset from an AD brain sample^30^. Finally, further evidence of transcription from a large proportion of these novel-gene transcripts (human cortex: 28 (46.67%)) was provided by our ONT nanopore sequencing datasets generated on an overlapping set of human cortical samples. We used the FANTOM5 CAGE dataset to show that about a quarter of the novel-gene transcripts (human cortex: n = 13 (21.7%), mouse cortex: n = 39 (25.0%)) were located within 50bp of a CAGE peak (**Supplementary Table 9**). Of note, there was an enrichment of antisense transcripts amongst those mapping to novel genes (human cortex: n = 34 transcripts (56.7%) mapping to 27 novel genes; mouse cortex: n = 78 transcripts (50%) mapping to 62 novel genes) (**Supplementary Table 9**). The majority of the antisense novel genes were found within an annotated gene (human cortex: n = 21 (77.8% of antisense novel genes), mouse cortex: n = 58 (93.5% of antisense novel genes)), with a relatively large proportion of these sharing exonic regions (exon-exon overlap, human cortex: n = 8 (38%), mouse cortex: n = 42 (72.4%) reflecting sense-antisense (SAS) pairs^37^. Furthermore, there were striking examples of antisense novel genes that shared exons from two genes in the mouse cortex (**Figure 4, Supplementary Figure 20**). With the majority of novel genes specific either to human or mouse cortex, we identified one common protein-coding novel gene that overlapped, upstream and antisense to *E2F3* (identity = 83.1%, E-value = 0, alignment length: 1,926bp, mismatches: 205bp, **Supplementary Table 11**). This common *E2F3*-associated novel gene was strongly conserved between both species with a similar transcript structure of two exons, and was supported by RNA-Seq in both human and mouse cortex (**Figure 4**).

### Many transcripts map to long non-coding RNA genes and a subset of these were found to contain open reading frames

Although the majority of transcripts mapping to annotated genes were classified as protein-coding by the presence of an ORF in *SQANTI2* (human cortex: n = 39,352 (92.4%) transcripts associated with 11,959 genes; mouse cortex: n = 47,757 (93.6%) transcripts associated with 12,748 genes), a relatively large number of transcripts mapped to genes annotated as encoding lncRNA (human cortex: n = 1,545 transcripts associated with 829 genes; mouse cortex: n = 1,041 transcripts associated with 587 genes). lncRNA transcripts were found to be longer than transcripts not defined by reference genome as lncRNA (non-lncRNA) (human cortex: mean length of lncRNA transcripts = 2.2kb, s.d = 1.1kb, range = 0.082 - 7.8kb, mean length of non lncRNA transcripts = 2.6kb, s.d = 1.3kb, range = 0.09 - 11.8kb), W = 3.84 × 10^7^, P = 1.17 × 10^−45^; mouse cortex: mean length of lncRNA transcripts = 2.1kb, s.d =1.3kb, range = 0.1 - 8.5kb, mean length of non lncRNA transcripts = 2.9 kb, s.d = 1.6kb, range = 0.08 - 15.9kb), Mann-Whitney-Wilcoxon test W = 3.36 × 10^7^, P = 2.83 × 10^−58^, Supplementary Figure 21), despite containing fewer exons (human cortex: W = 5.57 × 10^7^, P < 2.23 × 10^−308^; mouse cortex: W = 4.54 × 10^7^, P < 2.23 × 10^−308^, Supplementary Figure 21) and being enriched for mono-exonic molecules^38^ (human cortex: n = 533 (34.5%); mouse cortex: n = 282 (27.1%)) compared to non-lncRNA transcripts (human cortex: n = 1,021 (2.5%), mouse cortex: n = 1,325 (2.7%)). They were also characterised by lower transcript expression than non-lncRNA transcripts^39^ (human: W = 3.70 × 10^7^, P = 1.12 × 10^−30^; mouse: W = 3.28 × 10^7^, P = 4.13 × 10^−49^, Supplementary Figure 22), with fewer RNA isoforms identified per lncRNA gene (human cortex: mean n = 1.86, range = 1 - 87, mouse cortex: mean n = 1.8, range = 1 - 53) compared to coding genes (human cortex: mean n = 3.4, range = 1 - 40, mouse cortex: mean n = 3.9, range = 1 - 83) (human cortex: W = 7.11 × 10^6^, P = 1.47 × 10^−99^, mouse cortex: W = 5.71 × 10^6^, P = 2.09 × 10^−101^). Importantly, over a third of these annotated lncRNA transcripts actually contained a putative ORF (human cortex: n = 601 (38.9%); mouse cortex: n = 413 (39.7%)) supporting recent observations that lncRNA have potential protein coding capacity^40^; these ORFs were shorter (human cortex: mean length = 132.7bp, s.d = 80.4bp; mouse cortex: mean length = 133.6bp, s.d = 110.8bp) than those in non-lncRNA transcripts (human cortex: mean length = 434.4bp, s.d = 291.7bp, W = 2.11 × 10^7^, P = 1.01 × 10^−254^; mouse cortex: mean length = 502.1bp, s.d = 363.9bp, W = 1.79 × 10^7^, P = 9.50 × 10^−186^, Supplementary Figure 23).

### Alternative splicing events make a major contribution to RNA isoform diversity in the cortex

Alternative splicing (AS), the process by which different combinations of splice sites within a messenger RNA precursor are selected to produce variably spliced mRNAs, is the primary mechanism underlying transcript diversity in eukaryotes^41^ and a major source of transcriptional diversity in the central nervous system^10^. Numerous types of AS have been described (**Figure 5**) and we used a combination of the *SUPPA2*^42^ package and custom analysis scripts (see **Methods**) to identify transcripts associated with i) exon skipping (SE), ii) mutually exclusive exon use (MX), iii) alternative first (AF) and last (AL) exons, iv) alternative 3’ (A3’) and 5’ (A5’) splice sites, and v) intron retention (IR) in our cortical Iso-Seq datasets. The overall frequency of these specific AS events was similar in human and mouse cortex, with AF and AL being the most prevalent AS events in both species (human cortex: AF = 11,333 (33.0%) events associated with 5,307 (44.8%) genes; AL: 7,262 (21.5%) events associated with 4,272 (35.2%) genes; mouse cortex: AF = 14,111 (35.7%) events associated with 6,089 (46.9%) genes); AL = 9,883 (25.0%) events associated with 5,054 (38.9%) genes (**Figure 5, Supplementary Figure 24, Supplementary Table 12**). 5,473 unique genes produced AS transcripts in both the human (70.4% of AS genes) and mouse cortex (63.4% of AS genes) (**Figure 5, Supplementary Figure 25**). For the majority of AS genes, only one or two AS events were observed (human cortex: n = 4,858 (62.5% of AS genes), mouse cortex, n = 9,774 (83.2% of AS genes)), suggesting that AS events were often mutually independent (**Figure 5, Supplementary Figure 26**). In contrast, however, a small number of genes were characterized by all AS events analysed (human cortex: n = 7(0.09%)). Although skipped exons are known to be the most common AS events in both human and mouse^43^, our data conversely suggests that splice variants from a single gene are predominantly generated through alternative first and alternative last exons (**Supplementary Figure 27**).

**Figure 5:**
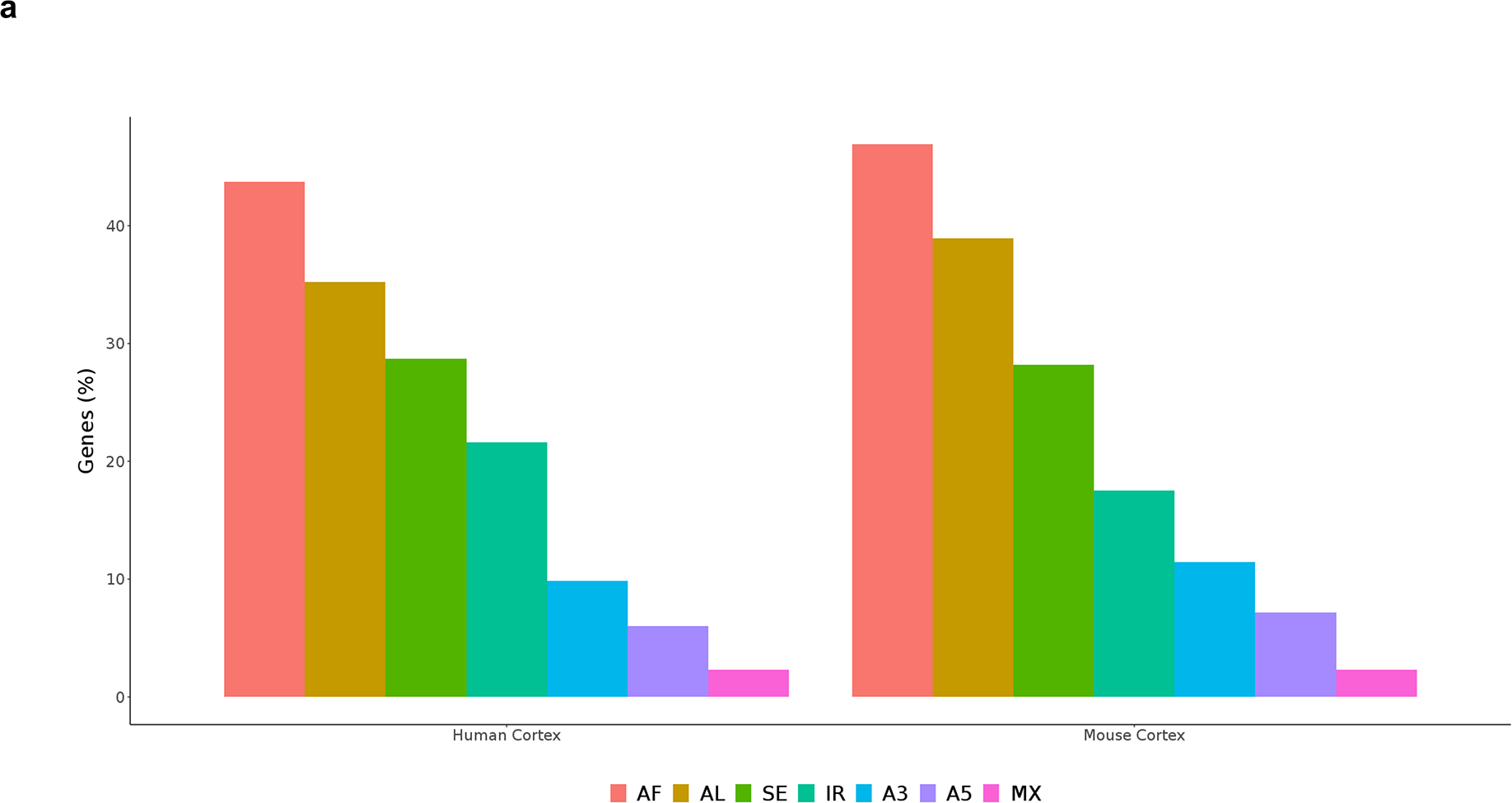
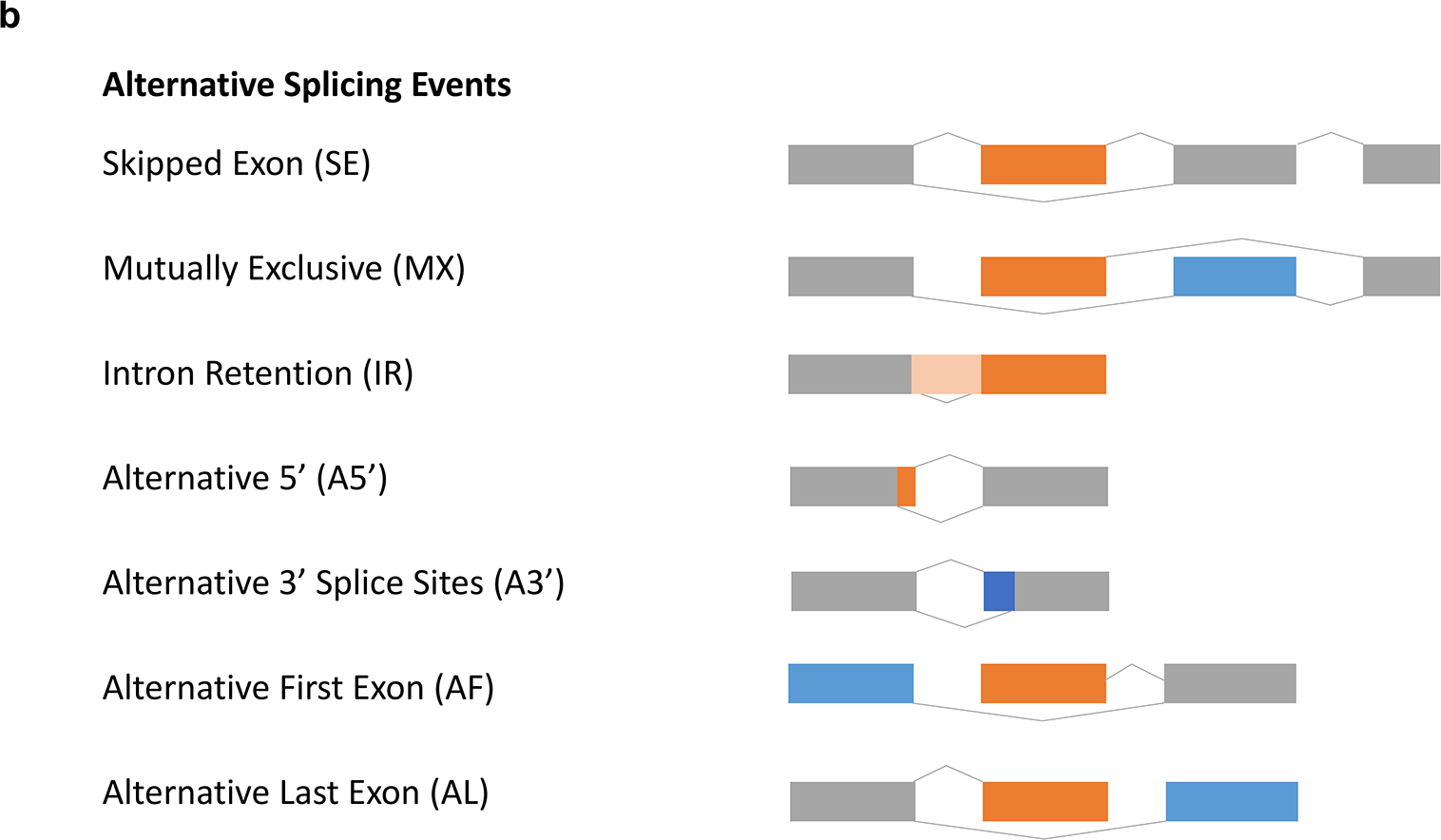
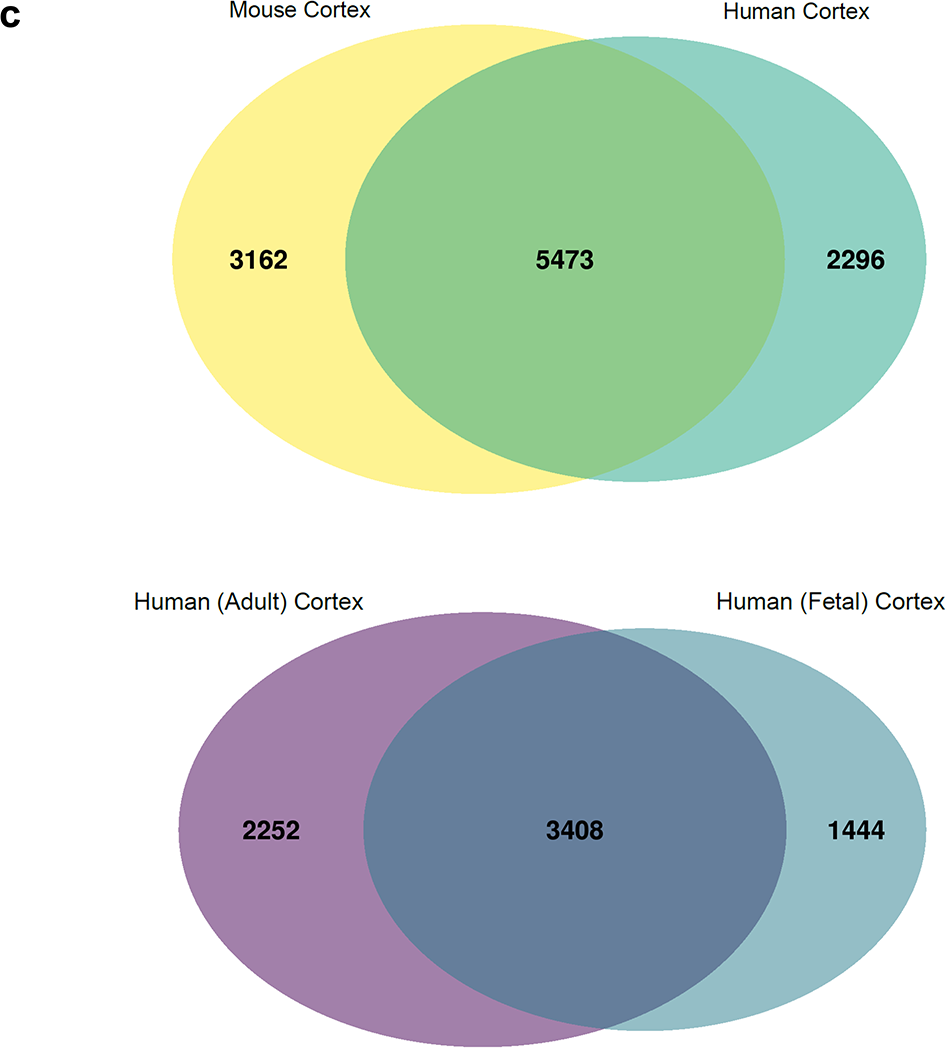
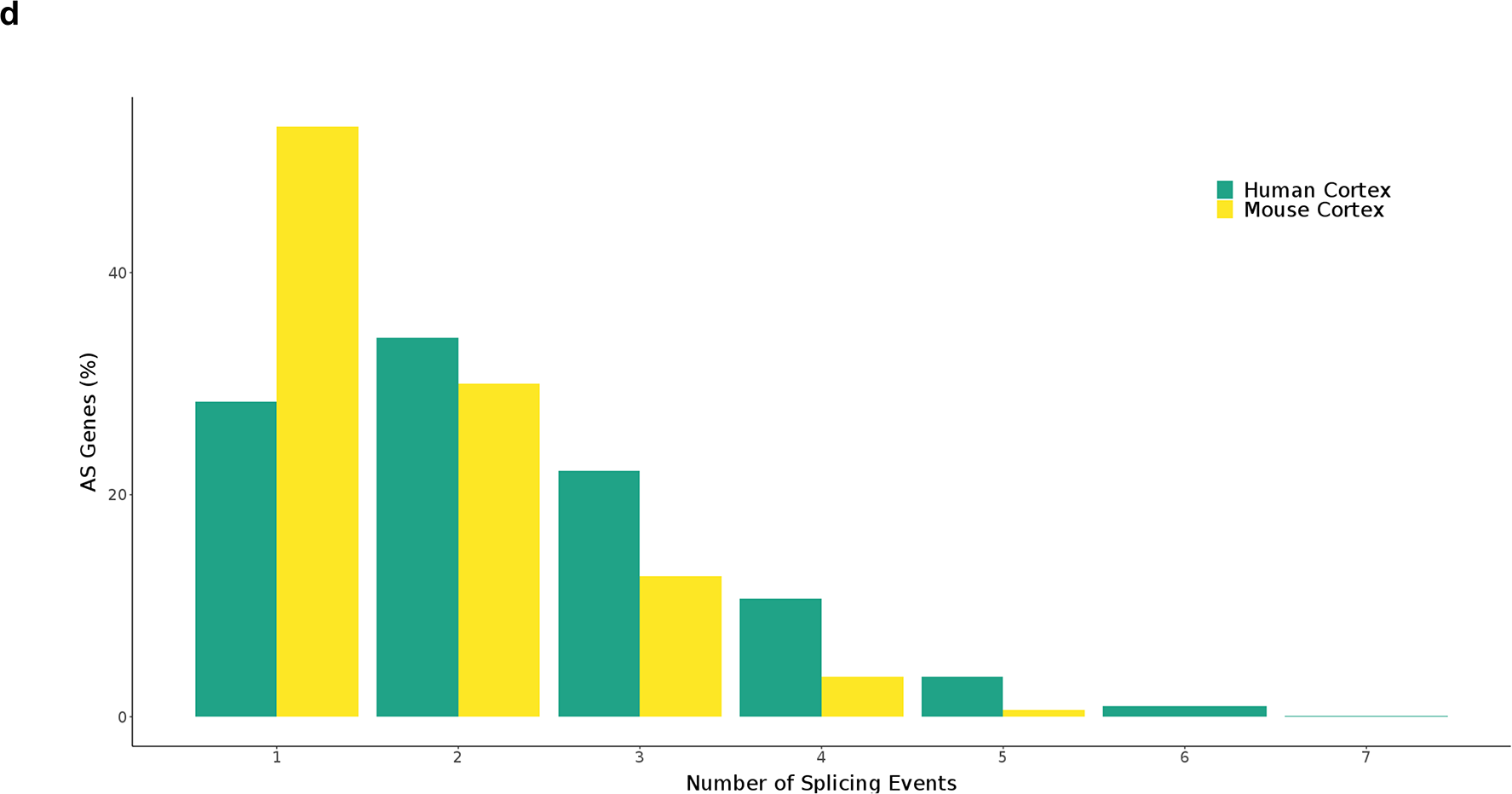
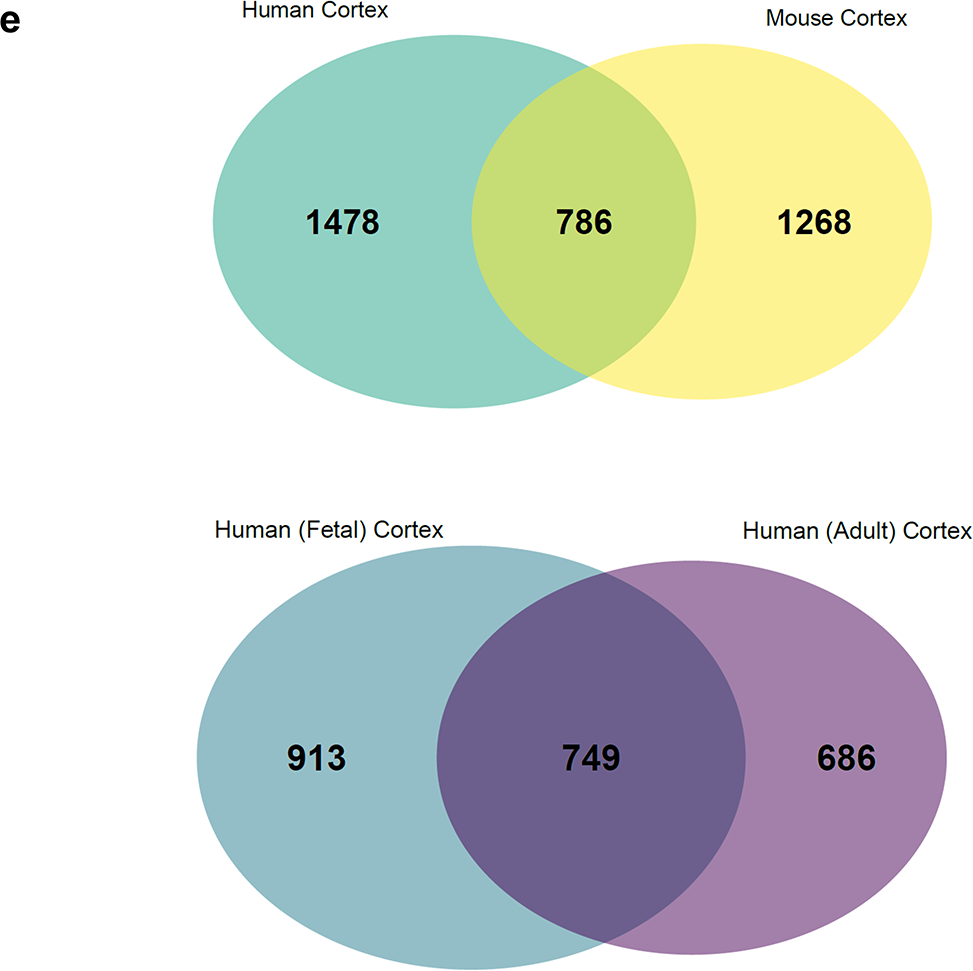
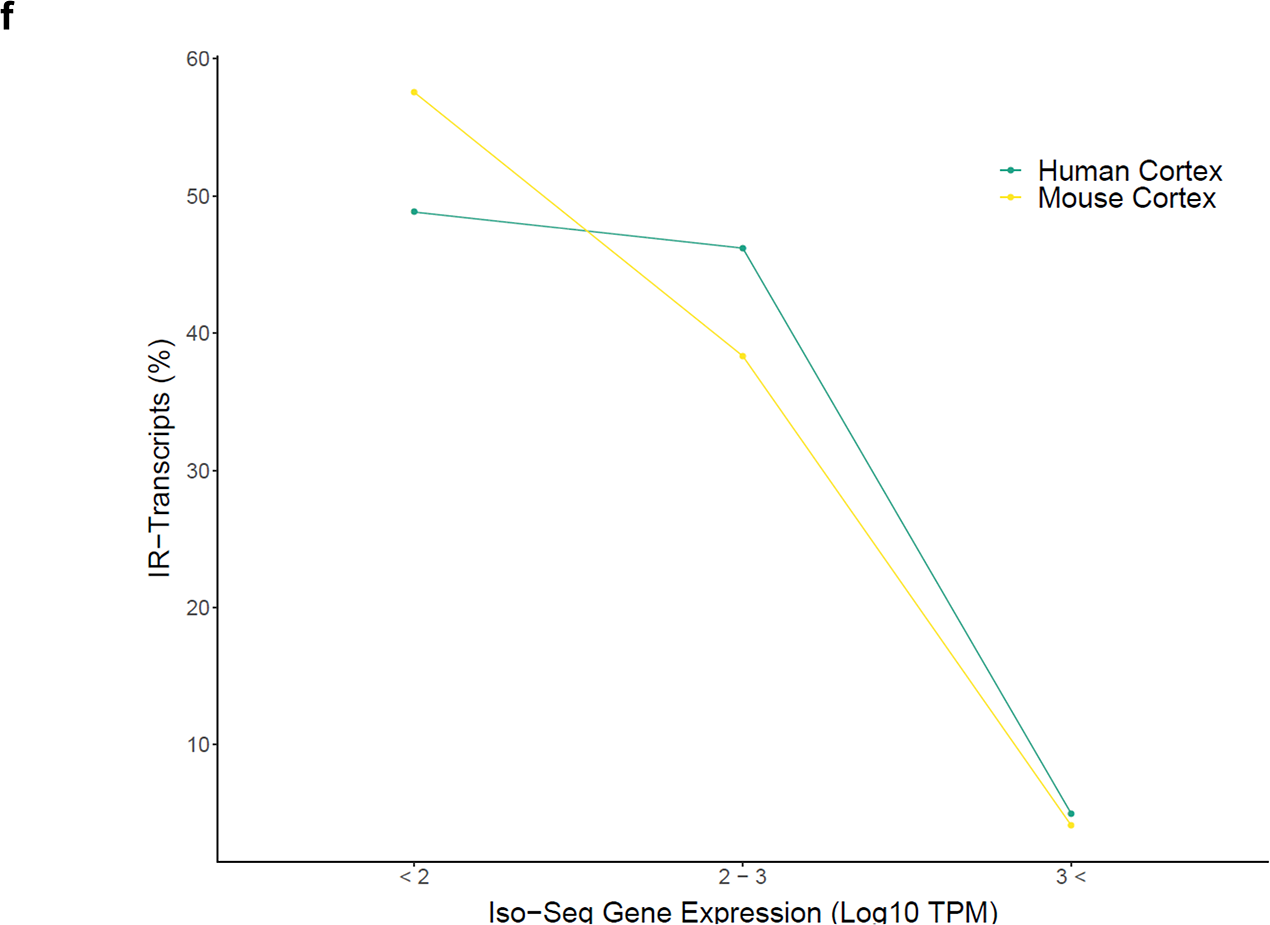
Alternative splicing events make a major contribution to RNA isoform diversity in the cortex. **a)** The majority of cortex-expressed genes were characterised by multiple transcripts generated by **b)** various alternative splicing (AS) events, with alternative first exon (AF) and alternative last exon (AL) usage being the most common. Further details can be found in **Supplementary Table 12**. **c)** A large overlap of genes influenced by AS was observed in both human and mouse cortex, and human adult and human fetal cortex, with **d)** a large proportion of these AS genes characterised by more than one type of splicing event. **e)** This can be seen in the large overlap in genes characterized by Intron retention (IR) in the human and mouse cortex, and between fetal and adult human cortex. See **Supplementary Figure 25** for the overlap of genes across the different splicing events across the cortical Iso-Seq datasets. **f)** IR is present at a higher rate amongst transcripts from lowly-expressed genes in both the human and mouse cortex. AF – Alternative First Exon, AL – Alternative Last Exon, A5’ – Alternative 5’ prime, A3’ – Alternative 3’ prime, IR – Intron Retention, MX – Mutually Exclusive, SE – Skipped Exon.

### Intron retention is a relatively common form of alternative splicing in the cortex that is associated with reduced expression and nonsense-mediated mRNA decay (NMD)

Intron retention (IR), the process by which specific introns remain unspliced in polyadenylated transcripts, is the least understood AS mechanism in vertebrates^44^ but is hypothesized to be a particularly important mechanism of transcriptional control in the brain^45^. We found evidence for IR in a relatively large proportion of genes (IR-genes) in both the human cortex (n = 5,752 IR-transcripts associated with 2,625 (20.4%) detected genes) and mouse cortex (n = 4,216 IR-transcripts associated with 2,279 (16.8%) genes) (**Supplementary Table 13**), with IR-genes themselves enriched for biological processes related to mRNA splicing (human cortex: odds ratio = 2.51, P = 8.43 × 10^−17^; mouse cortex: odds ratio = 2.45, P = 1.41 × 10^−13^, **Supplementary Table 4**). The majority of IR-transcripts were supported by matched short-read RNA-Seq data performed on the human fetal cortex samples (n = 2,956 (96.8%) IR-transcripts) and mouse cortex samples (n = 3,991 (94.7%) IR-transcripts). The majority of IR-genes were found to express more than one IR-transcript (human cortex: n = 1,411 (75%) IR-genes; mouse cortex: n = 1,385 (67%) IR-genes), with *MEG3* having the largest number of IR-transcripts (62 isoforms (71.3% of *MEG3* isoforms), **Supplementary Figure 8**) in human cortex and *Dlgap4* having the largest number of IR-transcripts (25 isoforms (78.1% of *Dlgap4* isoforms)) in mouse cortex. A small number of genes were found to *only* express transcripts characterized by IR (**Supplementary Table 14**) (human cortex: n = 194 (7.4% of genes with IR-transcripts, 1.5% of total detected genes), mouse cortex: n = 125 (5.5% of genes with IR-transcripts, 0.9% of total detected genes). Overall, there was considerable overlap in the list of IR-genes between human and mouse cortex (**Figure 5**), with 786 homologous genes showing evidence of IR in both the human (62.0% of IR-genes) and mouse (53.2% of IR-genes) cortex. Importantly, a larger proportion of lowly expressed genes showed evidence for IR than highly expressed genes in both human (< 2.5 Log_10_ TPM, n = 2,332 (88.8%) genes; > 2.5 Log_10_ TPM, n = 293 (11.2%) genes) and mouse (< 2.5 Log_10_ TPM, n = 2,030 (89.1%) genes; > 2.5 Log_10_ TPM, n = 249 (10.9%) genes, **Figure 5**) cortex, corroborating previous analyses suggesting that IR is associated with reduced transcript abundance^46^. Nonsense-mediated mRNA decay (NMD) acts to reduce transcriptional errors by degrading transcripts containing premature stop codons^47^ and is one mechanism by which IR can influence gene expression^48^. Overall, about a tenth of transcripts mapping to annotated genes (human cortex: n = 5,062 (11.9%) transcripts associated with 2,420 (18.9%) of genes), mouse cortex: n = 4,944 (9.7%) transcripts associated with 2,264 (16.8%) of genes) were predicted to undergo NMD (NMD-transcripts), characterised by the presence of an ORF and a coding sequence (CDS) end motif before the last junction. These NMD-transcripts were found to be more lowly expressed than non-NMD-transcripts (human cortex: mean expression of NMD-transcripts = 10.9TPM, s.d = 48.6 TPM, mean expression of non-NMD-transcripts = 24.5 TPM, s.d = 179.1 TPM, W = 6.54 × 10^7^, P = 3.19 × 10^−160^; mouse cortex: mean expression of NMD-transcripts = 11.2 TPM, s.d = 65.1 TPM, mean expression of non-NMD-transcripts = 19.5 TPM, s.d = 101.4TPM, W = 8.25 × 10^7^, P = 3.44 × 10^−126^, **Supplementary Figure 28**). NMD was found to be particularly enriched amongst IR-transcripts (human cortex: n = 2,122 (38.7%) IR-transcripts associated with 1,150 (0.09%) genes; mouse cortex: n = 1,374 (34.1%) IR-transcripts associated with 881 (0.07%) genes, **Supplementary Figure 29**) and transcripts with both IR and predicted NMD were particularly lowly expressed (human cortex: W = 6.00 × 10^6^, P = 4.43 × 10^−14^; mouse cortex: W = 3.40 × 10^6^, P = 1.21 × 10^−35^, **Supplementary Figure 28**). Only a small number of genes were associated with transcripts where IR and NMD were mutually exclusive (human cortex: n = 168 (0.01%) genes; mouse cortex: n = 186 (0.01%) genes, **Supplementary Figure 29**), providing additional support for the hypothesized relationship between these two transcriptional control mechanisms^49^.

### Although global patterns of RNA isoform diversity are generally similar between human and mouse cortex, there are some notable exceptions

Although previous studies have highlighted evidence of major splicing diversity between human and mouse^50^, we found that amongst genes present in both human and mouse cortex Iso-Seq datasets (n = 10,070 genes, **Supplementary Figure 4**), the number of RNA isoforms detected for each gene was significantly correlated between species (corr = 0.52, P < 2.23 × 10^−308^). There was a stronger relationship amongst highly-expressed genes (>2.5 Log_10_ TPM in both species, corr = 0.73, P = 1.28 × 10^−48^, **Supplementary Figure 30**), presumably reflecting the fact that these genes were sequenced closer to saturation in our datasets. Despite the overall stability in cortical RNA isoform diversity between human and mouse, there were some notable exceptions for specific genes (**Supplementary Table 6**). The largest absolute difference in numbers of multi-exonic isoforms detected between human and mouse cortex was observed in the genes encoding *SORBS1* (human cortex: n = 5 isoforms; mouse cortex: n = 55 isoforms, **Supplementary Figure 31**) and *DLGAP1* (human cortex = 12 isoforms; mouse cortex = 54 isoforms, **Supplementary Figure 32**). The gene encoding *NDUFS2* had the highest relative number of isoforms detected in human cortex (n = 18 isoforms) compared to mouse cortex (1 isoform, **Supplementary Figure 33**), whereas the gene encoding *TMEM191C* had the highest relative number of isoforms detected in mouse cortex (n = 28 isoforms) compared to human cortex (1 isoform, **Supplementary Figure 34**). We used the matched RNA-Seq data to show that the majority of splice junctions of multi-exonic isoforms of annotated genes in both the human (*GT/AG:* 99.1%, *GC/AG*: 0.79%, *AT/AC*: 0.07%) and mouse cortex were classified as canonical (*GT/AG*: 99.0%, GC/AG: 0.84%, *AT/AC*: 0.07%, **Supplementary Figure 35**). Of the 31 non-canonical junctions supported by RNA-Seq reads, nine were shared between human and mouse cortex, with greater splice junction diversity observed in the mouse cortex (**Supplementary Figure 35**).

### Developmental changes in cortical RNA isoform abundance

Our human cortex Iso-Seq dataset was generated using samples derived from both fetal and adult donors, enabling us to identify developmental variation in transcript diversity. Overall, we detected 23,191 transcripts mapping to 9,647 annotated genes in the fetal cortex (mean length = 2.8kb, s.d = 1.3kb, range = 0.1 - 11.8kb) and 27,842 transcripts mapping to 10,949 annotated genes in the adult cortex (mean length = 2.9kb, s.d = 1.2kb, range = 0.08 - 9.5kb), with a high degree of overlap in detected genes (n = 8,078 (83.7% of fetal annotated genes, 73.7% of adult annotated genes)) between datasets (**Supplementary Figure 4**). Using the Human Gene Atlas database^19^, we found that the most abundant genes (top 500, ranked by TPM) in the fetal cortex dataset were most significantly enriched for ‘fetal brain’ (odds ratio = 6.15, P = 3.32 × 10^−25^) and those in the adult cortex were most significantly enriched for ‘prefrontal cortex’ genes (odds ratio = 6.37, P = 7.11 × 10^−43^, **Supplementary 4**). General patterns of RNA isoform diversity were similar between fetal and adult human cortex. In both, the majority of genes (fetal cortex: n = 5,409 (56.0%); adult cortex: 6,386 (58.2%)) were characterized by more than one isoform (**Supplementary Figure 36**) and a strong correlation was observed between the number of RNA isoforms detected in fetal and in adult human cortex datasets (corr = 0.56, P < 2.23 × 10^−308^), which was even stronger amongst highly-expressed genes (>2.5 Log_10_ TPM in both fetal and adult cortex, corr = 0.72, P = 1.26 × 10^−41^, **Supplementary Figure 30**). Despite these similarities, there were some notable exceptions with certain genes characterized by significantly more isoforms in either the human fetal or human adult cortex; the gene encoding *BCAS1* had the highest relative number of isoforms detected in adult cortex compared to fetal cortex (19 vs 1 isoforms, 609 vs 7.7 TPM), whereas the gene encoding *MAGED4B* having the highest relative number of isoforms in fetal cortex compared to adult cortex (13 vs 1 isoforms, 1.9 vs 6.5 TPM) (**Supplementary Table 15**). The largest absolute difference in isoform numbers detected between human fetal and human adult cortex was for the genes encoding *MEG3* (adult cortex: n = 72 isoforms; fetal cortex: n = 34 isoforms), *SEPT4* (adult cortex: n = 33 isoforms; fetal cortex, n = 3 isoforms) and *MBP* (adult cortex: n = 32 isoforms; fetal cortex: n = 2 isoforms) (**Supplementary Figure 37**). A similar proportion of transcript reads represented novel (previously unannotated) transcripts of genes in both the fetal (n = 6,244 (26.9%) transcripts associated with 3,219 (33.4%) annotated genes) and adult cortex (n = 7,391 (26.6%) associated with 3,682 (33.6%) annotated genes). 1,768 common genes were characterized by novel transcripts in both the fetal cortex (54.9% of genes with novel transcripts) and adult (48.0% of genes with novel transcripts) cortex.

We next calculated differences in the expression of specific RNA isoforms for genes robustly detected (>20 TPM) in both the fetal and adult cortex (see **Methods**). Of note, we identified 1,424 transcripts mapping to 1,083 genes that were classified as ‘fetal specific’ (i.e. they were not detected in the adult cortex). Likewise, 1,062 transcripts mapping to 798 genes were classified as ‘adult specific’ (i.e. they were not detected in the fetal cortex). For 222 genes (6.09% out of 3,648 detected genes with at least one transcript showing >20 TPM), we identified a switch in the dominant isoform - i.e. differential transcript usage - between human fetal and human adult cortex (**Supplementary Table 16**). These transcripts were characterized by having (a) significant differences in transcript abundance between fetal and adult cortex (Mann-Whitney-Wilcoxon test P < 0.05), (b) fetal cortex specific expression (TPM > 20 in fetal cortex and TMP = 0 in adult cortex) or (c) adult cortex specific expression (TPM > 20 in adult cortex and TMP = 0 in fetal cortex). The gene *MAP1B*, which is involved in essential microtubule assembly during neurogenesis^51^, was characterized by the largest expression difference in dominant transcripts between adult and fetal specific RNA isoforms, with both transcripts differing at their 3’ untranslated region (UTR) (**Supplementary Figure 38**). Also of note is differential transcript usage for *SNAP25*, a gene involved in synaptic function^52^, which is characterized by strong expression of a transcript consisting of 8 exons (ENST00000304886.6) in the fetal cortex that is downregulated in the adult cortex in conjunction with increased expression of an alternative transcript incorporating an alternate fifth exon (ENST00000254976.6) (**Supplementary Figure 39**).

A similar frequency of AS events was observed in the human adult and human fetal cortex (adult: 5,660 unique AS genes with 19,379 AS events; fetal: 4,852 unique AS genes associated with 15,405 AS events), with considerable overlap (3,408 annotated genes (60.2% of AS genes in adult cortex, 70.2% of AS genes in fetal cortex)) (**Figure 5**). IR was more prevalent in the fetal cortex (3,053 transcripts associated with 1,662 genes (17.2% of annotated genes)) than adult human cortex (2,557 transcripts associated with 1,435 genes (13.1% of annotated genes)) (odds ratio = 1.50, P = 3.22 × 10^−46^, Fisher’s exact test), corroborating previous studies suggesting that IR plays a role in the developmental regulation of gene transcription in the brain^53^. Furthermore, although genes with IR-transcripts were generally more lowly expressed, they were more highly expressed in the fetal cortex than the adult cortex (W = 1.1 × 10^6^, P = 2.87 × 10^−8^, **Supplementary Figure 40**). GO analysis of the 913 genes uniquely associated with IR in the fetal cortex showed that the most enriched molecular function was also ‘RNA binding’ (odds ratio = 1.99, adjusted P = 5.6 × 10^−11^, **Supplementary Table 4**).

### Differential transcript usage across human fetal brain regions

We next generated Iso-Seq data on two additional fetal brain regions (hippocampus and striatum) from matched donors (**Supplementary Table 1**). Although the sequencing depth for these additional brain regions was lower than that obtained for the fetal cortex (fetal hippocampus: 0.483M CCS reads, 8,416 transcripts mapping to 5,606 genes; fetal striatum: 0.547M CCS reads, 9,678 transcripts mapping to 6,035 genes, **Supplementary Table 17**), we used these datasets to explore fetal transcriptional differences across the hippocampus, striatum and cortex. Amongst transcripts mapping to annotated genes, we again identified both known (hippocampus: n = 7,261 (86.28%) transcripts; striatum: n = 8,118 (83.88%) transcripts) and novel (hippocampus: n = 1,155 (13.72%) transcripts; striatum: n = 1,560 (16.12%)) transcripts (**Supplementary Table 17**). As expected, there was considerable overlap in genes detected across the three brain regions (3,385 transcripts associated with 2,650 genes based on TPM>20), although a notable subset of transcripts were uniquely expressed in each brain region (cortex: n = 2,180; hippocampus: n = 1,502; striatum: n = 2,346, **Supplementary Figure 41**). We further identified striking evidence for differential transcript usage across brain regions for a subset of genes; dominant isoform switches between cortex and hippocampus (n = 5 genes), cortex and striatum (n = 19 genes) and striatum and hippocampus (n = 6 genes) were observed (**Supplementary Table 18**). For example, *MEF2C*, a gene regulating excitatory neuronal development and associated with intellectual disability and autism^54^, expressed different primary RNA isoforms in the cortex and hippocampus; a ~2kb transcript consisting of 11 exons (ENST00000514015.5) was detected in the cortex, whereas a ~7kb transcript also consisting of 11 exons but an extended 3’UTR (ENST00000504921.6) was detected in the hippocampus (**Supplementary Figure 42**).

### Widespread isoform diversity in genes associated with brain disease

Alternative splicing has been increasingly implicated in health and disease, and is recognized to play a prominent role in brain disorders hypothesized to involve the cerebral cortex including autism^12^, schizophrenia (SZ)^13^ and AD^14^. There has been considerable progress in identifying genes associated with these disorders using genome sequencing and genome-wide association study (GWAS) approaches^55^, although the full repertoire of RNA isoforms transcribed from these genes in the cortex has not been systematically characterized. First, we used the human GWAS catalogue database^23^ to interrogate the most transcriptionally diverse genes in the human cerebral cortex, finding them to be enriched for genes implicated in relevant GWAS datasets (‘Alzheimer’s disease (late onset): odds ratio = 6.25, P = 0.04, ‘autism spectrum disorder or schizophrenia’: odds ratio = 2.39, P = 0.01, ‘schizophrenia’: odds ratio = 2.49, P = 0.005, **Supplementary Table 4**). Second, we assessed RNA isoform diversity in genes robustly associated with AD (three familial AD genes^56^ and 59 genes nominated from the most recent GWAS meta-analysis^57,58^), autism (393 genes nominated as being category 1 (high confidence) and category 2 (strong candidate) from the SFARI Gene database https://gene.sfari.org/), and SZ (339 genes nominated from the most recent GWAS meta-analysis^59^). Amongst disease-associated genes detected in the cortex, we found evidence for considerable isoform diversity (human cortex: 2765 transcripts were mapped to 619 disease-associated genes; mouse cortex: 3,846 transcripts were mapped to 687 disease-associated genes, **Supplementary Table 19**). The vast majority of disease-associated genes detected in the cortex were characterized by more than one RNA isoform in both the human (n = 472 (76.3%) genes) and mouse cortex (n = 561 (81.6%) genes). *MEF2C* (AD-associated) and *TCF4* (autism- and SZ-associated) were the most “isoformic” disease genes in both human (*MEF2C*: n = 19 isoforms; *TCF4*: n = 40 isoforms) and mouse (*Mef2c*: n = 36 isoforms; *Tcf4*: n = 83 isoforms) cortex; of note, both genes have been shown to be key members of transcriptional networks associated with neuropsychiatric disease^60^. Importantly, a large number of the transcripts mapping to disease-associated genes had not been previously annotated in existing databases in human (n = 994 (35.9%) isoforms) and mouse cortex (n = 1,654 (43%) isoforms), identifying novel transcripts that may have potential relevance to understanding neurodegenerative and neuropsychiatric disorders. Interestingly, transcripts from disease-associated genes were characterized by a relatively high level of IR in the human cortex (AD: n = 8 (25.8%); autism: n = 68 (22.2%); SZ: n = 75 (26.7%)) with a large proportion of these annotated IR-transcripts being predicted for NMD (AD: n = 5 (62.5% of IR-genes), autism: n = 27 (39.7% of IR-genes), SZ: n = 26 (34.7%, **Supplementary Table 20**). Given the links between fusion transcripts and disease^61^, a number of disease-associated genes were involved in fusion events in the human cortex (autism: n = 3, *MECP2-IRAK1*, *ELAC1-SMAD4*, *BAZ2B-AC009961.1*; SZ: n = 2, *EDC4*-*NRN1L*, *GPD3-MAPK3*; autism- and SZ-associated: *FOXG1-LINC01551*, **Figure 6, Supplementary Figure 43**). Given the hypothesized role of neurodevelopment and ageing in autism, SZ and AD, it is notable that we found striking differences in isoform diversity between human adult and human fetal cortex for many disease-associated genes (**Supplementary Table 20**).

**Figure 6:**
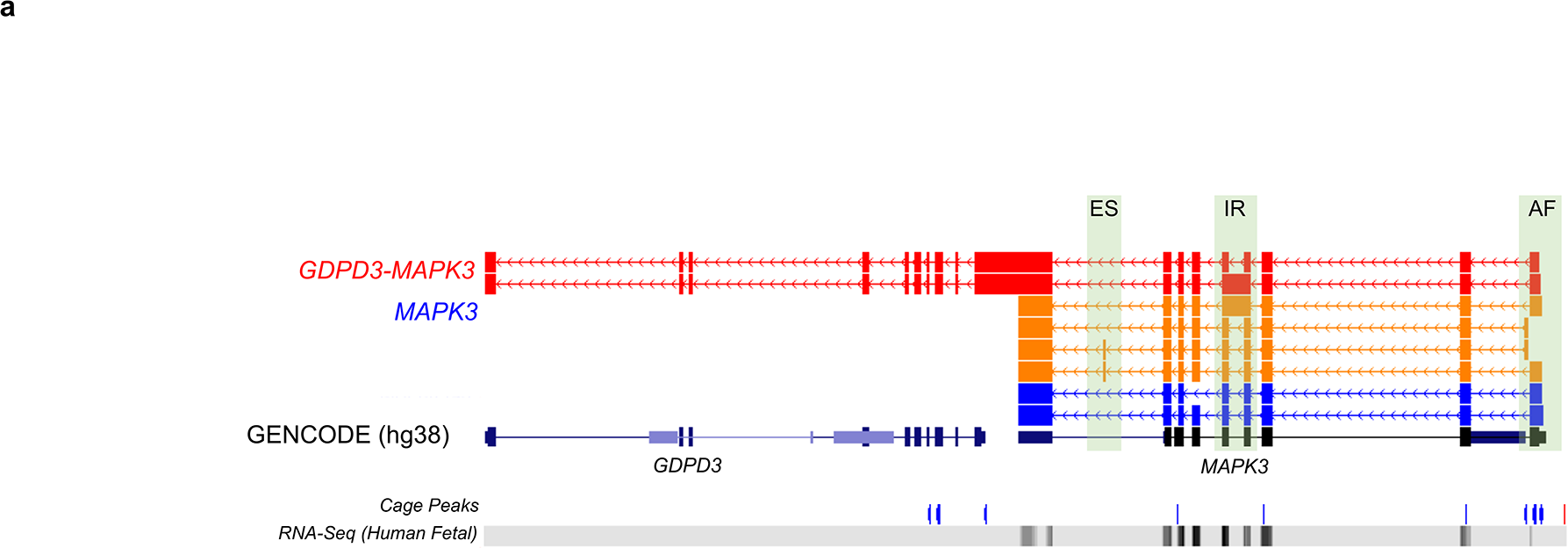
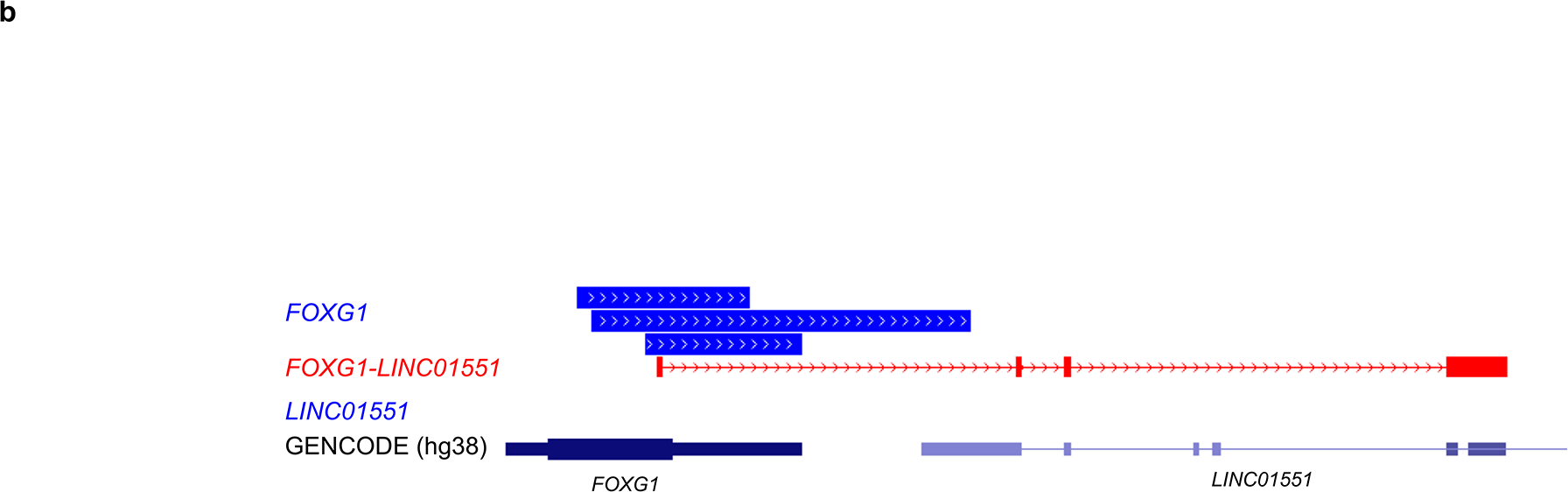
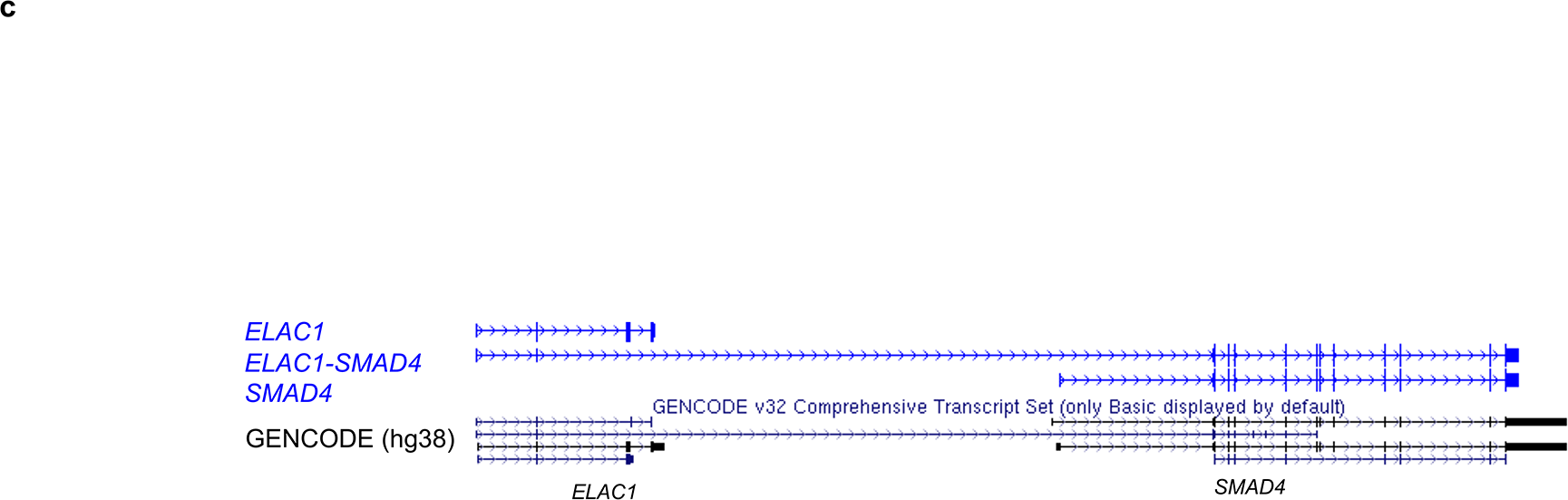
Examples of fusion transcripts in the human cortex incorporating exons from disease-associated genes. **a)** Two read-through transcripts incorporating exons from the SZ-associated gene *MAPK3* and *GDPD3* in the human cortex. Of note, one of the fusion transcripts contained an intron retention, as observed in another novel isoform of *MAPK3*. **b)** A fusion transcript incorporating exons from *FOXG1* (autism- and SZ-associated) and *LINC01551* in the human cortex. **c)** A fusion transcript incorporating exons from *ELAC1* (autism-associated) and *SMAD4* in the human cortex. Figures were taken from UCSC genome browser track and were adapted to display only the transcript of interest, colour-coded by *SQANTI2* categories (blue – FSM, orange – NIC, red – NNC, purple – antisense). SZ – Schizophrenia, ES – Skipped exon, IR – Intron retention, AF – Alternative first exon. FSM – Full splice match, ISM – Incomplete splice match, NIC – Novel in catalogue, NNC – novel not in catalogue. Isoforms are coloured based on *SQANTI2* classification categories (blue = FSM, cyan = ISM, red = NIC, orange = NNC, purple = antisense). FSM – Full Splice Match, ISM – Incomplete Splice Match, NIC – Novel In Catalogue, NNC – Novel Not in Catalogue.

## Discussion

We used long-read isoform sequencing to characterize full-length cDNA sequences and generate detailed maps of alternative splicing in the human and mouse cerebral cortex. We identify considerable RNA isoform diversity amongst expressed genes in the cortex, including many novel transcripts not present in existing genome annotations. Of note, we detect full-length transcripts from several previously unannotated genes in both the human and mouse cortex, and many examples of fusion transcripts incorporating exons from neighbouring genes. Although global patterns of RNA isoform diversity appear to be generally similar between human and mouse cortex, we identified some notable exceptions with certain genes showing species-specific transcriptional complexity. Furthermore, we also identify some striking developmental changes in transcript diversity, with certain genes characterized by differential transcript usage between fetal and adult cortex. Importantly, we show that genes associated with autism, schizophrenia and Alzheimer’s disease are characterized by considerable RNA isoform diversity, identifying novel transcripts that might play a role in pathology. Our data, which are available as a browsable resource for the research community (see **Resources**), confirm the importance of alternative splicing in the cortex and highlight its role as an important mechanism underpinning gene regulation in the brain.

Our findings highlight the power of novel long-read sequencing approaches for transcriptional profiling. By generating reads spanning entire transcripts it is possible to systematically characterize the repertoire of expressed RNA isoforms and fully assess the prevalence of alternative splicing. To our knowledge, our analysis represents the most comprehensive characterization of full-length transcripts and isoform diversity in the cerebral cortex yet undertaken. Several findings are particularly notable. First, we highlight that existing gene annotations are incomplete and that novel transcripts are likely to exist for a large proportion of expressed genes. Our data show examples of novel exons and even entire genes not currently annotated in existing databases. Importantly, it has been shown that such incomplete annotation has a disproportionate impact on our understanding of Mendelian and complex neurogenetic disorders^62^. Our resource enhances our understanding of the repertoire of expressed transcripts in the cerebral cortex. Second, we show that read-through RNA transcripts (or gene fusion transcripts) - formed when exons from two genes fuse together - occur at relatively high levels in the cortex. Although many of these fusion transcripts appear to be associated with NMD, many have the potential to be translated into proteins or may have a regulatory effect at the RNA level. Despite gene fusion transcripts having a well-documented role in several human cancers^32^, the systematic analysis of gene fusion and read-through transcripts has been limited to date given the limitations of existing short-read sequencing technologies^63^. Our data support recent data suggesting that read-through transcripts occur naturally^64^, and suggest that some fusion transcripts may have protein-coding potential, with important implications for brain disease. Third, we are able to highlight the extent to which alternative splicing events make a major contribution to isoform diversity in the cortex. In particular we show that IR is a relatively common form of alternative splicing in the cortex that is associated with reduced expression and NMD. Importantly, IR was more prevalent in the fetal human cortex than adult human cerebral cortex, supporting previous studies suggesting that IR plays a role in the developmental regulation of gene transcription in the brain^45^. Finally, we highlight major developmental changes in cortical isoform abundance in the human brain. In particular, we identify striking examples of transcript usage between fetal and adult cortex, and also highly differences in isoform expression between different regions of the human brain.

Our results should be interpreted in the context of several limitations. First, we profiled tissue from a relatively small number of human and mouse donors. Although we found highly consistent patterns of alternative splicing across these biological replicates and rarefaction curves confirmed our sequencing dataset was close to saturation, we were unable to explore inter-individual variation in alternative splicing. Nonetheless, we were able to identify transcripts from very lowly-expressed genes (such as the antisense novel gene upstream of *E2F3* (**Figure 4**) upon merging multiple Iso-Seq datasets that would have otherwise been filtered out due to low read count. Recent studies have highlighted considerable evidence for genetic influences on isoform diversity in the human cortex, with splicing quantitative trait loci (sQTL) widely implicated in health and disease^65^. Future work will aim to extend our analyses to larger numbers of samples to explore population-level variation in transcript abundance in the cerebral cortex and differences associated with pathology. Second, despite the advantages of long-read sequencing approaches for the characterization of novel full-length transcripts, these methods are often assumed to be less quantitative than traditional short-read RNA sequencing methods^66^. We implemented a stringent QC pipeline (see **Methods**) and undertook considerable filtering of our data, finding high consistency across biological replicates and validating our findings using complementary approaches (i.e. nanopore sequencing, RNA-Seq, and by comparison to existing genomic databases).

We show that transcriptional profiles generated using Iso-Seq reflect those expected from the tissues we assessed (i.e. the cerebral cortex), and we found a strong correlation with both gene- and transcript-level expression measured using short-read RNA-Seq on the same samples. Given that we have adopted stringent QC approaches, many true transcripts from our final dataset - particularly lowly-expressed transcripts, are likely to have been filtered out - our analyses probably underestimate the extent of RNA isoform diversity in the cerebral cortex; therefore, we also provide a less conservatively-filtered dataset for download from our online track hub (see **Web Resources**). Third, our analyses were performed on ‘bulk’ cortex tissue containing a heterogeneous mix of neurons, oligodendrocytes and other glial cell-types. It is likely that these different cell-types express a specific repertoire of RNA isoforms and we are not able to explore these differences in our data. Of note, novel approaches for using long-read sequencing approaches in single cells will enable a more granular approach to exploring transcript diversity in the cortex. Although such approaches are currently limited by technological and analytical constraints, a recent study used long-read transcriptome sequencing to identify cell-type-specific transcript diversity in the mouse hippocampus and prefrontal cortex^67^. Finally, although we explored the extent to which novel transcripts contained ORFs, the extent to which they are actually translated and contribute to cortical proteomic diversity is not known.

In summary, our data confirm the importance of alternative splicing and alternative first exon usage in the cerebral cortex, dramatically increasing transcriptional diversity and representing an important mechanism underpinning gene regulation in the brain. We highlight the power of long-read cDNA sequencing for completing our understanding of human gene annotation, and our transcript annotations, isoform data, and Iso-Seq analysis pipeline are available as a resource to the research community (see **Web Resources**).

## Methods

### Samples

Adult human prefrontal cortex tissue was obtained from the MRC London Neurodegenerative Diseases Brain Bank (https://www.kcl.ac.uk/ioppn/depts/bcn/our-research/neurodegeneration/brain-bank). Subjects were approached in life for written consent for brain banking, and all tissue donations were collected and stored following legal and ethical guidelines (NHS reference number 08/MRE09/38; the UK Human Tissue Authority HTA license number 12293). Fetal human brain tissue from three brain regions (frontal cortex, hippocampus and striatum) was obtained from the Human Developmental Biological Resource (HDBR) (http://www.hdbr.org). Ethical approval for the HDBR was granted by the Royal Free Hospital research ethics committee under reference 08/H0712/34 and HTA material storage license 12220. For each human sample, ~20mg of flash frozen tissue was homogenized in Trizol (Thermo Fisher Scientific, UK) and RNA isolated using Direct-zol columns (Zymo, USA). Mouse entorhinal cortex tissue was dissected from eight wild-type mice from a colony housed at Eli Lilly and Company (Windlesham, UK) in accordance with the UK Animals (Scientific Procedures) Act 1986 and with approval of the local Animal Welfare and Ethical Review Board. For each mouse sample, RNA was isolated using the AllPrep DNA/RNA Mini Kit (Qiagen, UK) from ~5mg tissue. RNA samples were quantified using the Nanodrop 1000 spectrophotometer and RNA integrity numbers (RIN) derived using a Bioanalyzer 2100 (Agilent, UK). Additional details on mouse breeding conditions can be found in Castanho et al. (2020)^68^, and further details on all samples used in this study are provided in **Supplementary Table 1**.

### Whole transcriptome Iso-Seq library Preparation and SMRT sequencing

First strand cDNA synthesis was performed on ~1μg RNA using the SMARTer PCR cDNA Synthesis Kit (Clontech, UK) followed by PCR amplification with PrimeSTAR GXL DNA Polymerase (Clontech, UK). Optimal PCR cycle number was determined through collection of 5μl aliquots during every two cycles of a test PCR and assessed on a 1% Agarose gel electrophoresis. Large-scale PCR was subsequently performed using the optimal number of cycles and the resulting amplicons divided into two fractions and purified with 0.4X and 1X Ampure PB beads (PacBio, USA). Quantification and size distribution of each fraction was then determined using the Qubit DNA High sensitivity assay (Invitrogen, UK) and Bioanalyzer 2100 (Agilent, UK). The two fractions were recombined at equimolar quantities and library preparation performed using SMRTbell Template Prep Kit v1.0 (PacBio, USA). Sequencing was performed on the PacBio Sequel 1M SMRT cell. Samples were processed using either the version 3 chemistry (parameters: diffusion loading at 5pM, pre-extension 4 hours, Capture time 20 hours) or version 2.1 chemistry (parameters: magbead loading at 50pM with a 2 hour pre-extension and 10 hour capture).

### SMRT sequencing quality control (QC) and data processing

QC of raw reads was performed using SMRT Link Portal v7.0, with subsequent analysis using the *Iso-Seq3.1.2* pipeline^18^. Briefly, CCS reads were generated from a minimum of 1 pass (*Iso-Seq3 CCS*, v3.4.1). cDNA primers and SMRT adapters were then removed using *Lima* (v1.9) to generate full-length (FL) reads, followed by removal of artificial concatemers reads and trimming of polyA tails in *Iso-Seq3 Refine*. Full-length, non-concatemers (FLNC) reads were then collapsed, according to default parameters in *Iso-Seq3 Cluster*, to high-quality transcripts. *Cupcake* scripts were subsequently applied using default parameters^21^, to reduce redundancy, remove 5’ degradation products and chain samples for downstream comparison. High-quality, full-length transcripts were then mapped to the human (hg38, GENCODE v31) or mouse (mm10, GENCODE vM22) reference genome using *minimap2*^69^ (v2.17) with the following parameters “-ax splice -uf --secondary=no -C5 -O6,24 -B4”^21^.

### RNA-Seq library Preparation, Illumina sequencing and data processing

RNA from a subset of human fetal (n = 3) and mouse (n = 8) cortex tissue samples was prepared with TruSeq Stranded mRNA Sample Prep Kit (Illumina) and subjected to 125bp paired-end sequencing using a HiSeq2500 (Illumina). Briefly, cDNA libraries were prepared from ~450ng of total RNA plus ERCC spike-in synthetic RNA controls (Ambion, dilution 1:100), purified using Ampure XP magnetic beads (Beckman Coulter) and profiled using D1000 ScreenTape System (Agilent). Raw sequencing reads, with Phred (Q) ≥ 35, were trimmed (ribosomal sequence removal, quality threshold 20, minimum sequence length 35) using *fastqmcf* (v1.0), yielding a mean untrimmed read depth of ~20 million reads/sample. Subsequent filtered reads were then mapped to the human (hg38) or mouse (mm10) reference genome using *STAR*^70^ (v1.9). Gene and transcript expression were determined by aligning merged RNA-Seq reads to RNA isoforms (*Cupcake* collapsed) from Iso-Seq datasets using *Kallisto*^71^ (v0.46.0) with default parameters, as input to *SQANTI2*^18^.

### Transcriptome Annotation and filtering

Isoforms detected using SMRT sequencing were characterized and classified using *SQANTI2* (v7.4) in combination with GENCODE (human v31, mouse vM22) comprehensive gene annotation, FANTOM5 CAGE peaks^25^ (human – hg38, mouse – mm10), polyA motifs, Intropolis junction dataset^53^ or *STAR* output junction file, FL read counts (abundance file), and *Kallisto* counts from mouse and fetal RNA-Seq data. An RNA isoform was classified as FSM if it aligned with reference genome with the same splice junctions and contained the same number of exons, ISM if it contained fewer 5’ exons than reference genome, NIC if it is a novel isoform containing a combination of known donor or acceptor sites, or NNC if it is a novel isoform with at least one novel donor or acceptor site. Depictions of RNA isoform classifications can be found in **Figure 1**. Potential artifacts such as reverse transcription jumps or intrapriming of intronic lariats were filtered out using the *SQANTI2* filter script with an intrapriming rate of 0.6. Identification of fusion transcripts, intron retention, polyA motifs and proximity to CAGE peaks were defined based on *SQANTI2* filtered isoforms. The occurrence of mutually exclusive exons (MX) and skipped exons (SE) were assessed using *SUPPA2*^42^ with the parameter -f *ioe*, intron retention (IR) with *SQANTI2*, and alternative first exons (AF), alternative last exons (AL), alternative 5’ splice sites (A5), and alternative 3’ splice sites (A3) using custom scripts based on splice junction coordinates. Classification of isoforms as lncRNA (long non-coding RNA) was performed by using *SQANTI2* in combination with GENCODE (human - v31, mouse - vM22) long non-coding RNA gene annotation.

### Comparison of RNA-Seq and Iso-Seq Expression

Gene and transcript expression between Iso-Seq and RNA-Seq were compared for human and mouse cortex using *SQANTI2* output, with Kallisto expression file as input. Iso-Seq gene expression was determined with the summation of associated transcript FL read counts, with mono-exonic transcripts removed, and normalised to TPM (calculated from FL read counts/total transcriptome counts * 1,000,000). RNA-Seq gene and isoform expression was determined from alignment of RNA-Seq reads to Iso-Seq isoforms, generated from *Cupcake*^21^ scripts, using *Kallisto*. For more stringent investigation of the relationship between the gene length, number of exons (determined by representative longest transcript) and the number of transcripts, an Iso-Seq gene expression threshold (> 2.5 Log_10_ TPM) was applied. This threshold was selected based on the gene expression that gave the most statistically-significant correlation between human and mouse isoform number (**Supplementary Table 21**).

### Gene ontology

*EnrichR*^23^ based gene enrichment analysis was performed on three sets of analyses: i) top 500 most abundantly expressed genes, ii) 100 most isoformic genes and iii) genes with intron-retained transcripts in human and mouse Iso-Seq cortical datasets. Iso-Seq gene expression was calculated as before from summation of associated transcript FL read counts within the *SQANTI2* classification file. The functional categories examined were: GO_Biological_Process, GO_Cellular_Component, GO_Molecular_Function, Panther_2016, Human_Gene_Atlas, ARCHS4_Tissues and GWAS_Catalogue_2019. Mouse_Gene_Atlas was used for the mouse transcriptome. Genes with IR-transcripts were further assessed with GOLiath (https://www.bioinformatics.babraham.ac.uk/goliath/goliath.cgi), the latter to enable a custom gene background derived from the expressed genes listed in the *SQANTI2* classification file.

### Human and Mouse Comparison

For appropriate comparison of the Iso-Seq datasets, the mouse gene names were converted to the equivalent homologous human gene names according to mouse genome informatics syntenic gene list (http://www.informatics.jax.org/downloads/reports/HOM_MouseHumanSequence.rpt), considering only mouse-specific homologous genes. 17,042 genes were identified from the list as homologous, of which 267 genes from the human homologous list and 282 genes from the mouse homologous list were removed due to cross genome annotation. This gene set was then used to determine the relationship between human and mouse cortex of Iso-Seq gene expression and number of identified isoforms (**Supplementary Table 15**).

### Iso-Seq human dataset comparisons

Each human Iso-Seq dataset was assigned a unique transcript ID. We used *Cupcake’s* chain_samples.py^22^ script to merge full datasets (pre-*SQANTI2* filtered) to allow cross comparison followed by *SQANTI2* reannotation. FL read counts from each individual SMRT cell were extracted from read_stat.txt file and normalised to TPM (calculated from FL read counts/total transcriptome counts * 1,000,000) * 1,000,000). Testing for differential transcript usage between human fetal and adult samples was then performed with a Wilcoxon rank sum test. Transcripts annotated to a gene were then examined for a P < 0.05 with at least two transcripts with exclusive expression to fetal or adult samples respectively, each with a normalised expression level minimum of >20 TPM. Testing for differential transcript usage between fetal brain regions was first performed by determining differential transcript expression using a more stringent threshold (>100 TPM and no transcript expression in respective brain regions) - only two SMRT cells were generated from fetal hippocampus and striatum samples and thus limiting power of Wilcoxon rank sum test - and then by the detection of at least two transcripts with expression exclusive in two or more brain regions.

### Oxford Nanopore library preparation, sequencing and data processing

As validation, RNA from human fetal (n = 1) and human adult (n = 1) cortex was sequenced on Oxford Nanopore Technology (ONT). Maxima H Minus RT (Thermo Fisher Scientific) followed by 15 cycles of PCR using Takara LA Taq (Clontech) was used. Quantification and size distribution was then determined using Qubit DNA High sensitivity assay (Invitrogen) and Bioanalyzer 2100 (Agilent), and library preparation was proceeded with ONT’s PCR barcoding kit (SQK-PCB109). Sequencing was then performed on ONT PromethION using a FLO-PRO002 flow cell for human samples, and base-called using *Guppy* (v4.0). Resulting fastq files were processed through the *Pychopper/Pinfish*^72^ pipeline to produce polished transcripts, *SQANTI2* filtering was then applied

### Validation of Transcriptome Landscape

Novel transcripts, such as those not associated with GENCODE defined transcripts, were compared with publicly-available data from PacBio’s Alzheimer’s disease brain Iso-Seq dataset^30^, which was re-processed using the *Iso-Seq 3.1.2* pipeline^18^. Presence of CAGE peaks near novel transcripts were further checked, with liftOver (https://genome.ucsc.edu/cgi-bin/hgLiftOver) being performed on the mouse dataset to convert mm9 to mm10 genome coordinates.

### Web Resources

Gene transfer format files (GTF) from SQANTI2 were processed to a bigGenePred (https://genome.ucsc.edu/goldenPath/help/bigGenePred.html) followed by bed and bigBed format (https://genome.ucsc.edu/goldenPath/help/hubQuickStartSearch.html) to construct the hub. Further annotation enhancements were made to the bigBed files through an R script to extract SQANTI2 classification file structural categories and associated gene names to relabel and colour individual transcript transcripts. Separate colouring schemes are also made available to indicate level of expression.

## Data and code availability

Unprocessed PacBio Iso-Seq data has been deposited in the Sequence Read Archive (SRA) database (https://www.ncbi.nlm.nih.gov/sra) under accession numbers PRJNA664117 (human cortex) and PRJNA663877 (mouse cortex). Analysis scripts used for the analyses presented in this manuscript and browsable UCSC genome browser tracks of our processed Iso-Seq data (filtered and unfiltered) are available as a resource at: http://genome.exeter.ac.uk/BrainIsoforms.html.

## Acknowledgements

This work was supported by a grant from SFARI (Grant Number 573312, awarded to J.M.), a grant from the UK Medical Research (MR/R005176/1, awarded to J.M.) and a SFARI Bridge to Independence Award (NIMH R01-MH121521 and NICHD-P50-HD103557, awarded to M.J.G). S.K.L. is funded by a UK Medical Research Council CASE PhD studentship. The human fetal material was provided by the Joint MRC/Wellcome Trust (grant #099175/Z/12/Z) Human Developmental Biology Resource (www.hdbr.org). The human fetal material was provided by the Joint MRC/Wellcome Trust (grant #099175/Z/12/Z) Human Developmental Biology Resource (www.hdbr.org). Sequencing infrastructure was supported by a Wellcome Trust Multi User Equipment Award (WT101650MA) and Medical Research Council (MRC) Clinical Infrastructure Funding (MR/M008924/1). We acknowledge the help of R. Flynn from the University of Exeter Medical School for his help with screening fusion transcripts.

## Author contributions

A.R.J. and S.K.L. conducted long-read sequencing experiments. A.R.J, I.C. and K.M. conducted short-read RNA-seq experiments. J.D. provided technical assistance with sample preparation. N.J.B. provided fetal cortex RNA samples. K.M. advised on library preparation and aspects of sequencing. Z.A. provided mouse cortex tissue. J.M., E.H. L.S., and E.L.D. obtained funding. J.M. and A.R.J. designed the study. A.R.J. and S.K.L. undertook primary data analyses and bioinformatics, with analytical and computational input from E.H., P.O’N. and M.J.G. M.J.G. S.P., L.S., E.L.D., D.A.C. and N.J.B. helped interpret the results. A.R.J.,S.K.L. and J.M. drafted the manuscript. All authors read and approved the final submission.

