## Supplementary Figures for "Full-length transcript sequencing of human and mouse identifies widespread isoform diversity and alternative splicing in the cerebral cortex"

1. CCS Read per Iso-Seq dataset
2. Rarefaction Curves
3. Bioinformatics Pipeline
4. Number of genes in human and mouse, and of genes in fetal and adult
5. GO results of top 500 most abundantly expressed genes
6. Fetal: RNA-Seq vs Iso-Seq
7. Boxplot of number of genes per dataset
8. Tracks of *Meg3* in human cortex
9. Tracks of *TCF4* in mouse cortex
10. Correlation of gene length vs number of isoforms
11. Correlation of exons vs number of isoforms
12. Novel transcripts of annotated genes: categories
13. Novel vs annotated transcripts of annotated genes: expression
14. Novel vs annotated transcripts of annotated genes: length and exons
15. Cage peaks of novel transcripts
16. Junction support of annotated genes, novel transcripts
17. Transcripts supported by ONT
18. Fusion Gene across three genes
19. Novel vs Annotated genes; transcript length and gene expression
20. Further examples of antisense-sense pair fusion genes in mouse cortex
21. lncRNA vs non-lncRNA: length and exons
22. lncRNA vs non-lncRNA: expression
23. lncRNA vs non-lncRNA: ORF length
24. Proportion of splicing events
25. Gene overlap across each splicing event
26. Distribution of splicing events across fetal and adult cortex
27. Example of Alternative First in human cortex
28. Transcripts with NMD more lowly expressed than non-NMD
29. NMD enriched in IR transcripts
30. Isoform diversity in human adult and human fetal cortex, human and mouse
31. Tracks of *SORBS1* in human and mouse cortex
32. Tracks of *DLGAP1* in human and mouse cortex
33. Tracks of *NDUFS2* in human and mouse cortex
34. Tracks of *TEMEM191C* in human and mouse cortex
35. Frequency of splice sites in transcripts
36. Number of isoforms in human adult and human fetal cortex
37. Tracks of *MEG3* and *MBP* in human and mouse cortex
38. Tracks of *MAP1B*: Differential Transcript Usage
39. Tracks of *SNAP25*: Differential Transcript Usage
40. IR-gene expression in adult vs fetal cortex
41. Overlap of genes in hippocampus, striatum and cortex in fetal
42. Tracks of *MEF2C* in human and mouse cortex
43. Fusion transcripts associated with disease

**Supplementary Figure 1: Consensus distribution of CCS read lengths across all cortical samples with expected peak at 2-3kb, corresponding to mean length of mRNA in human and mouse cerebral cortex.** CCS for each sample from Iso-Seq Sequel in **a)** human adult cortex (n = 4 biologically independent samples, n = 5 SMRT cells), **b)** human fetal cortex (n = 3 biologically independent samples, n = 5 SMRT cells) and **c)** mouse cortex (n = 8 biologically independent samples, n = 8 SMRT cells) was generated using *Iso-Seq3* pipeline, with a minimum of 1 pass. Some of the human adult and human fetal cortex samples were sequenced more than once to maximise coverage (**Supplementary Table 1**). Number of CCS reads generated per SMRT cell can be found in **Supplementary Table 2**. CCS – Circular consensus sequence. SMRT – Single-molecule real-time

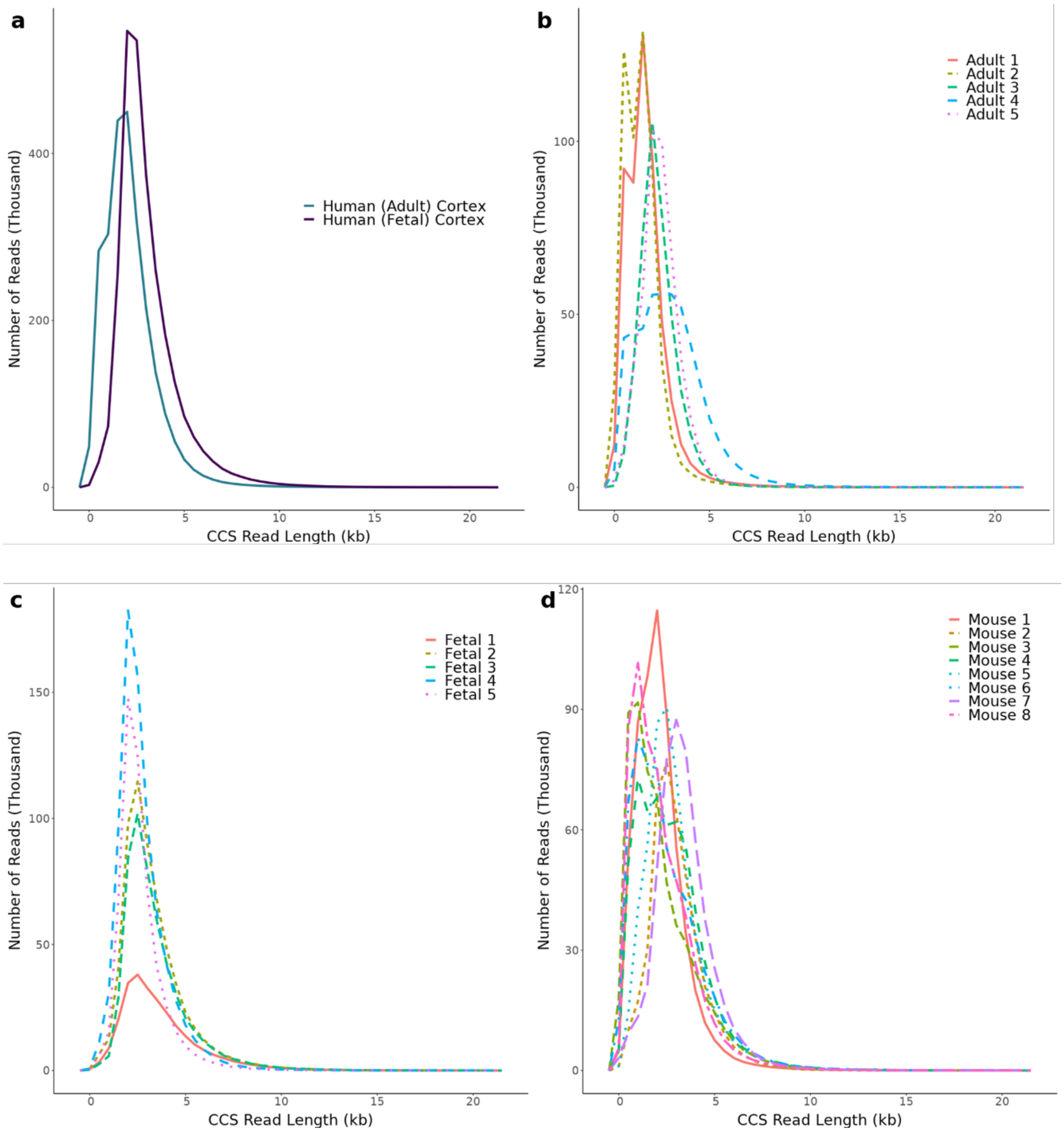

**Supplementary Figure 2: Saturation is reached across all Iso-Seq datasets at the gene and isoform level.** Subsampling to generate rarefaction curves, using *cDNA Cupcake* scripts, was performed on each dataset at a gene and isoform level in **a)** human adult (n = 4 biologically independent samples) and human fetal cortex (n = 3 biologically independent samples) and for each *SQANTI2* isoform category in **b)** human cortex, **c)** human adult cortex, **d)** human fetal cortex and **e)** mouse cortex (n = 8 biologically independent samples). FSM – Full splice match, ISM – Incomplete Splice Match, NIC – Novel In Catalogue, NNC – Novel Not in Catalogue.

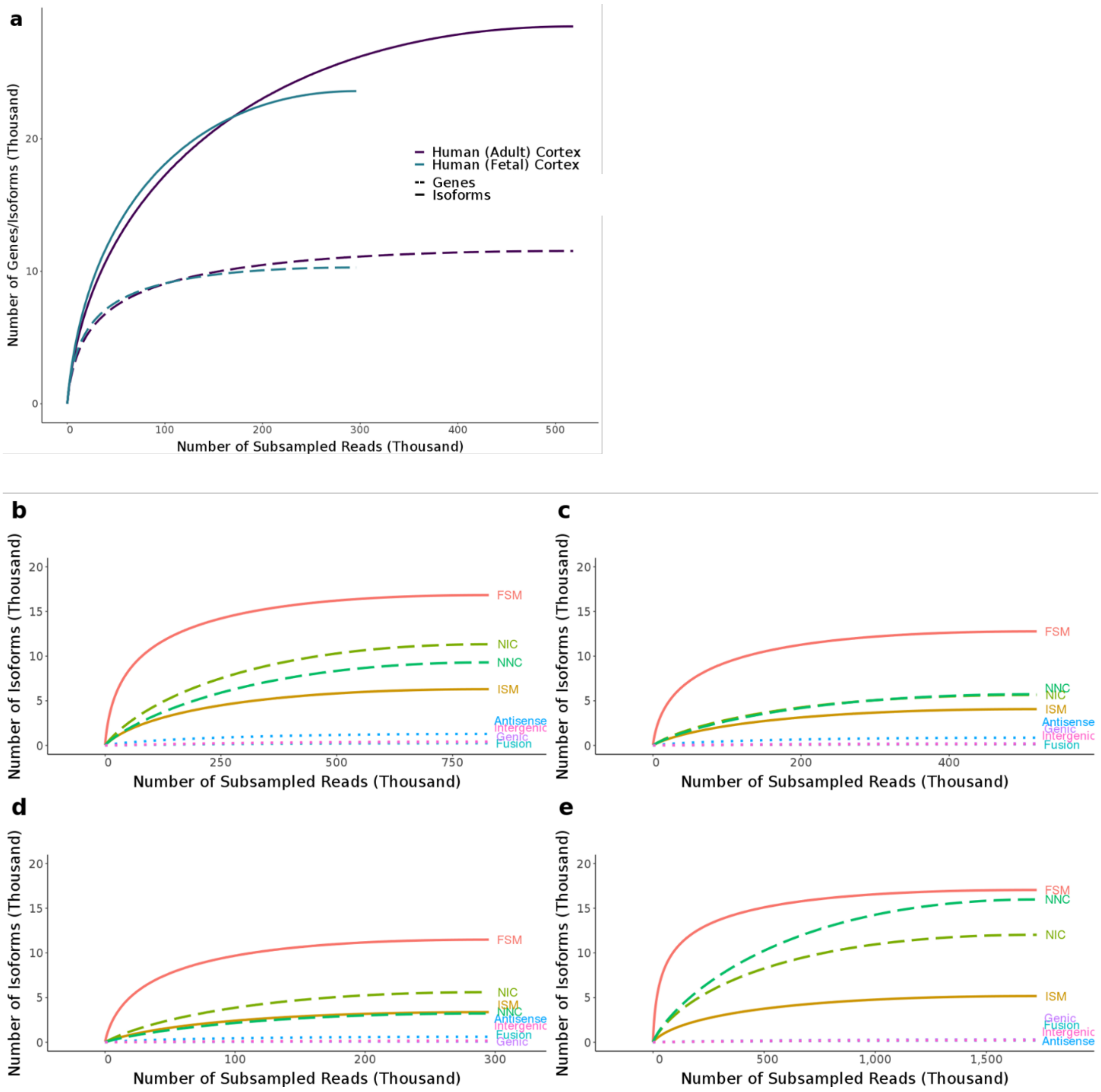

**Supplementary Figure 3: An overview of the analysis pipeline used to generate full-length transcriptional annotations in human and mouse cerebral cortex.** Polymerase reads from PacBio Sequel for each dataset were processed using *Iso-Seq 3.1.2* and *Cupcake* scripts to generate high quality, full-length isoforms. *SQANTI2* was used to fully annotate individual isoforms, with comparison to short-read RNA-Seq reads, nanopore sequencing (ONT), and reference annotation. A full overview of our methodological approach is given in the **Online Methods**. PacBio – Pacific Biosciences, ONT – Oxford Nanopore Technology

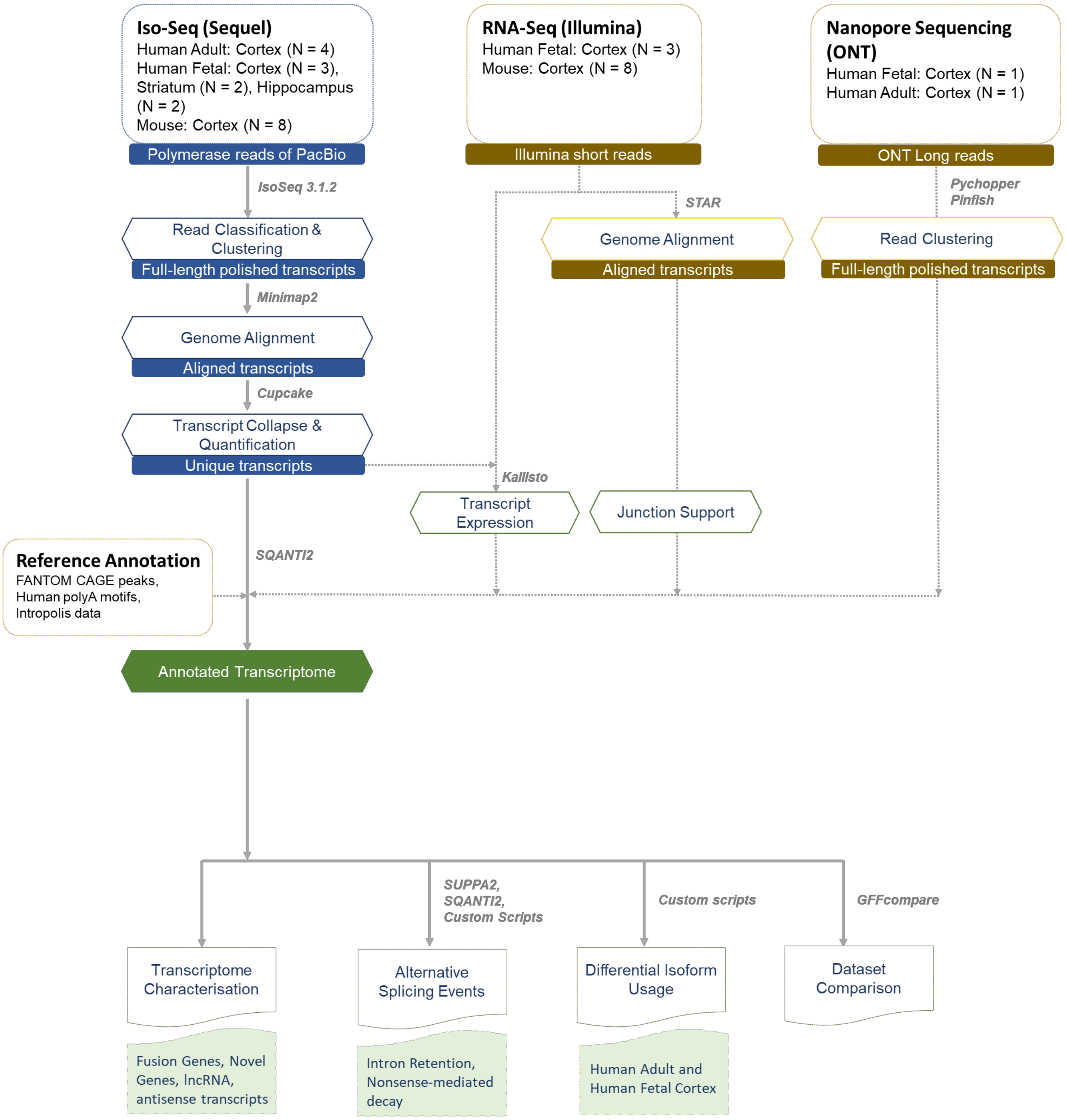

**Supplementary Figure 4: Overlap of genes detected in human and mouse cortical Iso-Seq datasets.** Shown is the number of common and unique annotated genes from GENCODE (human: hg38, mouse: mm10) identified in **a)** human and mouse cortex and **b)** human adult and human fetal cortex.

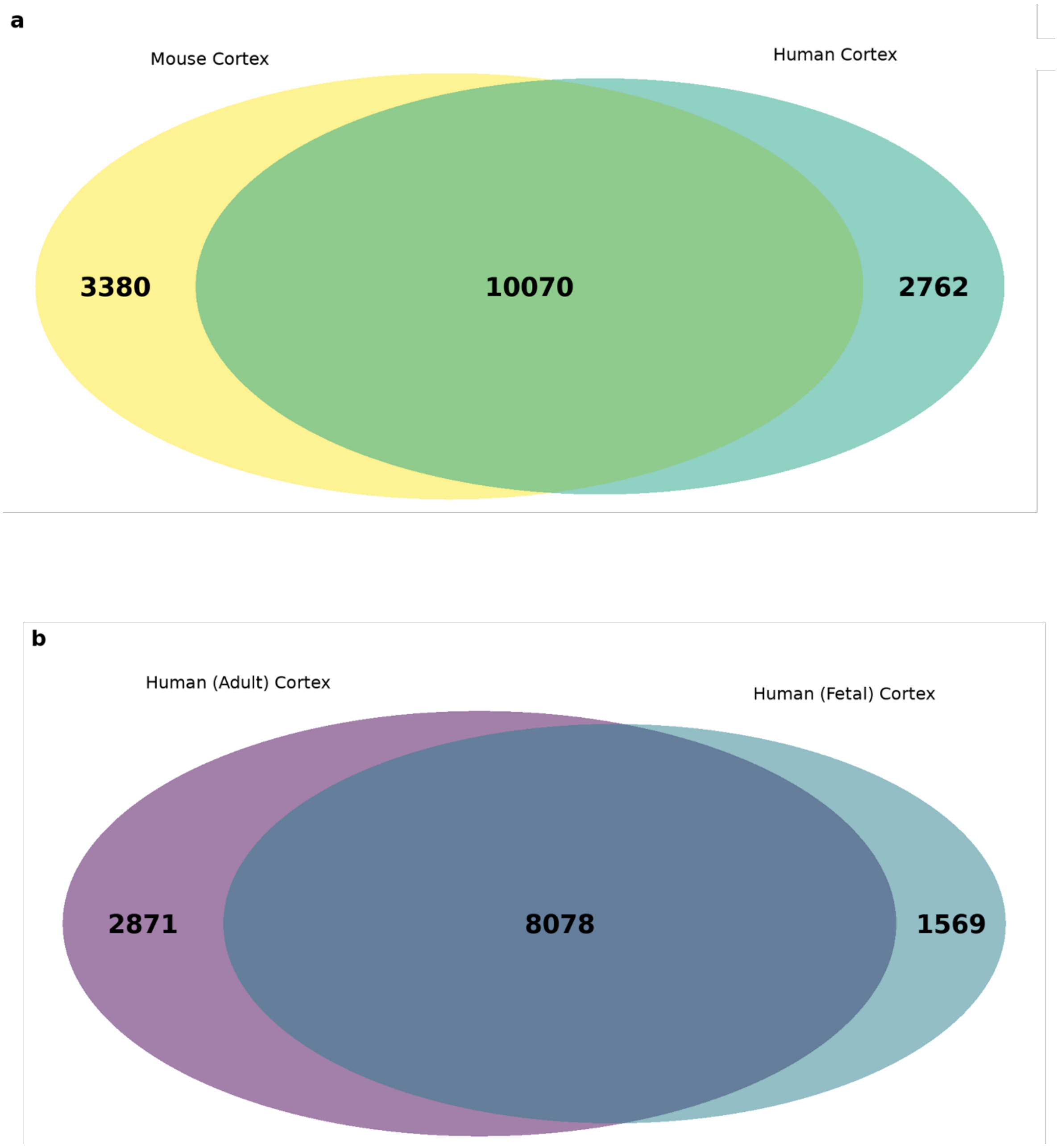

**Supplementary Figure 5: Gene expression patterns reflect expected cortical transcriptional profiles.** The most abundantly-expressed genes (top 500, ranked by TPM) were most significantly enriched for 'prefrontal cortex' genes (human cortex: odds ratio = 5.91, adjusted P = $1.93 \times 10^{-35}$, mouse cortex: odds ratio = 5.47, adjusted P = $1.58 \times 10^{-19}$). TPM – Transcripts per Million

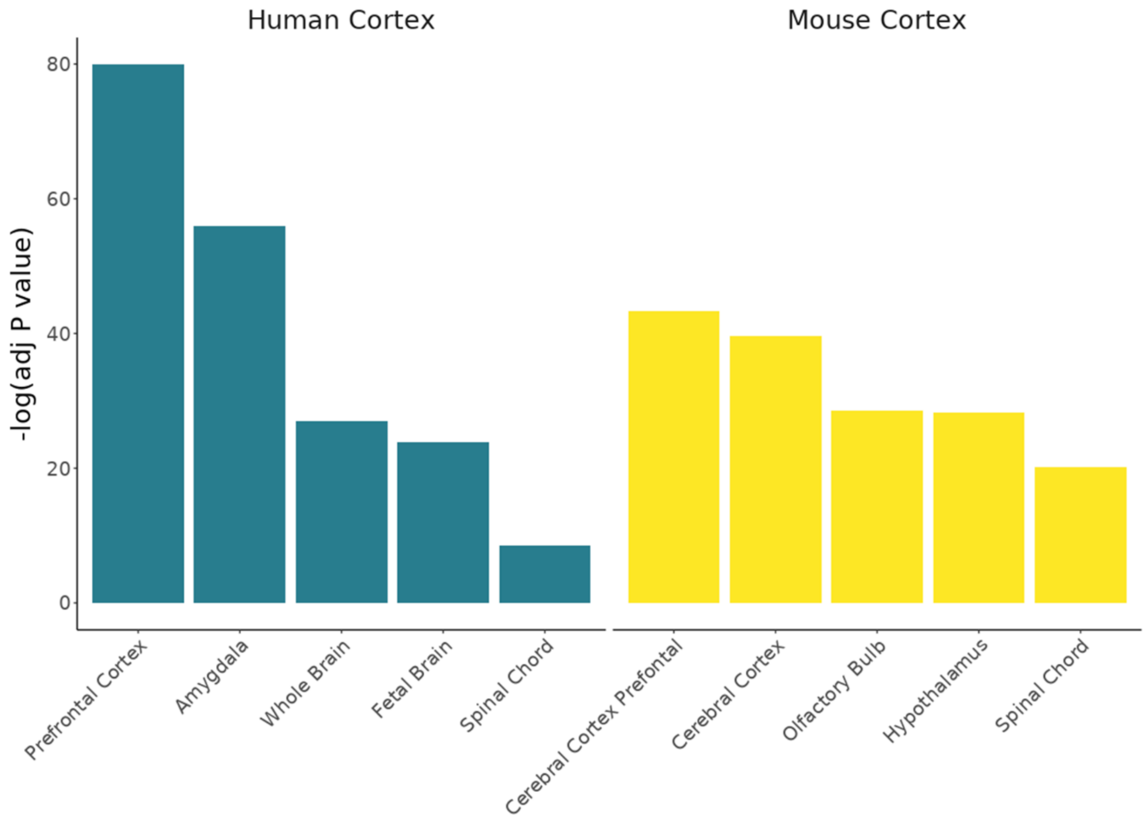

**Supplementary Figure 6: Long-read Iso-Seq data can accurately quantify levels of gene expression in the human cortex.** Shown is the relationship between expression estimated using RNA-Seq and Iso-Seq at the **a)** gene level (n = 9,223 genes, Pearson's correlation = 0.58, $P < 2.23 \times 10^{-308}$) and **b)** transcript level (n = 21,144 multiexonic transcripts, Pearson's correlation = 0.36, $P < 2.23 \times 10^{-308}$, RNA-Seq threshold of 0.01TPM) in the human cortex (data derived from three biologically independent fetal samples). RNA-Seq gene expression was determined after aligning short-read RNA-Seq to the Iso-Seq transcriptome. Iso-Seq gene expression was determined from the sum of full-length, multi-exonic transcript reads associated for each gene, with TPM values calculated by dividing the number of full length reads per gene by total full-length reads, multiplied by a million. The density of values is represented in increasing scale from light green to dark blue. Equivalent plots for the mouse cortex is shown in **Figure 2**. TPM – Transcripts per Million.

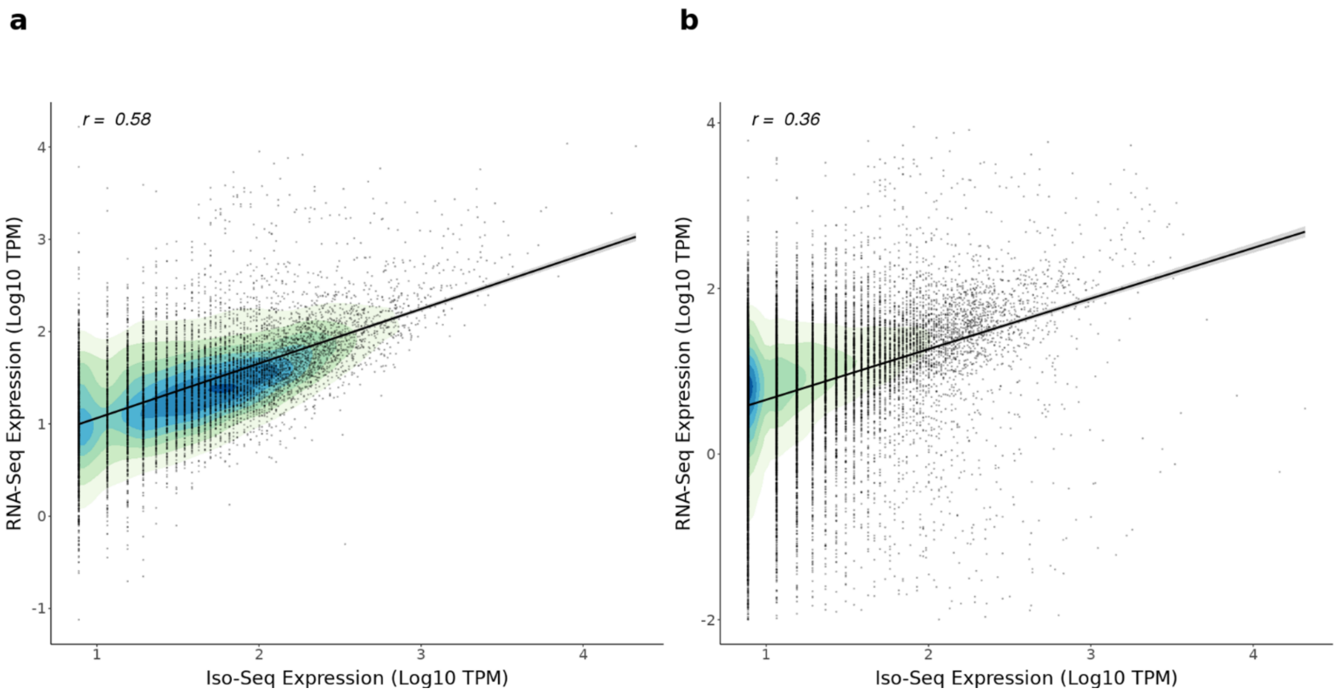

**Supplementary Figure 7: Widespread isoform diversity identified for many genes expressed in the human and mouse cortex.** Shown is the number of isoforms per gene identified in **a)** human and mouse (n = 8 biologically independent samples) cortex and in **b)** human adult (n = 4 biologically independent samples) and human fetal (n = 3 biologically independent samples) cortex.

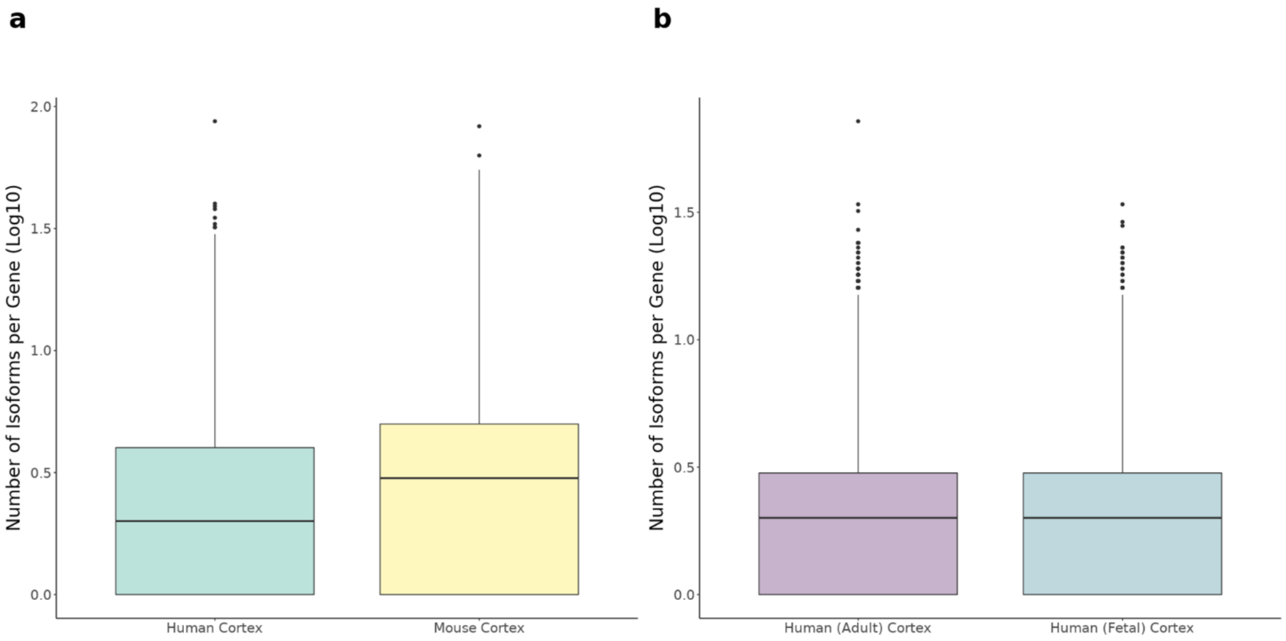

**Supplementary Figure 8: *MEG3* was the most transcriptionally diverse gene in human cortex with 87 isoforms**. Shown is a UCSC genome browser track of *MEG3* in the human cortex (n = 7 biologically independent samples), complemented with GENCODE reference genome (hg38), and RNA-Seq from a subset of samples (n = 3 biologically independent fetal samples). A maternally expressed imprinted long non-coding RNA (lncRNA) gene involved in synaptic plasticity, *MEG3* displayed the greatest isoform diversity. Isoforms are coloured based on *SQANTI2* classification categories (blue = FSM, cyan = ISM, red = NIC, orange = NNC). FSM – Full Splice Match, ISM – Incomplete Splice Match, NIC – Novel In Catalogue, NNC – Novel Not in Catalogue.

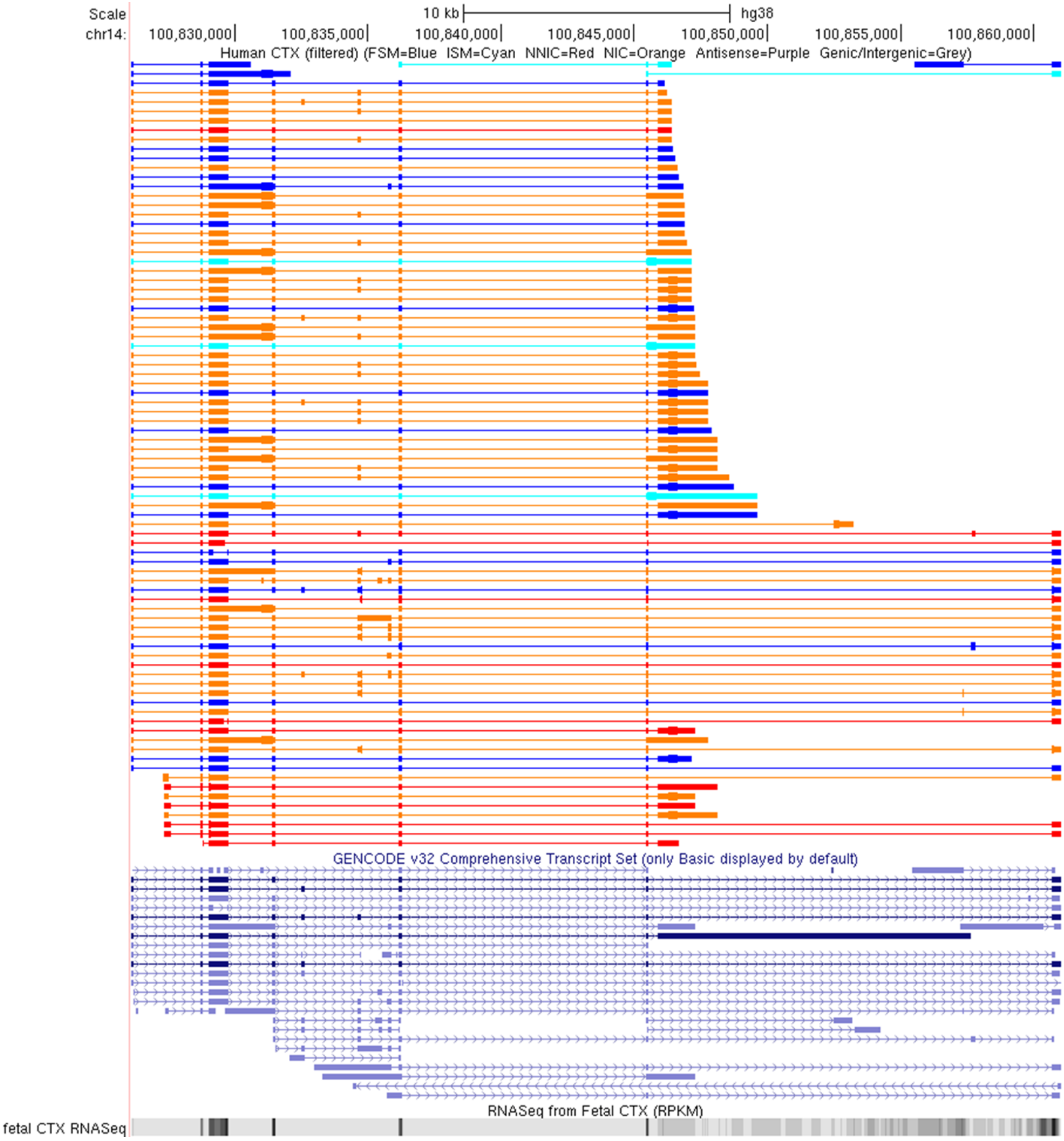

**Supplementary Figure 9: *Tcf4* was the most transcriptionally diverse gene in mouse cortex with 79 isoforms**. Shown is a track of *Tcf4* in the mouse cortex (n = 8 biologically independent samples), complemented with GENCODE reference genome (mm10), and matched RNA-Seq data (n = 8 biologically independent samples). A neurodevelopmental gene implicated in schizophrenia, *Tcf4* displayed the greatest isoform diversity in the mouse cortex. Isoforms are coloured based on *SQANTI2* classification categories (blue = FSM, cyan = ISM, red = NIC, orange = NNC). FSM – Full Splice Match, ISM – Incomplete Splice Match, NIC – Novel In Catalogue, NNC – Novel Not in Catalogue.

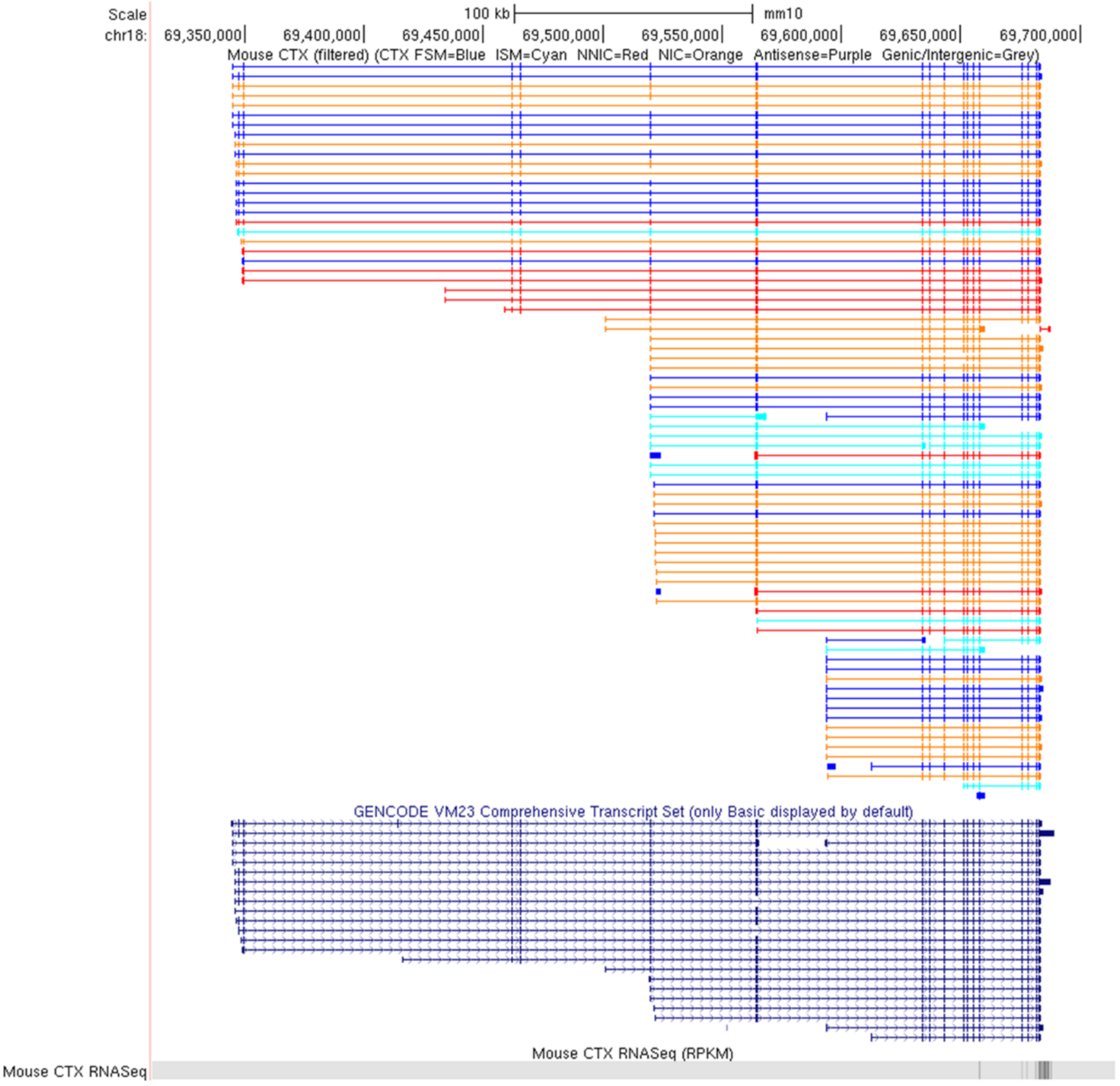

**Supplementary Figure 10: Longer genes are associated with more isoforms in human and mouse cortex.** The number of detected multi-exonic isoforms in **a)** human cortex (n = 7 biologically independent samples) and **b)** mouse cortex (n = 8 biologically independent samples) was correlated with gene length (human cortex: Pearson's correlation = 0.28, P = $1.71 \times 10^{-223}$; mouse cortex: Pearson's correlation = 0.32, P = $3.2 \times 10^{-301}$). A stronger relationship was observed among 'highly-expressed' genes (threshold of 2.5 $Log_{10}$ TPM) in both **c)** human cortex (Pearson's correlation = 0.43, P = $2.62 \times 10^{-27}$) and **d)** mouse cortex (Pearson's correlation = 0.45, P = $2.42 \times 10^{-31}$), consistent with the additional sensitivity of detecting isoforms in highly expressed genes. Gene length is represented by the longest isoform, and density of genes is represented in increasing scale from light green to dark blue. TPM – Transcripts per Million

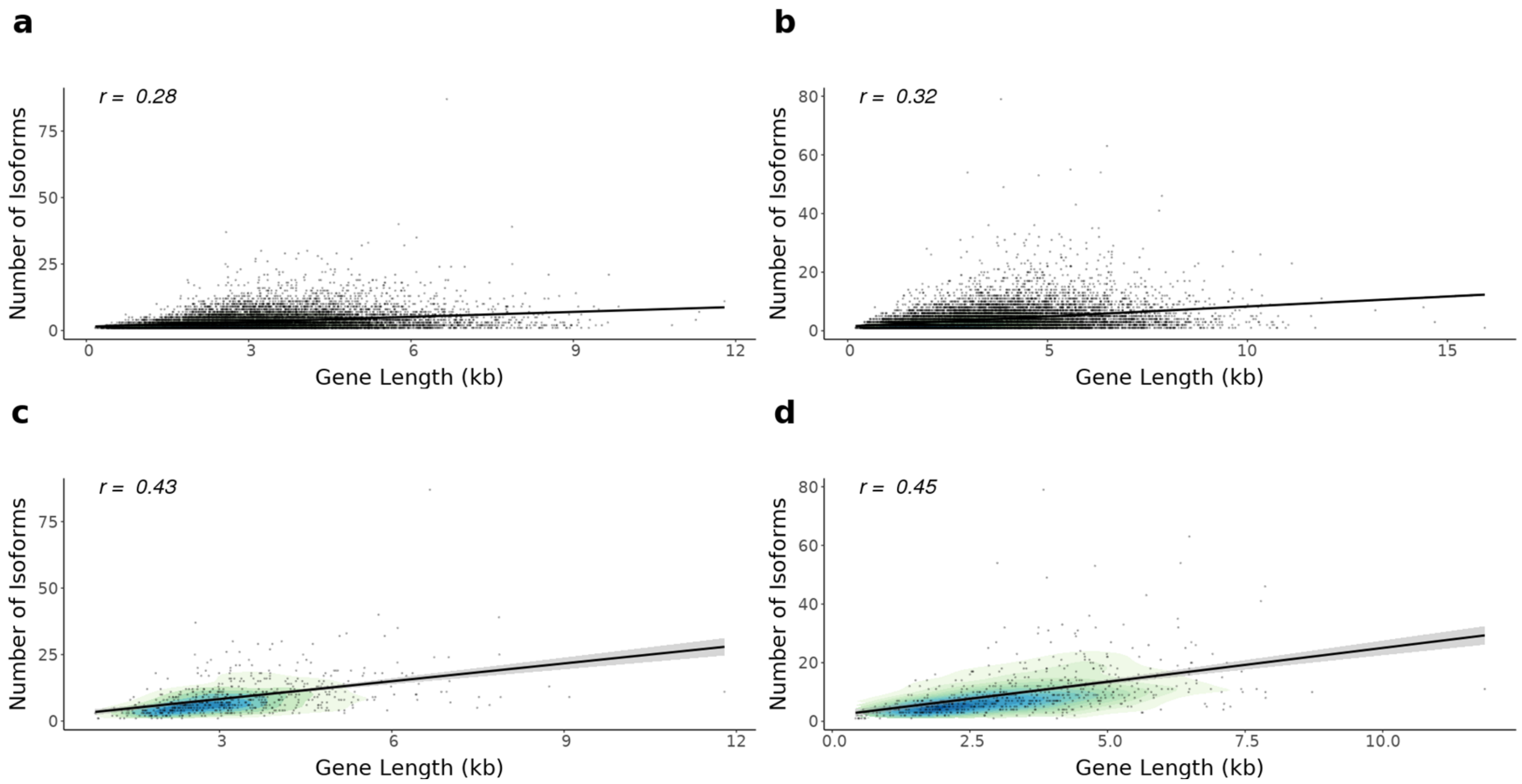

**Supplementary Figure 11: Genes with more exons were associated with more isoforms in human and mouse cortex.** The number of detected isoforms in **a)** human cortex (n = 7 biologically independent samples) and **b)** mouse cortex (n = 8 biologically independent samples) was correlated with the number of exons (human cortex: Pearson's correlation = 0.24, P = $2.42 \times 10^{-153}$; mouse cortex: Pearson's correlation = 0.22, P = $2.44 \times 10^{-143}$). A stronger relationship was observed among 'highly-expressed' genes (threshold of 2.5 $Log_{10}$ TPM) in both **c)** human cortex (Pearson's correlation = 0.42, P = $3.75 \times 10^{-25}$) and **d)** mouse cortex (Pearson's correlation = 0.43, P = $1.45 \times 10^{-28}$), consistent with the additional sensitivity of detecting isoforms in highly expressed genes. Density of genes is represented in increasing scale from light green to dark blue.

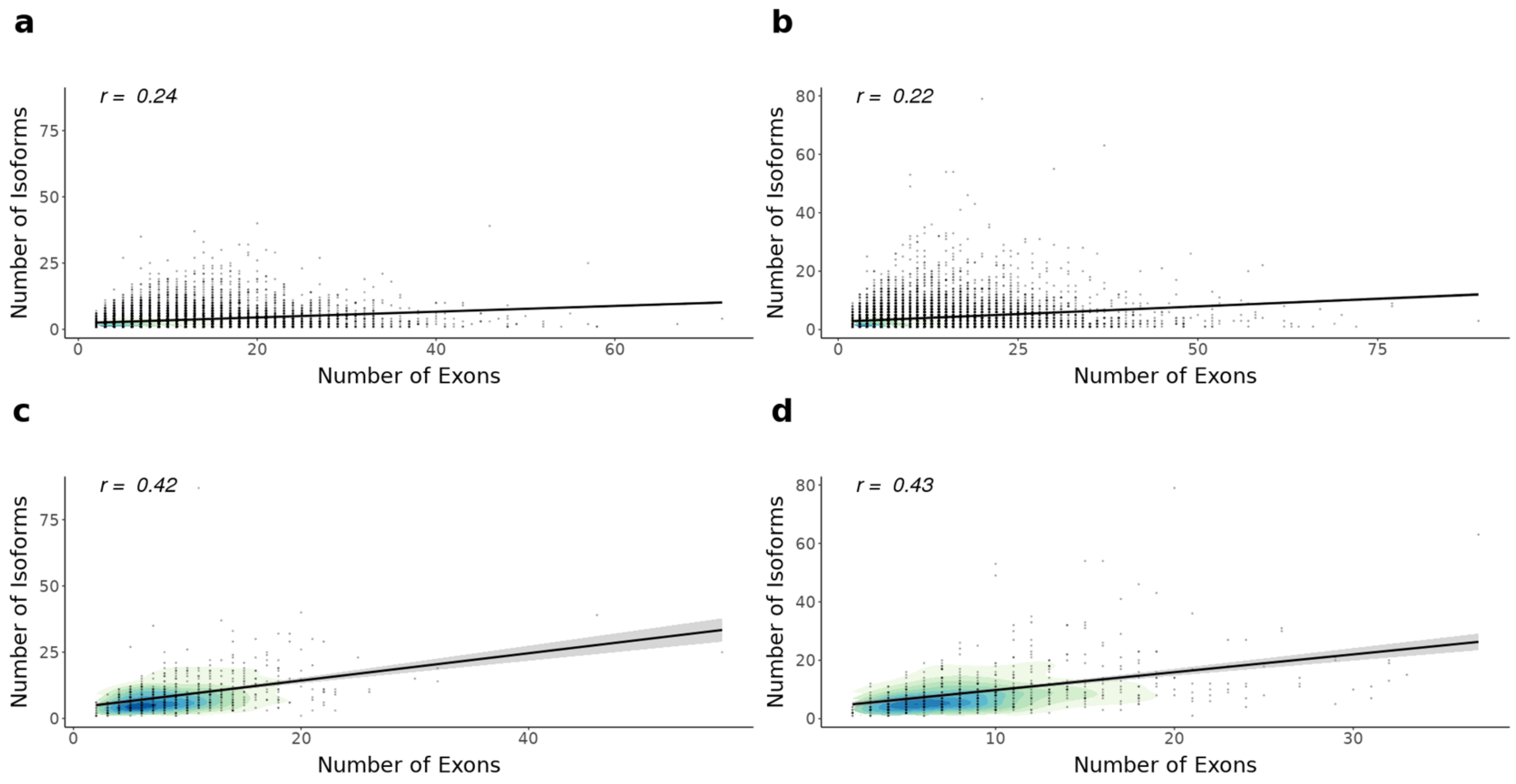

**Supplementary Figure 12: The majority of novel transcripts from annotated genes contain a combination of known donor or acceptor sites.** Shown is the proportion of novel transcripts of genes known in reference genome comparing **a)** human cortex (n = 7 biologically independent samples) and mouse cortex (n = 8 biologically independent samples), and **b)** human adult cortex (n = 4 biologically independent samples) and human fetal cortex (n = 3 biologically independent samples). An isoform was classified as novel it did not align with reference genome, and can be further subdivided into NIC if it contained a combination of known donor or acceptor sites, NNC if it contained at least one novel donor or acceptor site, genic/genomic if it overlapped with introns and exons, and fusion if it contained exons from two or more adjacent genes. Depictions of isoform classifications can be found in **Figure 1**. NIC – Novel In Catalogue, NNC – Novel Not in Catalogue.

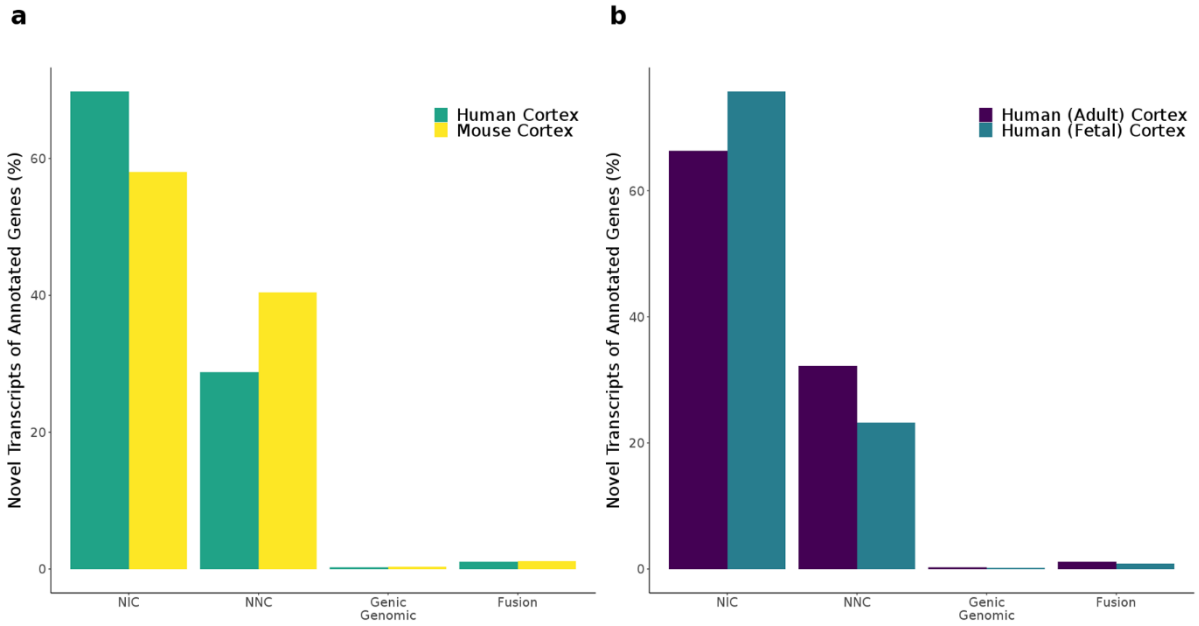

**Supplementary Figure 13: Novel transcripts from annotated genes were less expressed than already annotated transcripts.** Shown is the **a)** overall Iso-Seq transcript expression of novel and known transcripts and **b)** different RNA isoform categories of annotated genes in human cortex (n = 7 biologically independent samples) and mouse cortex (n = 8 biologically independent samples). Known transcripts were characterised by a higher expression in both human (Mann-Whitney-Wilcoxon test, $W = 2.59 \times 10^{8}$, $P < 2.23 \times 10^{-308}$) and mouse cortex (Mann-Whitney-Wilcoxon test, $W = 3.98 \times 10^{8}$, $P < 2.23 \times 10^{-308}$), primarily driven by FSM. FSM – Full Splice Match, ISM – Incomplete Splice Match, NIC – Novel In Catalogue, NNC – Novel Not in Catalogue.

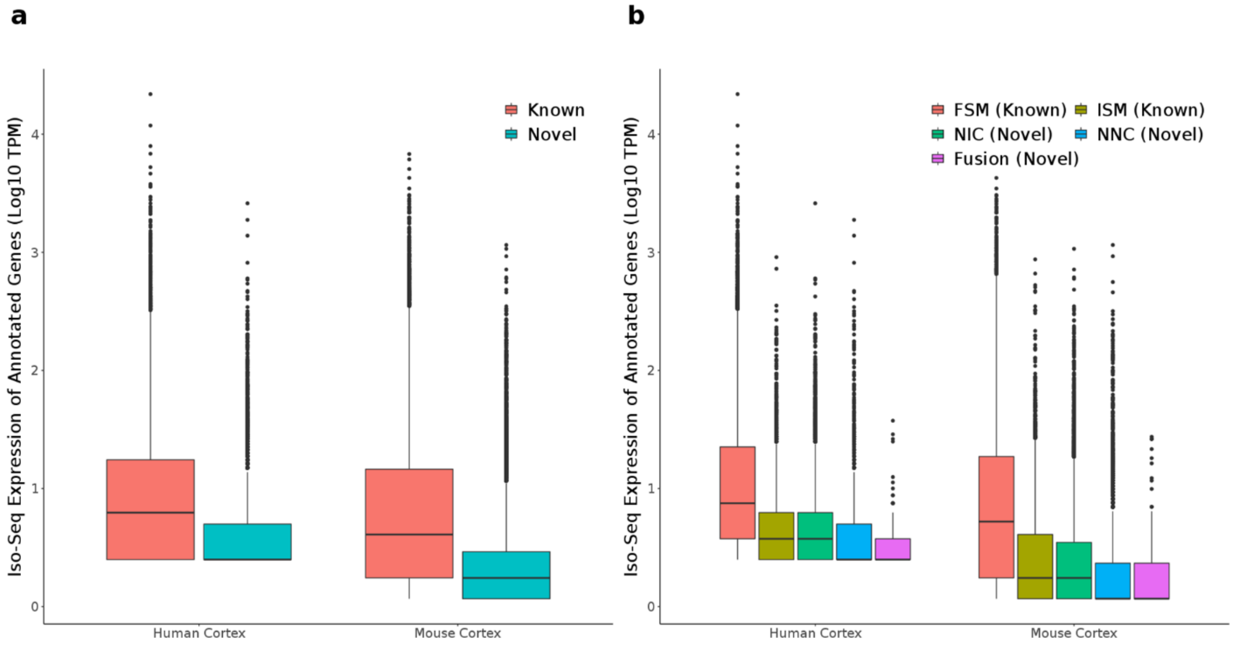

**Supplementary Figure 14: Novel transcripts were longer and had more exons than known transcripts.** Shown is the **a, b)** transcript length and **c, d)** number of exons for novel and known transcripts of annotated genes in human cortex (n = 7 biologically independent samples) and mouse cortex (n = 8 biologically independent samples). Novel transcripts were longer (human cortex: Mann-Whitney-Wilcoxon test, $W = 1.61 \times 10^{8}$, $P = 1.03 \times 10^{-133}$; mouse cortex: Mann-Whitney-Wilcoxon test, $W = 2.38 \times 10^{8}$, $P = 9.75 \times 10^{-203}$) and had more exons (human cortex: $W = 1.43 \times 10^{8}$, $P < 2.23 \times 10^{-308}$; mouse cortex: $W = 2.22 \times 10^{8}$, $P < 2.23 \times 10^{-308}$) than known transcripts, primarily driven by NIC. FSM – Full Splice Match, ISM – Incomplete Splice Match, NIC – Novel In Catalogue, NNC – Novel Not in Catalogue.

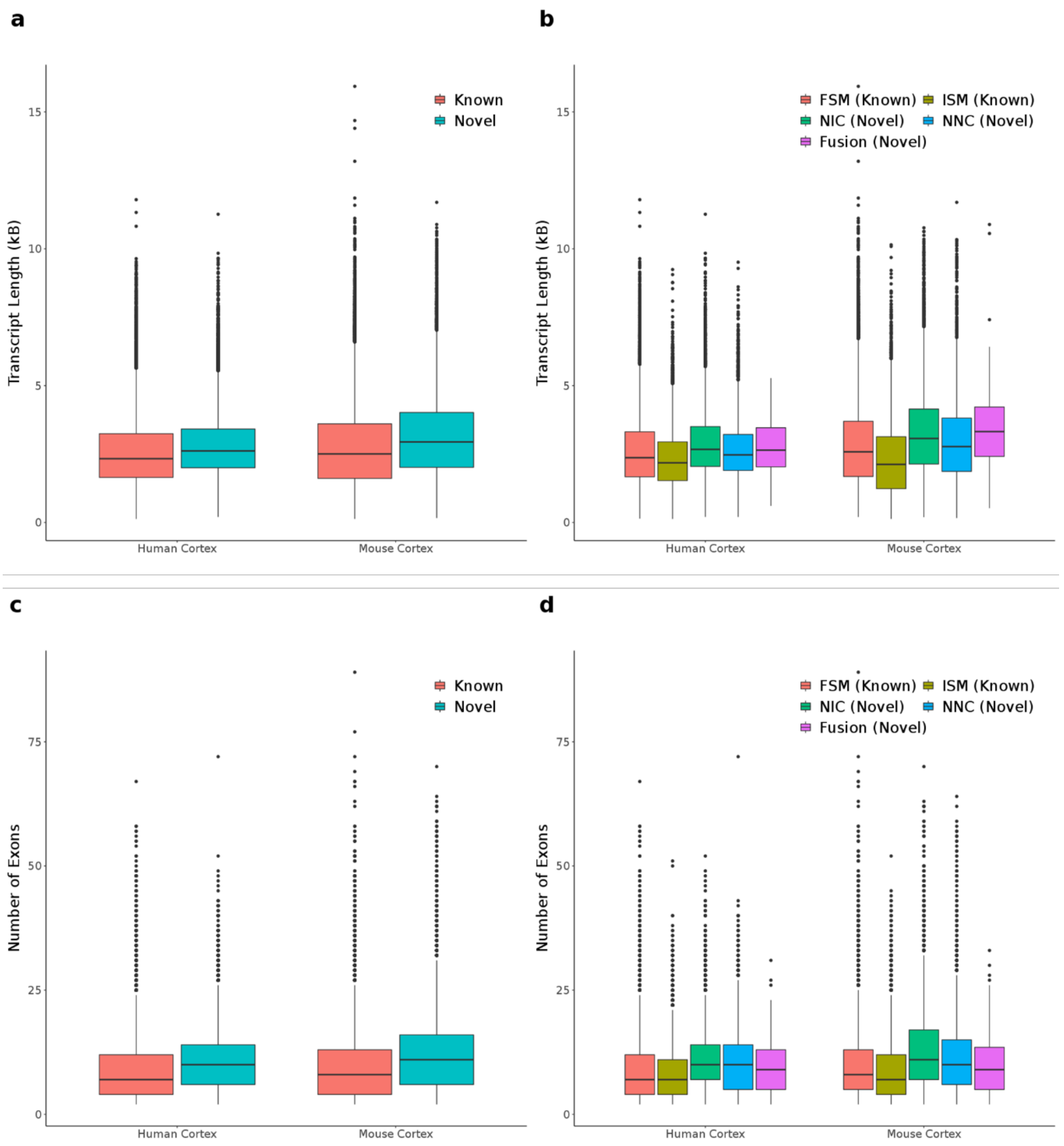

**Supplementary Figure 15: Majority of novel transcripts of annotated genes are located within 50bp of annotated CAGE peaks in the human and mouse cortex.** Shown is the location of novel transcripts relative to annotated CAGE peaks in the **a)** human (n = 7 biologically independent samples) and **b)** mouse (n = 8 biologically independent samples) cortex. Novel transcripts are classified as NIC, NNC, antisense, genic/genomic, and fusion. NIC – Novel In Catalogue, NNC – Novel Not in Catalogue.

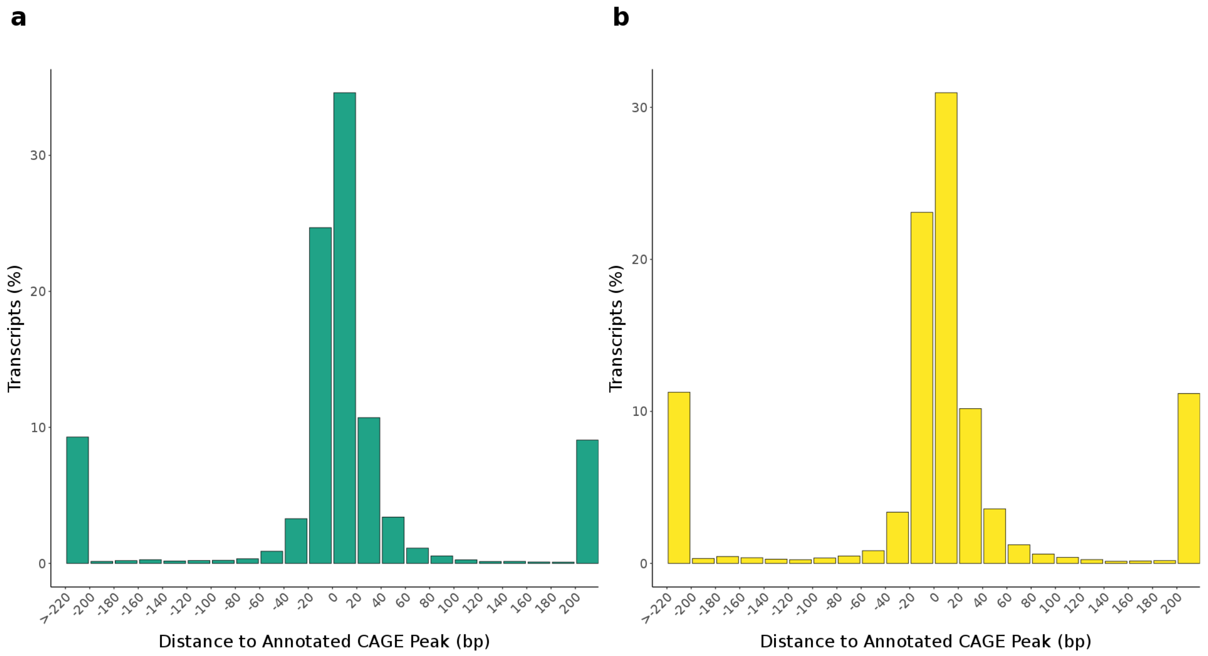

**Supplementary Figure 16: Majority of novel transcripts of annotated genes had full coverage of RNA-Seq across all splice junctions in human and mouse cortex.** Shown is the distribution of RNA-Seq coverage across splice junctions in novel transcripts of annotated genes in human fetal cortex (n = 3 biologically independent samples) and mouse cortex (n = 8 biologically independent samples). A junction coverage of 1 indicates RNA-Seq coverage across all splice junction of the transcript of interest.

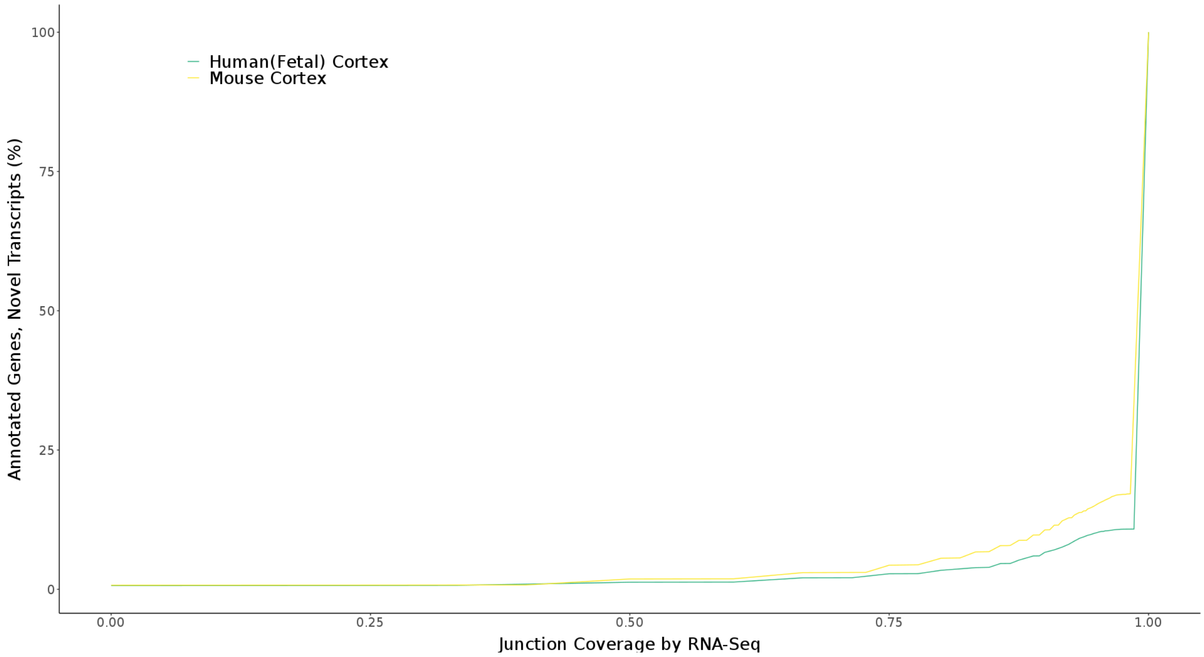

**Supplementary Figure 17: A large proportion of transcripts identified in human cortex were supported by nanopore sequencing**. Shown is the proportion of transcripts in human Iso-Seq cortex dataset (n = 7 biologically independent samples) that was also detected in human ONT cortex dataset (n = 2 biologically independent samples). 27,715 (65.0%) transcripts were also detected using nanopore sequencing (ONT), of which 7,081 (50.8%) transcripts were classified as novel and mapped to annotated genes. ONT – Oxford Nanopore Technology

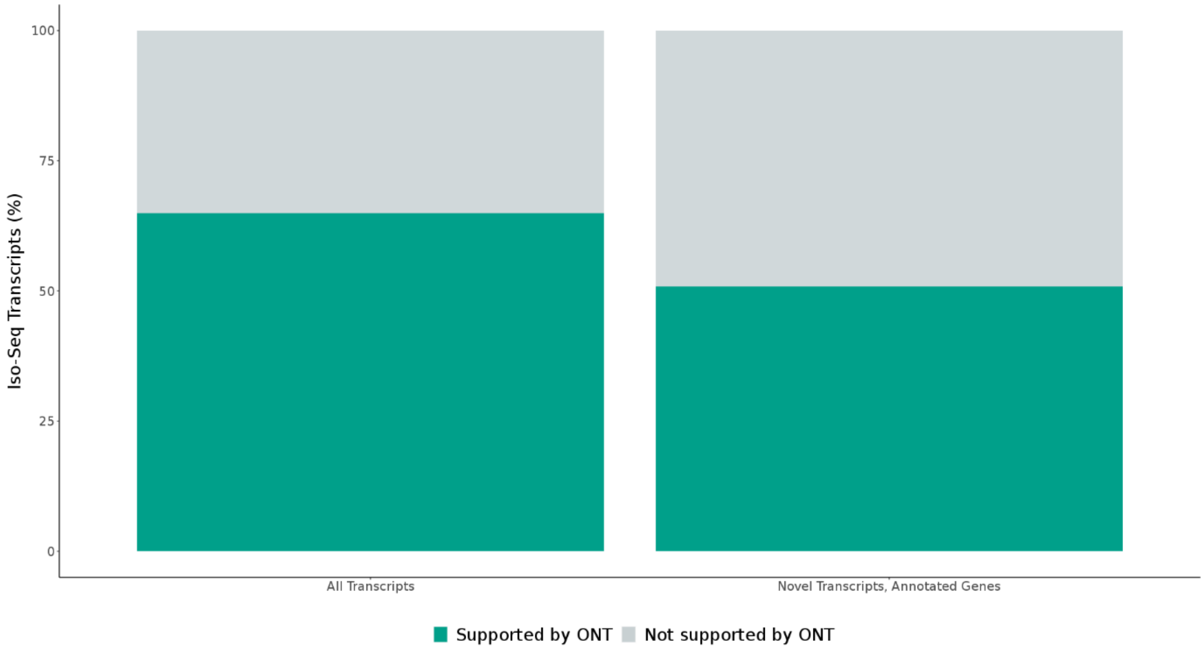

**Supplementary Figure 18: Example of a fusion transcript across three genes in the human cortex.** Shown is a UCSC genome browser track of a fusion transcript incorporating exons from three pseudogenes in the human cortex (n = 7 biologically independent samples), *AC138649.4_AC138649.1_PDCD6IPP1,* identified from GENCODE reference genome (hg38). Isoforms are coloured based on *SQANTI2* classification categories (blue = FSM, cyan = ISM, red = NIC, orange = NNC). FSM – Full Splice Match, ISM – Incomplete Splice Match, NIC – Novel In Catalogue, NNC – Novel Not in Catalogue.

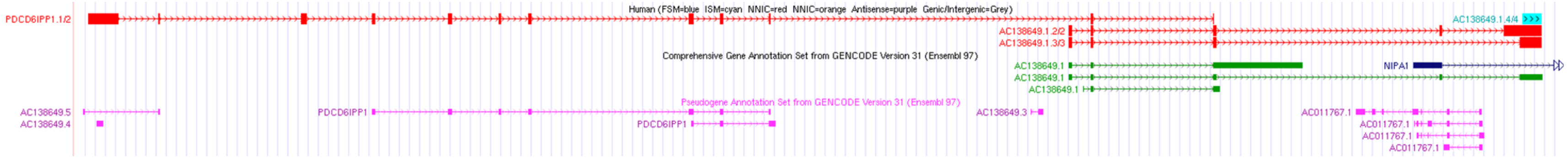

**Supplementary Figure 19: Novel genes were shorter and less abundant than annotated genes in human and mouse cortex.** Shown is the **a)** transcript length of multi-exonic transcripts and **b)** gene expression of annotated genes and novel genes in Iso-Seq dataset of human (n = 7 biologically independent samples) and mouse (n = 8 biologically independent samples) cortex. Transcripts from the same novel gene were identified by having identical first part of PacBio ID assigned from *IsoSeq 3.1.2* pipeline, and expression normalised to TPM was recalculated by discounting mono-exonic transcripts. Novel genes were shorter than annotated genes in human (Mann-Whitney-Wilcoxon test, $W = 1.6 \times 10^{6}$, $P = 1.5 \times 10^{-4}$) and mouse cortex (Mann-Whitney-Wilcoxon test, $W = 5.7 \times 10^{6}$, $P = 2.8 \times 10^{-25}$), and less abundant than annotated genes in human (Mann-Whitney-Wilcoxon test, $W = 5.1 \times 10^{5}$, $P = 9.8 \times 10^{-18}$) and mouse (Mann-Whitney-Wilcoxon test, $W = 1.5 \times 10^{6}$, $P = 7.7 \times 10^{-59}$). PacBio – Pacific Biosciences

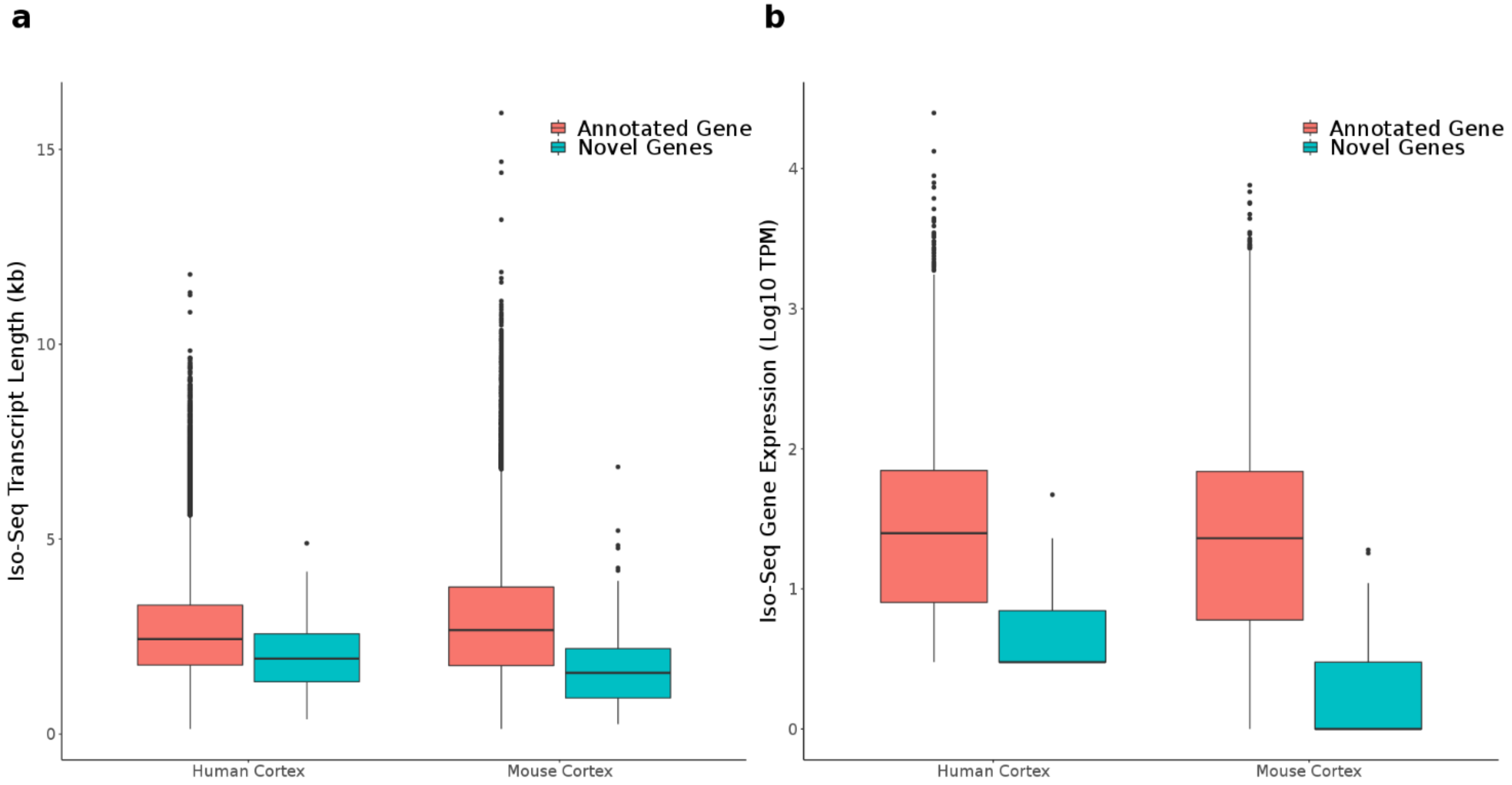

**Supplementary Figure 20: Example of an antisense novel gene that shares exons across two genes in the mouse cortex.** Shown is a UCSC genome browser track of an antisense novel gene (purple) overlapping *Serpinale* and *Serpina11*, complemented with GENCODE reference genome (mm10), and matched RNA-Seq data (n = 8 biologically independent samples).

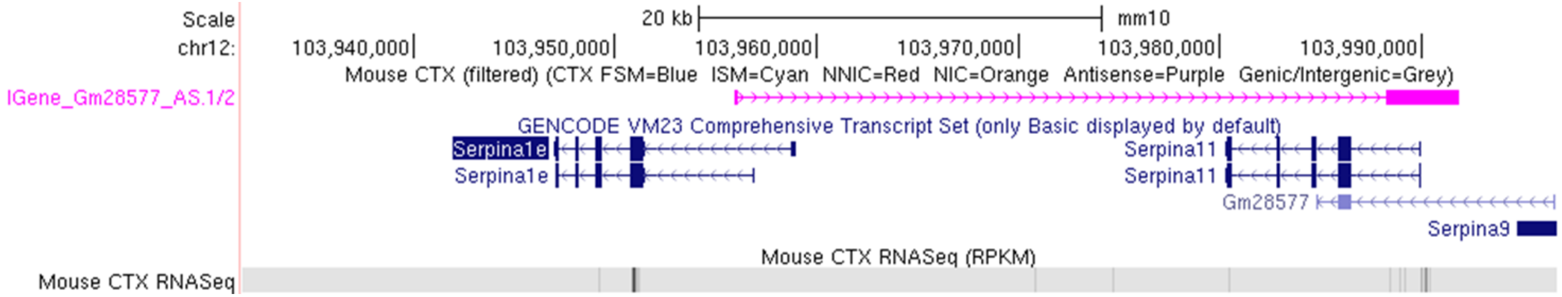

**Supplementary Figure 21: lncRNA transcripts were typically longer than non-lncRNA transcripts, despite containing fewer exons.** Shown is the distribution transcript length and exon number for lncRNA and non-lncRNA in **a, c)** human (n = 7 biologically independent samples) and **b, d)** mouse cortex (n = 8 biologically independent samples). LncRNA transcripts were found to be longer in both **a)** human cortex (Mann-Whitney-Wilcoxon, W = $3.84 \times 10^7$, P = $1.17 \times 10^{-45}$) and **b)** mouse cortex (Mann-Whitney-Wilcoxon test, W = $3.36 \times 10^7$, P = $2.83 \times 10^{-58}$), despite containing fewer exons in both **c)** human cortex (Mann-Whitney-Wilcoxon test, W = $5.57 \times 10^7$, P < $2.23 \times 10^{-308}$) and **d)** mouse cortex (Mann-Whitney-Wilcoxon test, W = $4.54 \times 10^7$, P < $2.23 \times 10^{-308}$). lncRNA – long non-coding RNA.

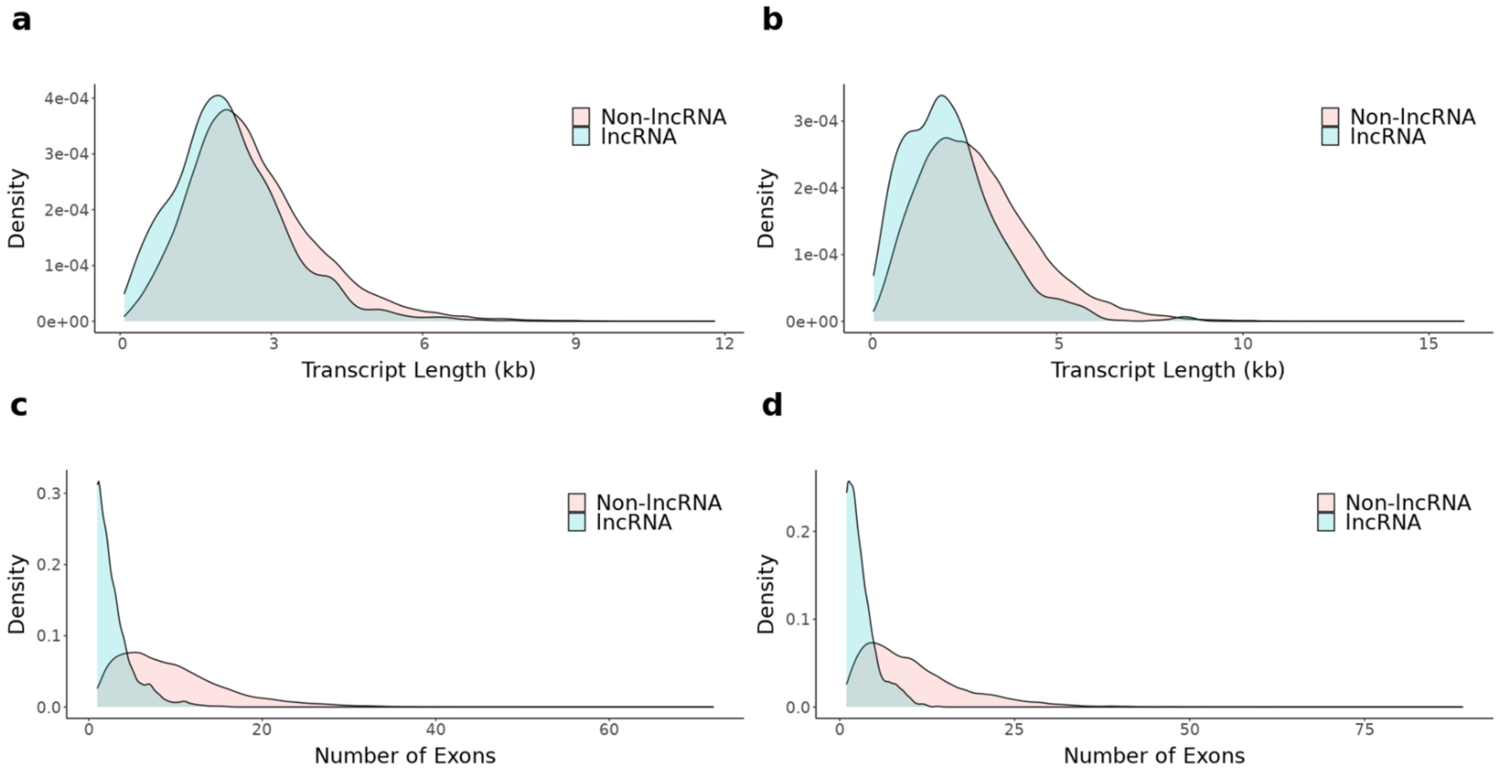

**Supplementary Figure 22: LncRNA were characterized by lower transcript expression and fewer isoforms than non-lncRNA.** Shown is the transcript expression and isoform diversity for lncRNA and non-lncRNA genes in **a, c)** human (n = 7 biologically independent samples) and **b, d)** mouse cortex (n = 8 biologically independent samples). LncRNA isoforms were characterized by lower transcript expression than non-lncRNA transcripts in both **a)** human (Mann-Whitney-Wilcoxon test, W = $3.70 \times 10^{7}$, P = $1.12 \times 10^{-30}$) and **b)** mouse cortex (Mann-Whitney-Wilcoxon test, W = $3.28 \times 10^{7}$, P = $4.13 \times 10^{-49}$), and lower isoform diversity in **c)** human (Mann-Whitney-Wilcoxon test, W = $7.11 \times 10^{6}$ , P = $1.47 \times 10^{-99}$) and **d)** mouse cortex (Mann-Whitney-Wilcoxon test, W = $5.71 \times 10^{6}$, P = $2.09 \times 10^{-101}$). Lnc-RNA – long non-coding RNA.

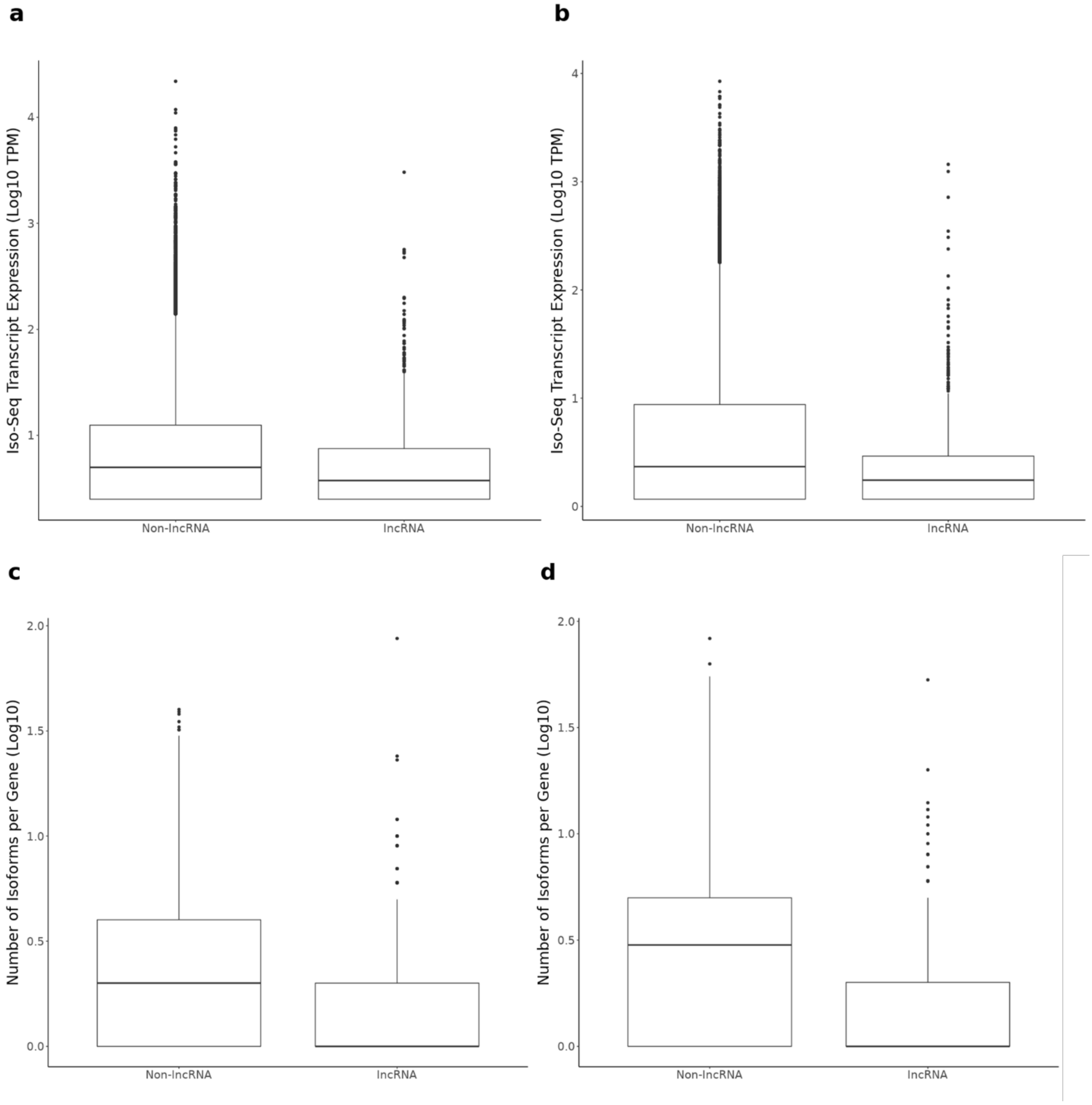

**Supplementary Figure 23: LncRNA transcripts with potential protein coding domains were identified in both the human and mouse cortex, with shorter open reading frames (ORFs) than non-lncRNA.** Shown is the distribution of ORF length for lncRNA and non-lncRNA transcripts in the **a)** human (n = 7 biologically independent samples) and **b)** mouse cortex (n = 8 biologically independent samples). ORF in lncRNA transcripts is generally shorter than ORF in non-lncRNA transcripts in both human (Mann-Whitney-Wilcoxon test, $W = 2.11 \times 10^{7}$, $P = 1.01 \times 10^{-254}$) and mouse (Mann-Whitney-Wilcoxon test, $W = 1.79 \times 10^{7}$, $P = 9.50 \times 10^{-186}$). ORF – Open reading frame, LncRNA – long non-coding RNA.

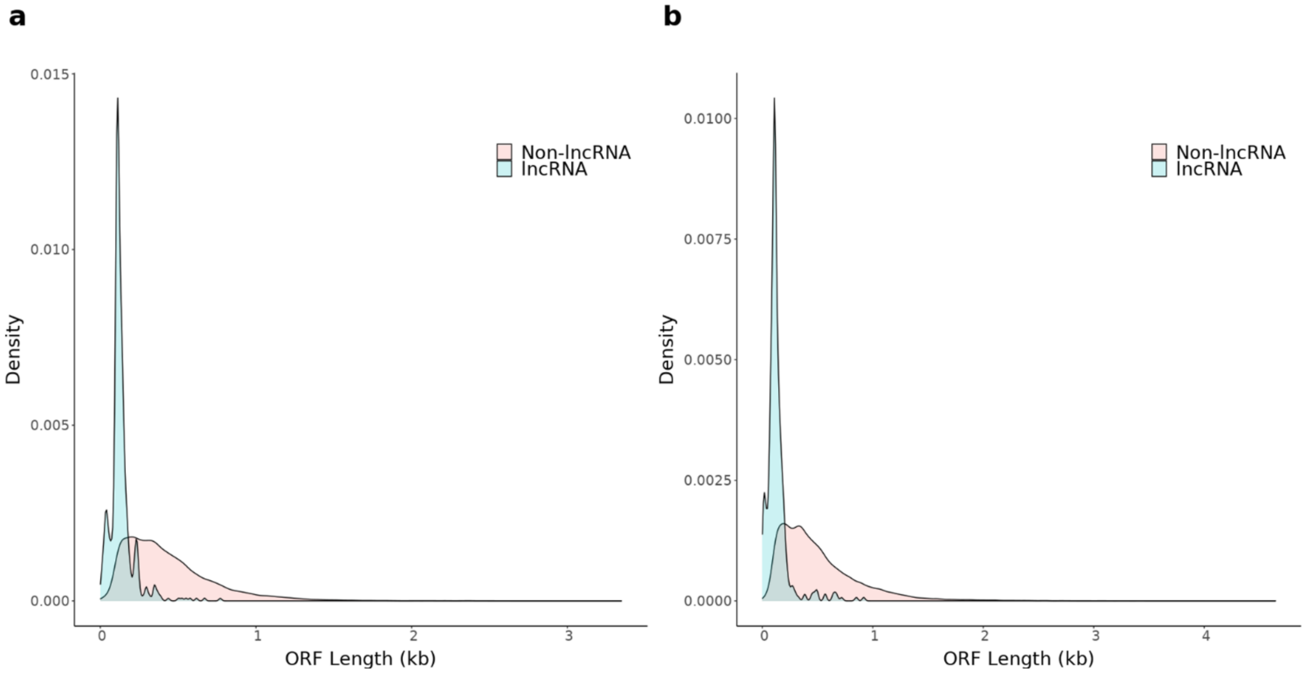

**Supplementary Figure 24: Alternative first exon and alternative last exon are the most prominent AS events in human and mouse cortex.** Shown is the proportion of alternative splicing events in human adult cortex (n = 4 biologically independent samples) and human fetal cortex (n = 3 biologically independent samples). MX and SE events were determined using *SUPPA2*, IR with *SQANTI2* and A3', A5', AF and AL with custom scripts. AF – Alternative First Exon, AL – Alternative Last Exon, A5' – Alternative 5' prime, A3' – Alternative 3' prime, IR – Intron Retention, MX – Mutually Exclusive, SE – Skipped Exon.

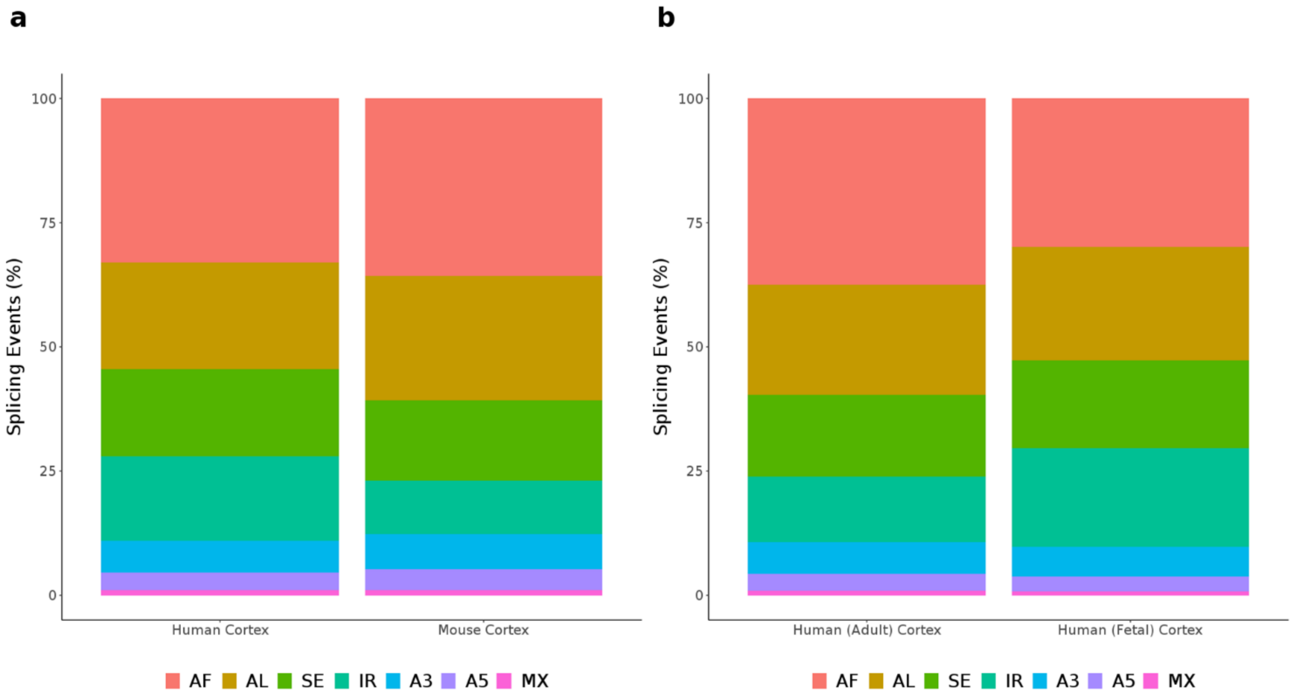

**Supplementary Figure 25: There was a large overlap of genes influenced by specific AS events in human and mouse cortex.** Shown is the number of genes observed with the different AS events (A3, A5, AF, AL, IR, MX, and SE) common and unique to **a)** human (n = 7 biologically independent samples) and mouse cortex (n = 8 biologically independent samples) and **b)** human adult (n = 4 biologically independent samples) and human fetal (n = 3 biologically independent samples) cortex. MX and SE events were determined using *SUPPA2*, IR with *SQANTI2* and A3', A5', AF and AL with customised scripts. AF – Alternative First Exon, AL – Alternative Last Exon, A5' – Alternative 5' prime, A3' – Alternative 3' prime, IR – Intron Retention, MX – Mutually Exclusive, SE – Skipped Exon.

**a**

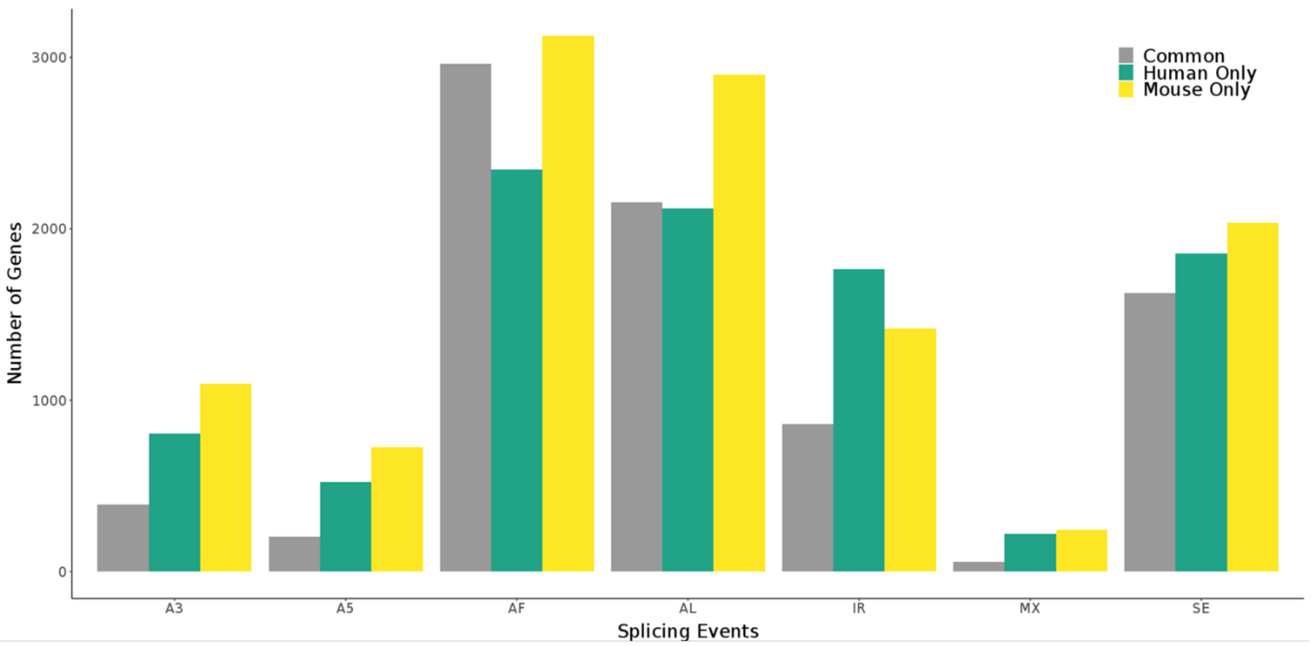


**b**

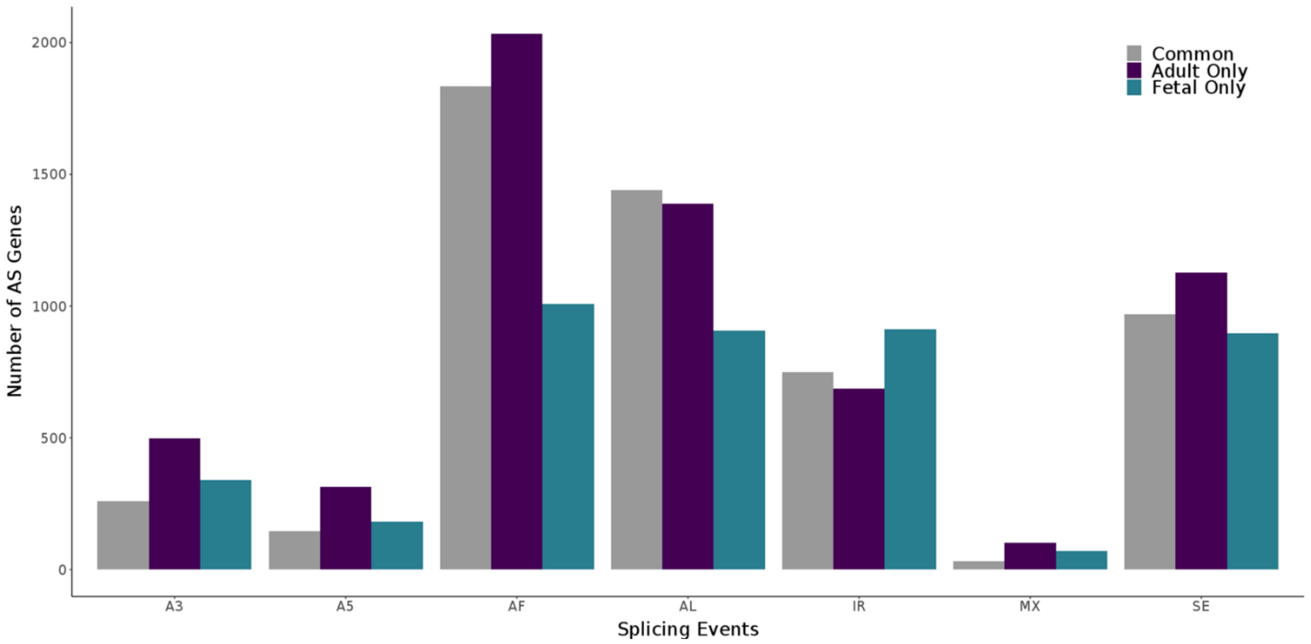

**Supplementary Figure 26: The majority of genes were characterized by a single AS event in both fetal and adult cortex.** Shown is the number of splicing events across genes identified with alternative splicing events in **a)** human cortex (n = 7 biologically independent samples) and mouse cortex (n = 8 biologically independent samples), and **b)** human adult cortex (n = 4 biologically independent samples) and human fetal cortex (n = 3 biologically independent samples), with the majority reporting one AS event. A small number of genes were characterised with more than 7 AS events, the maximum possible number of AS events (AF, AL, A5', A3', IR, MX, SE) that can be identified from this analysis. AS – Alternative Splicing, AF – Alternative First Exon, AL – Alternative Last Exon, A5' – Alternative 5' prime, A3' – Alternative 3' prime, IR – Intron Retention, MX – Mutually Exclusive, SE – Skipped Exon.

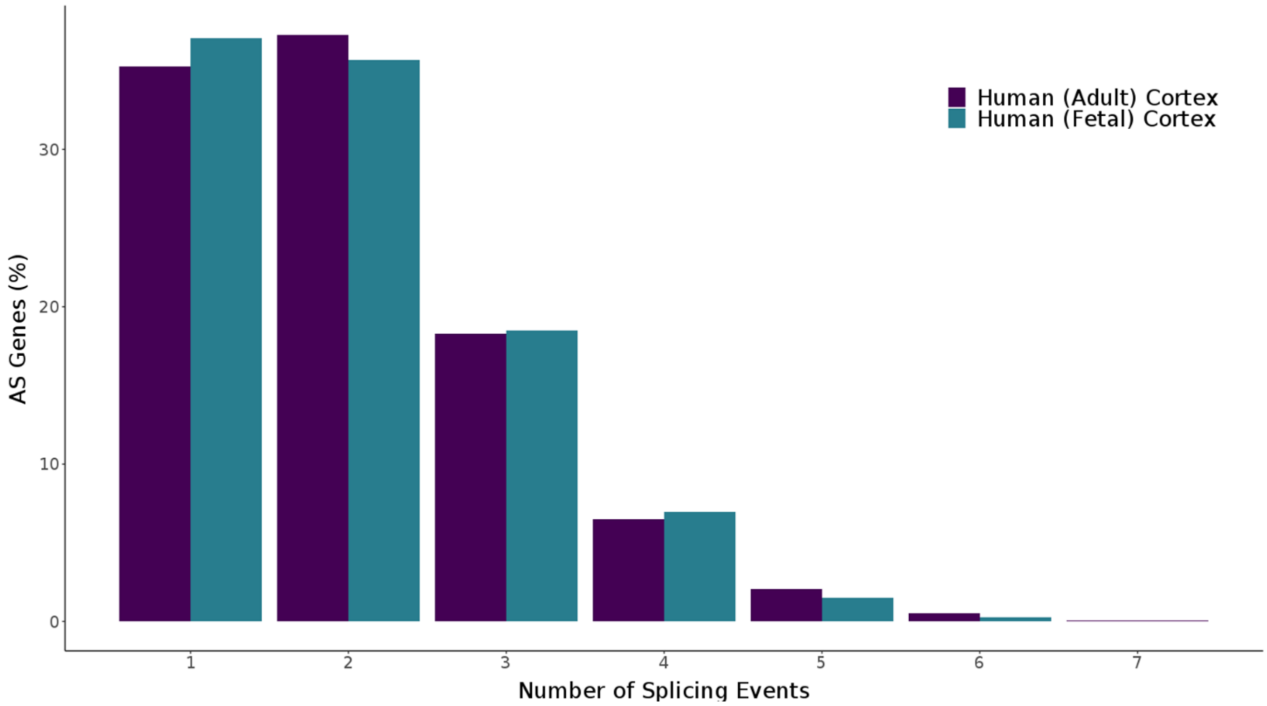

**Supplementary Figure 27: Alternative first (AF) is the most prevalent AS event in human and mouse cortex.** Shown is a UCSC genome track of *CELF2* in human cortex (n = 7 biologically independent samples), highlighted with a number of transcripts characterised by AF. The differing lengths of first exon across these transcripts corresponded with RNA-Seq coverage in human fetal cortex (n = 3 biologically independent samples). Isoforms are coloured based on *SQANTI2* classification categories (blue = FSM, cyan = ISM, red = NIC, orange = NNC). AF – Alternative First, AS – Alternative Splicing, FSM – Full splice match, ISM – Incomplete splice match, NIC – Novel in catalogue, NNC – novel not in catalogue.

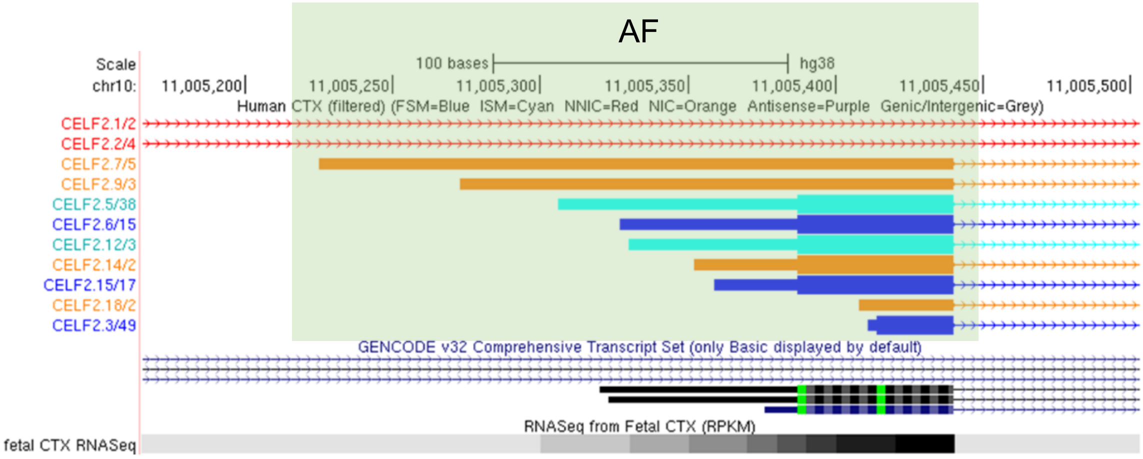

**Supplementary Figure 28: IR-transcripts predicted for NMD were more lowly expressed than non-NMD-transcripts.** Shown is the Iso-Seq transcript expression in **a)** human cortex (n = 7 biologically independent samples) and **b)** mouse cortex (n = 8 biologically independent samples) of intron-retained (IR) transcripts predicted for nonsense-mediated decay (IR-NMD), IR transcripts, NMD transcripts, and transcripts not characterised by either IR or NMD (Non IR-NMD). As shown, IR-NMD transcripts were particularly lowly expressed (human cortex: Mann-Whitney-Wilcoxon test, W = $6.00 \times 10^{6}$, P = $4.43 \times 10^{-14}$; mouse cortex: Mann-Whitney-Wilcoxon test, W = $3.40 \times 10^{6}$, P = $1.21 \times 10^{-35}$). IR – Intron Retention, NMD – Nonsense-mediated mRNA decay

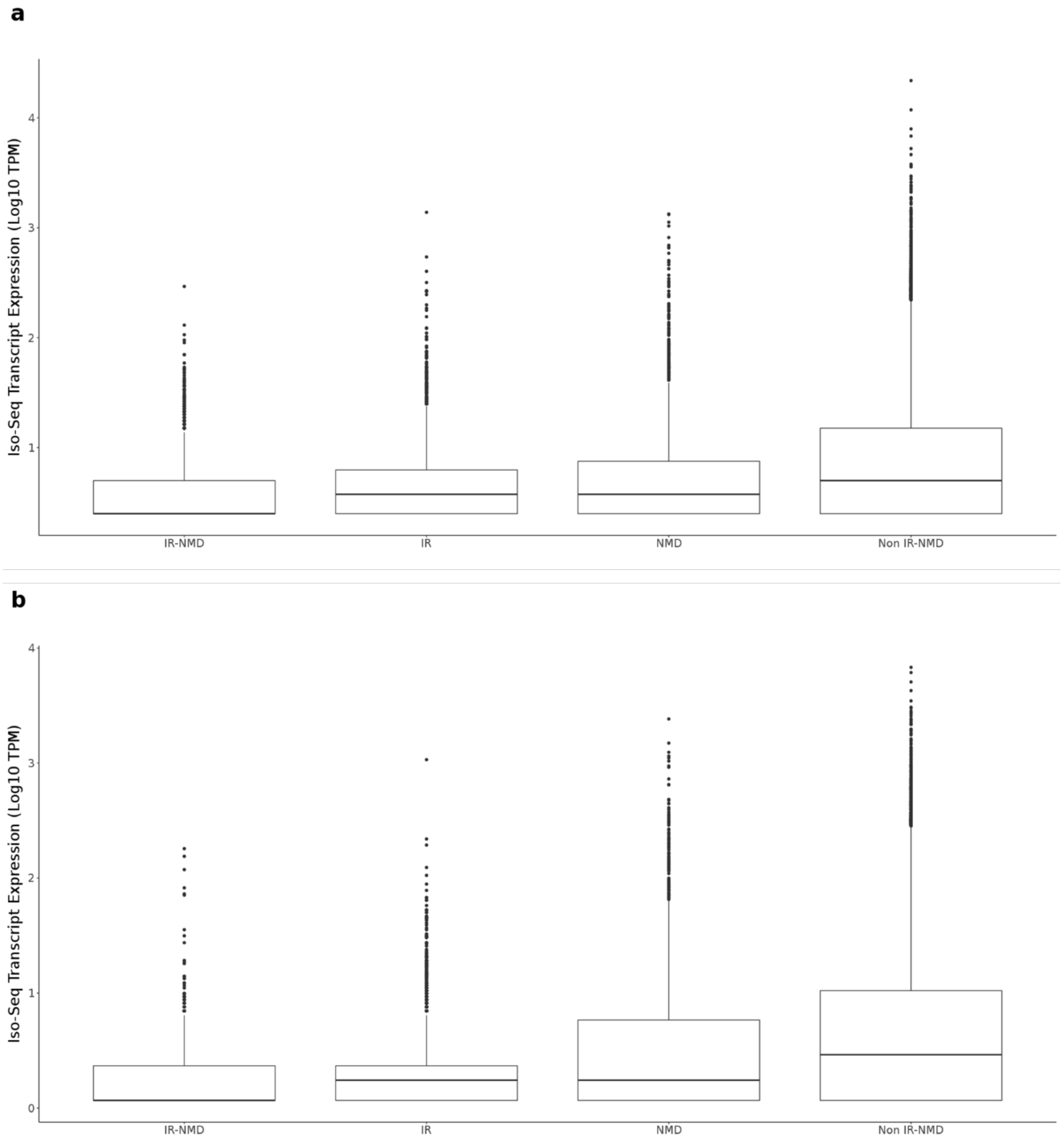

**Supplementary Figure 29: NMD was found to be particularly enriched amongst transcripts with intron retention.** Shown is the overlap of genes with IR-transcripts, NMD-transcripts, and transcripts with both IR and NMD in **a)** human cortex, **b)** human adult cortex (n = 4 biologically independent samples), **c)** human fetal cortex (n = 3 biologically independent samples) and **d)** mouse cortex (n = 8 biologically independent samples). Of note, genes containing both IR and NMD transcripts were further classified into genes that contain transcripts that were *both* IR and NMD (purple) and genes that contain transcripts where IR and NMD were mutually exclusive (dark orange). As such in **a)** human cortex, 1150 genes were associated with IR-transcripts that were predicted for NMD, and 168 genes that contained IR-transcripts and NMD-transcripts. IR – Intron retention, NMD – Nonsense-mediated mRNA decay.

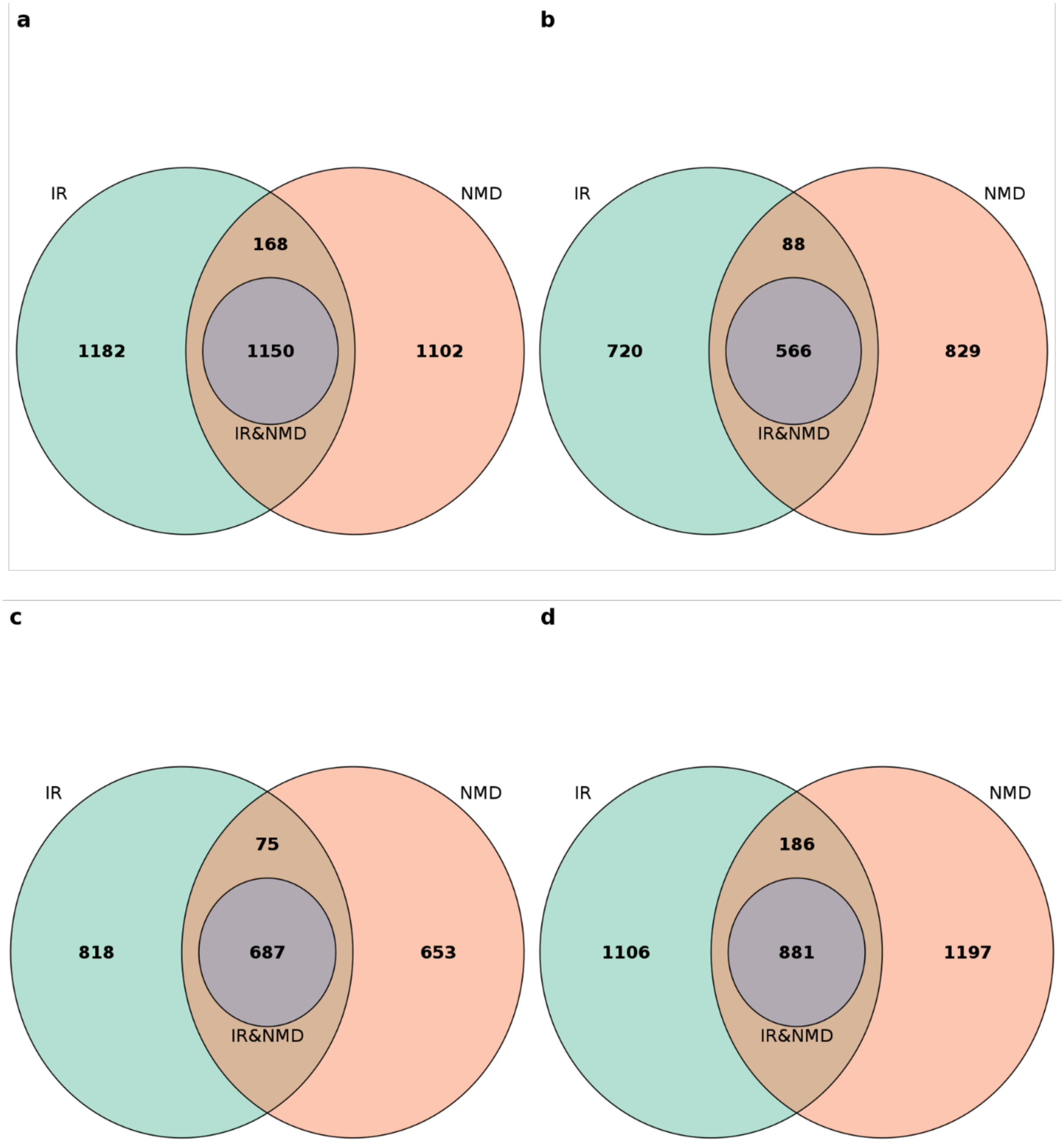

**Supplementary Figure 30: The number of unique isoforms for commonly expressed genes is correlated between human and mouse.** Shown is the relationship between the number of multi-exonic isoforms in **a)** human and mouse cortex (n = 8 biologically independent samples), and **b)** after applying gene expression threshold, and **c)** human adult (n = 4 biologically independent samples) and human fetal (n = 3 biologically independent samples) cortex and **d)** after applying gene expression threshold. A relationship was observed between human and mouse (Pearson's correlation = 0.52, $P < 2.23 \times 10^{-308}$), human adult and human fetal cortex (Pearson's correlation = 0.56, $P < 2.23 \times 10^{-308}$). A gene expression threshold of 2.5 $Log_{10}$ TPM was applied to all cortical datasets, and a stronger relationship was observed between human and mouse cortex (Pearson's correlation = 0.73, $P = 1.28 \times 10^{-48}$), and between human adult and human fetal cortex (Pearson's correlation = 0.72, $P = 1.26 \times 10^{-41}$). Of note, homology was considered for human and mouse comparison (see **Online Methods**).

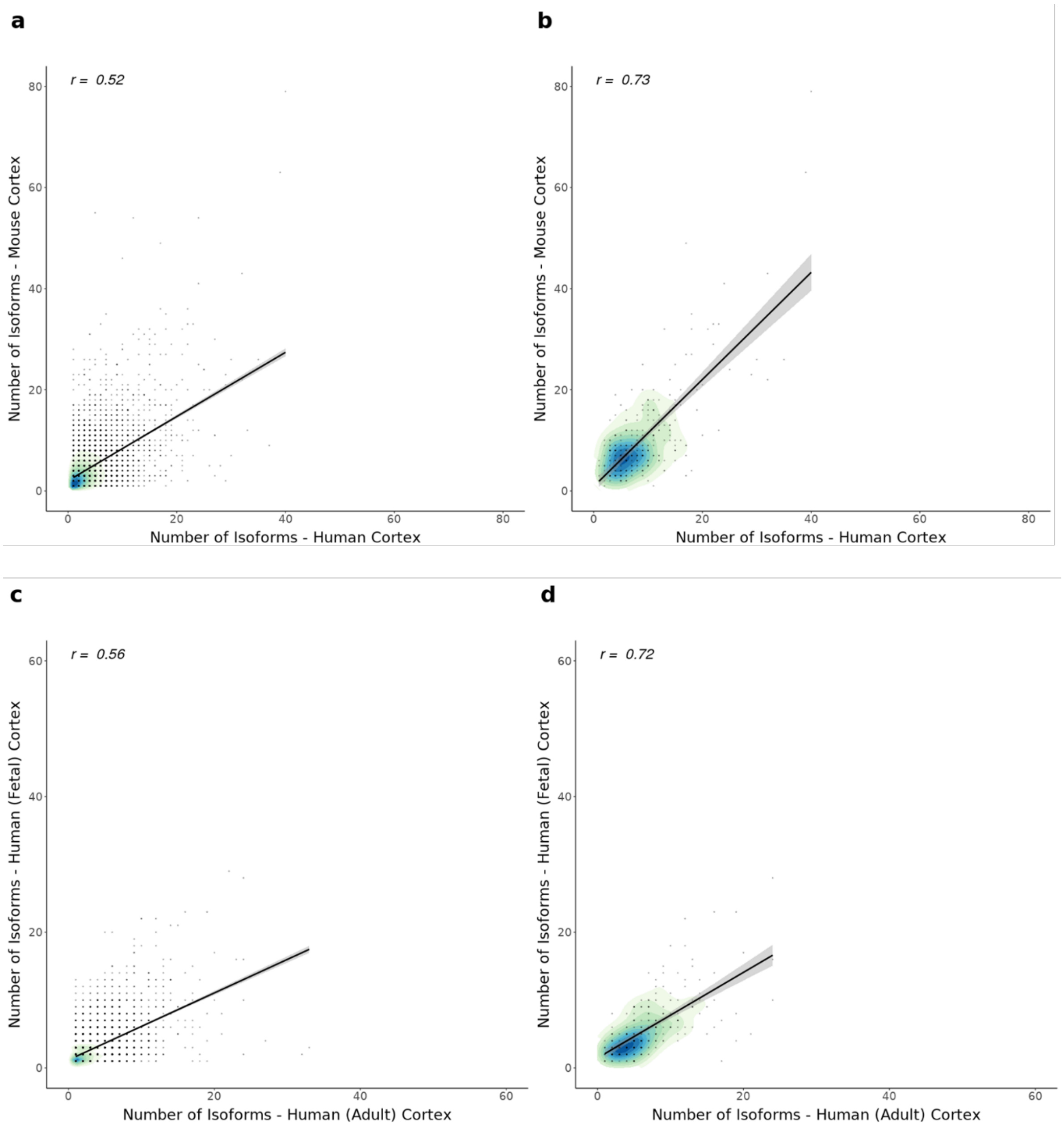

**Supplementary Figure 31: Notable difference in isoform number in gene encoding *SORBS1* in human and mouse cortex.** Shown is UCSC genome browser track of *SORBS1* gene in **a)** human cortex (n = 5 multi-exonic isoforms) and in **b)** mouse cortex (n = 55 multi-exonic isoforms), with notable difference in isoform diversity. Isoforms are coloured based on *SQANTI2* classification categories (blue = FSM, cyan = ISM, red = NIC, orange = NNC). FSM – Full Splice Match, ISM – Incomplete Splice Match, NIC – Novel In Catalogue, NNC – Novel Not in Catalogue

a

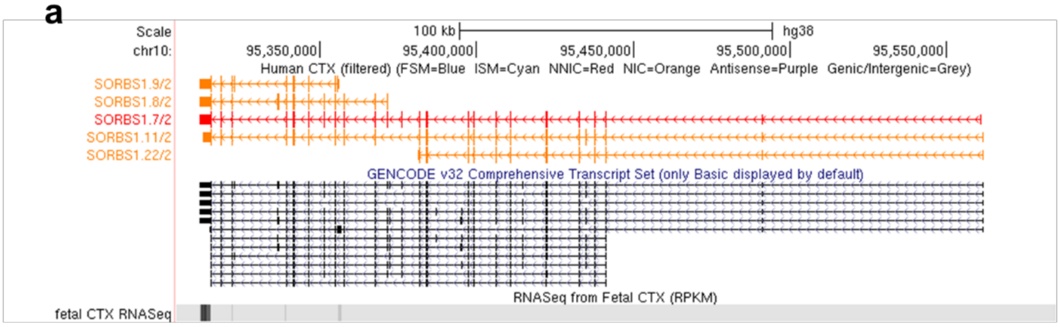


b

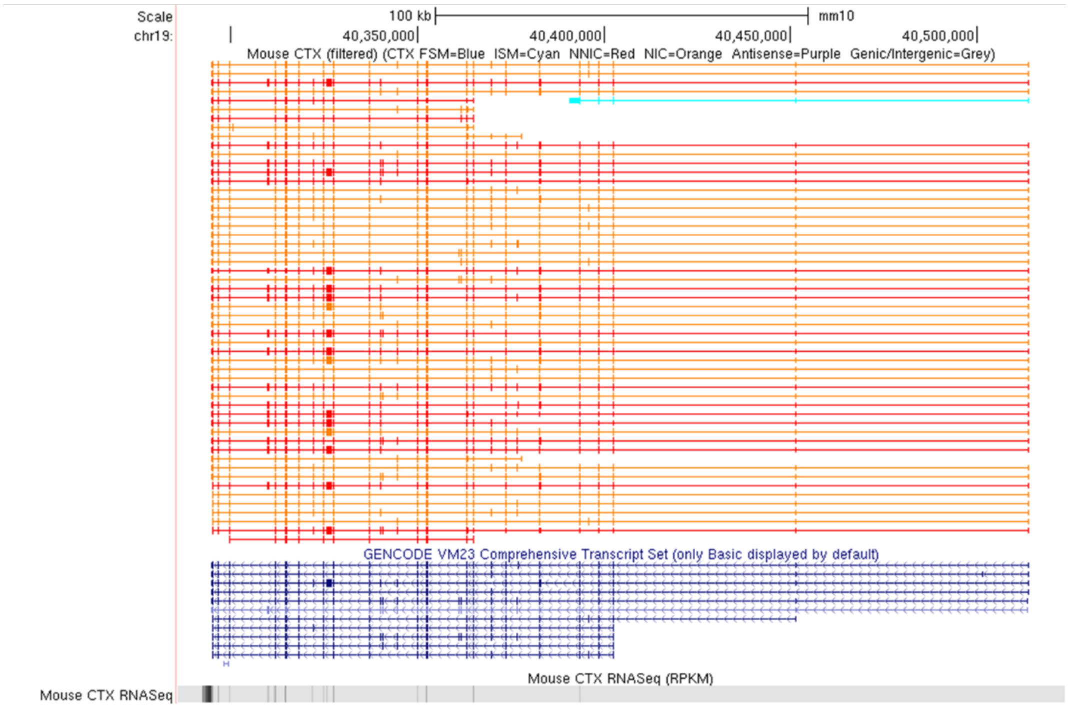

**Supplementary Figure 32: Notable difference in isoform number in gene encoding *DLGAP1* in human and mouse cortex.** Shown is UCSC genome browser track of *DLGAP1* gene in **a)** human cortex (n = 12 multi-exonic isoforms) and in **b)** mouse cortex (n = 54 multi-exonic isoforms), with notable difference in isoform diversity. Isoforms are coloured based on *SQANTI2* classification categories (blue = FSM, cyan = ISM, red = NIC, orange = NNC). FSM – Full Splice Match, ISM – Incomplete Splice Match, NIC – Novel In Catalogue, NNC – Novel Not in Catalogue

**a**

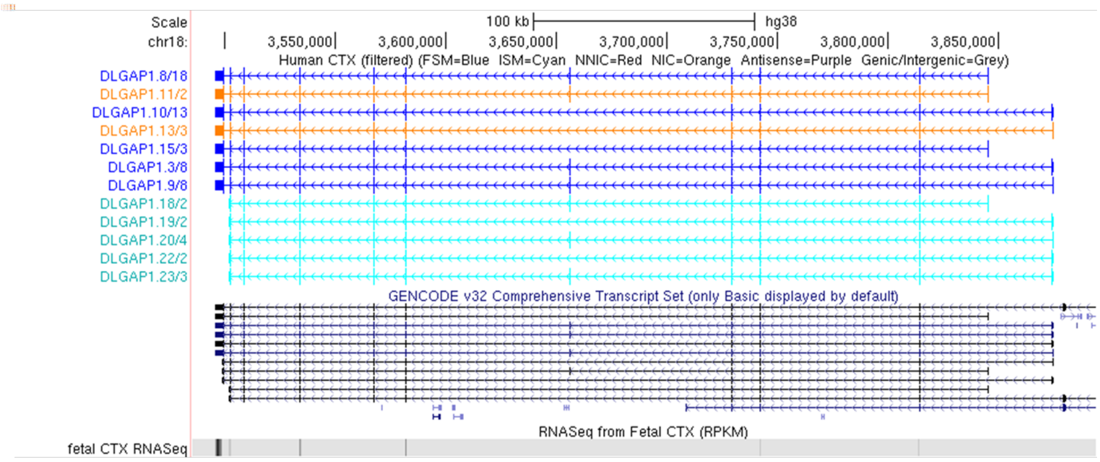


**b**

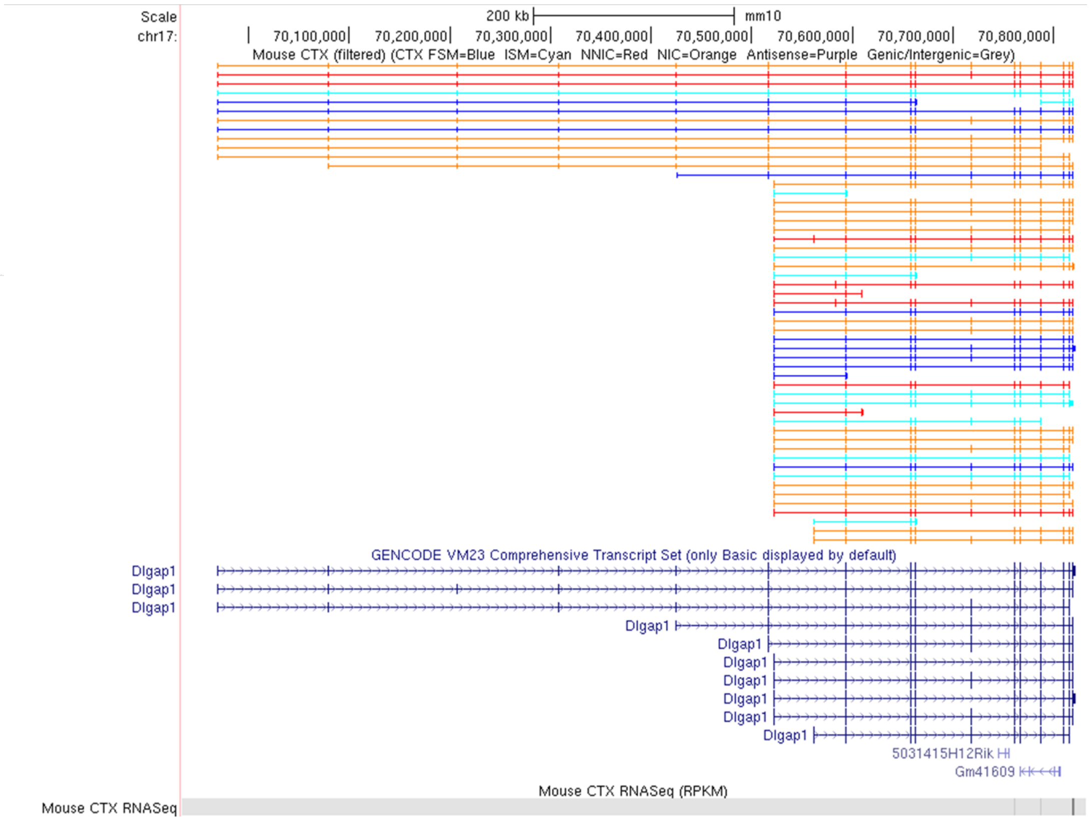

**Supplementary Figure 33: Notable difference in isoform number in gene encoding *NDUFS2* in human and mouse cortex.** Shown is UCSC genome browser track of *NDFUS2* gene in **a)** human cortex (n = 18 multi-exonic isoforms) and in **b)** mouse cortex (n = 1 multi-exonic isoform), with notable difference in isoform diversity. Isoforms are coloured based on *SQANTI2* classification categories (blue = FSM, cyan = ISM, red = NIC, orange = NNC). FSM – Full Splice Match, ISM – Incomplete Splice Match, NIC – Novel In Catalogue, NNC – Novel Not in Catalogue

**a**

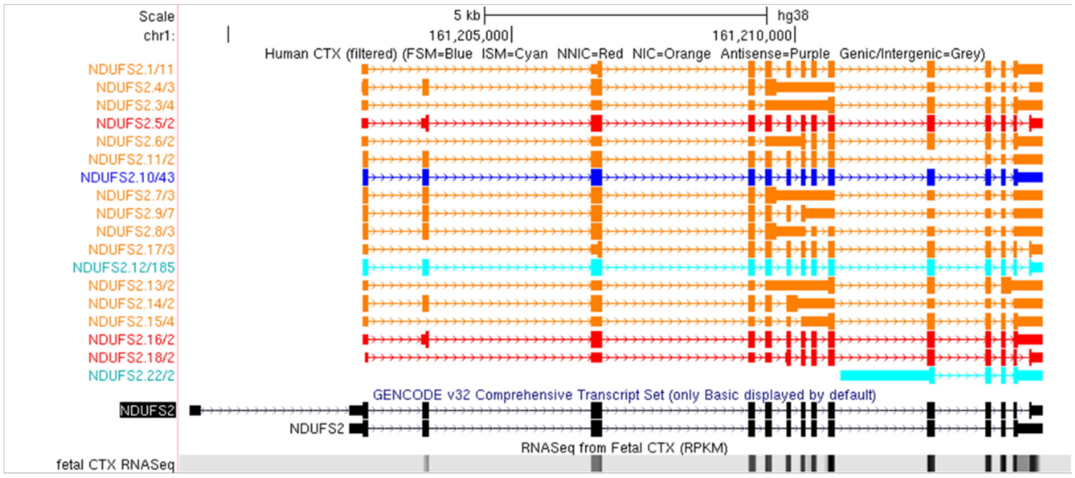


**b**

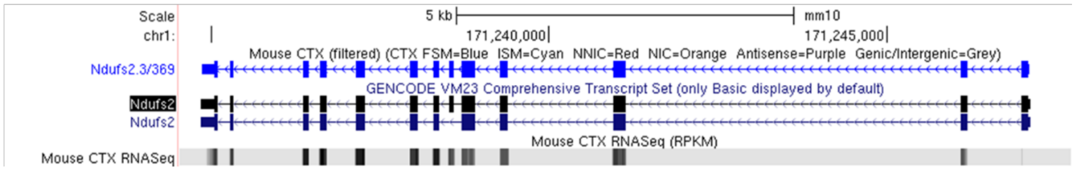

**Supplementary Figure 34: Notable difference in isoform number in gene encoding TMEM191C in human and mouse cortex.** Shown is UCSC genome browser track of *TMEM191C* gene in **a)** human cortex (n = 1 multi-exonic isoform) and in **b)** mouse cortex (n = 28 multi-exonic isoforms), with notable difference in isoform diversity. Isoforms are coloured based on *SQANTI2* classification categories (blue = FSM, cyan = ISM, red = NIC, orange = NNC). FSM – Full Splice Match, ISM – Incomplete Splice Match, NIC – Novel In Catalogue, NNC – Novel Not in Catalogue

**a**

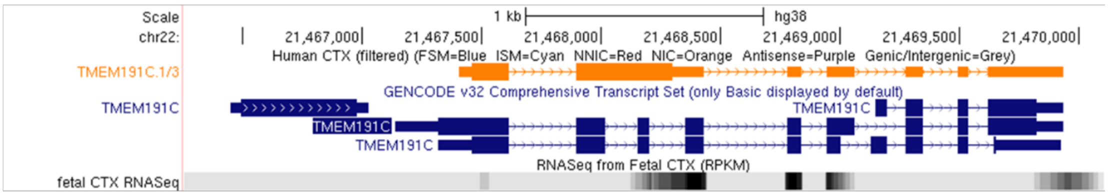


**b**

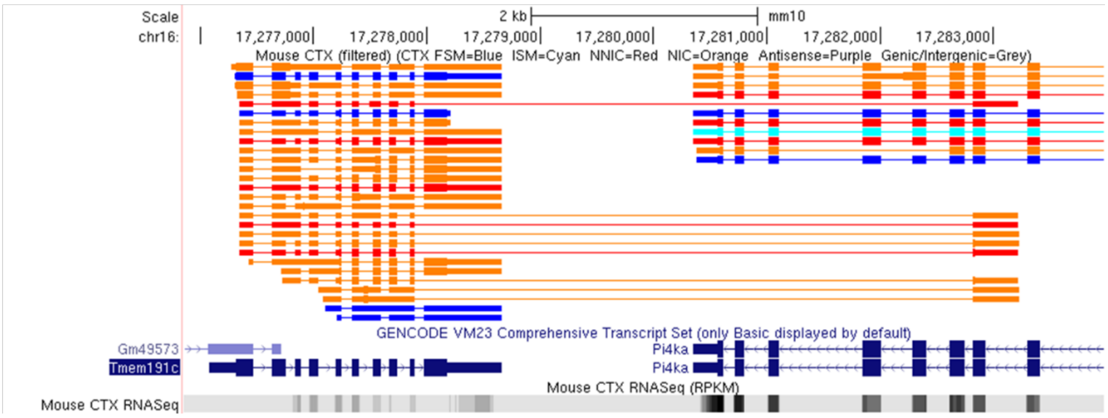

**Supplementary Figure 35: Majority of splice junctions in transcripts associated with annotated genes are canonical in human and mouse cortex, with greater splicing junction diversity observed in mouse.** Shown is the **a)** number of different splice junctions identified from transcripts mapped to annotated genes, classified by canonical (GT/AG, GC/AG, ATAC) and non-canonical according to *SQANTI2*. Although the number of non-canonical splice junctions is relatively low (human: 0.04%, mouse: 0.09%), the **b)** majority of non-canonical splice junctions (n = 31) were supported by RNA-Seq with 17 splice junctions unique to mouse and 4 splice junctions unique to human. To accurately infer splice site identity, *STAR* output from matched RNA-Seq data from human fetal cortex (n = 3 biologically independent samples), a subset of the human cortex Iso-Seq data, and mouse cortex (n = 8 biologically independent samples) was used rather than Intropolis junction dataset.

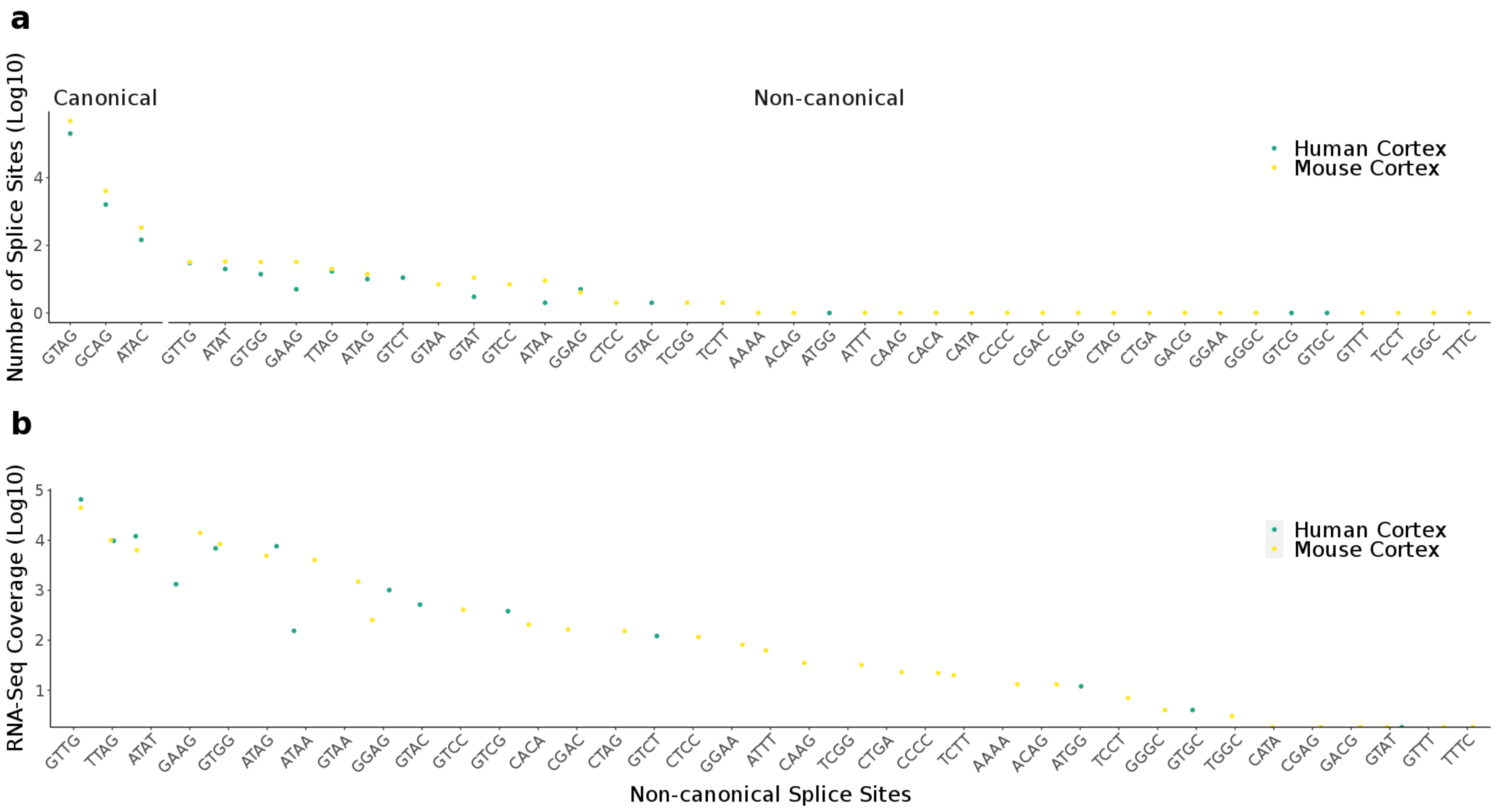

**Supplementary Figure 36: The majority of genes are characterized by more than one isoform in human adult and fetal cortex.** Shown is the distribution of isoform numbers identified for each detected gene in the human adult (n = 4 biologically independent samples) and human fetal (n = 3 biologically independent samples) cortex.

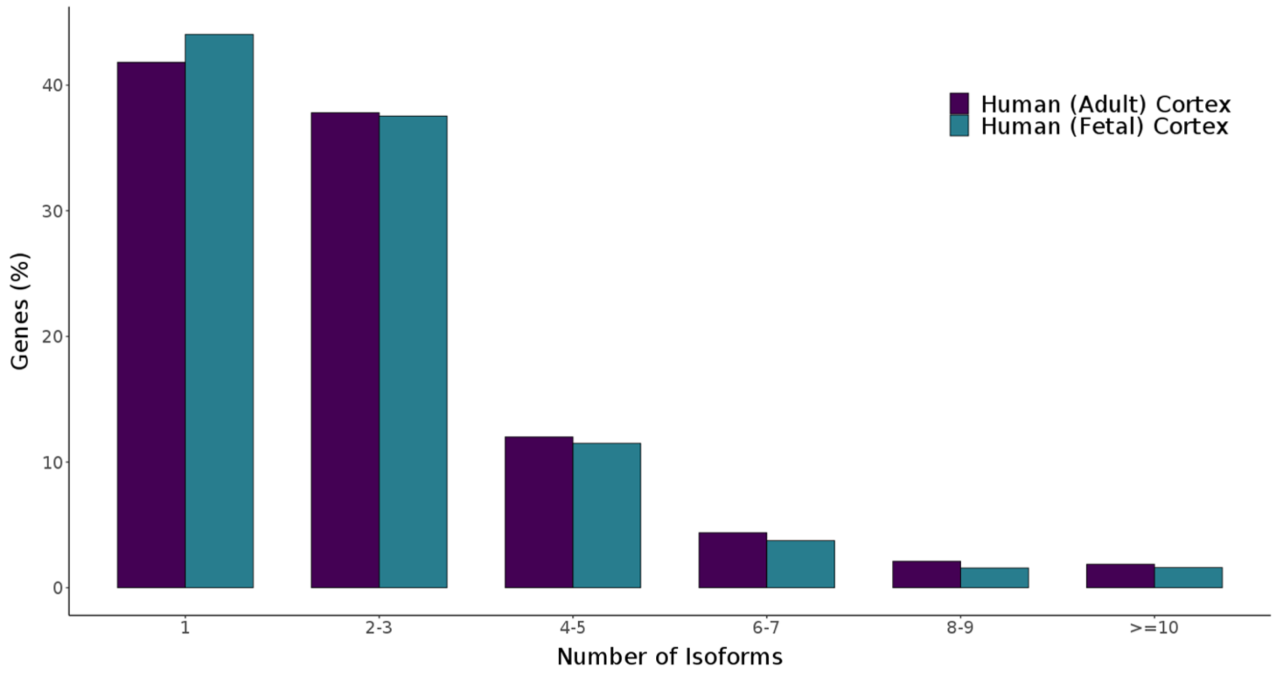

**Supplementary Figure 37: Widespread isoform diversity between human adult and human fetal cortex.** Shown is UCSC genome browser track for **a)** *MEG3* (adult cortex = 72 multi-exonic isoforms, fetal cortex = 34 multi-exonic isoforms) and **b)** *MBP* (adult cortex = 32 multi-exonic isoforms, fetal cortex = 2 multi-exonic isoforms), selected for having the largest absolute difference in isoform numbers between human fetal and human adult. Isoforms are coloured based on *SQANTI2* classification categories (blue = FSM, cyan = ISM, red = NIC, orange = NNC). FSM – Full Splice Match, ISM – Incomplete Splice Match, NIC – Novel In Catalogue, NNC – Novel Not in Catalogue

**a**

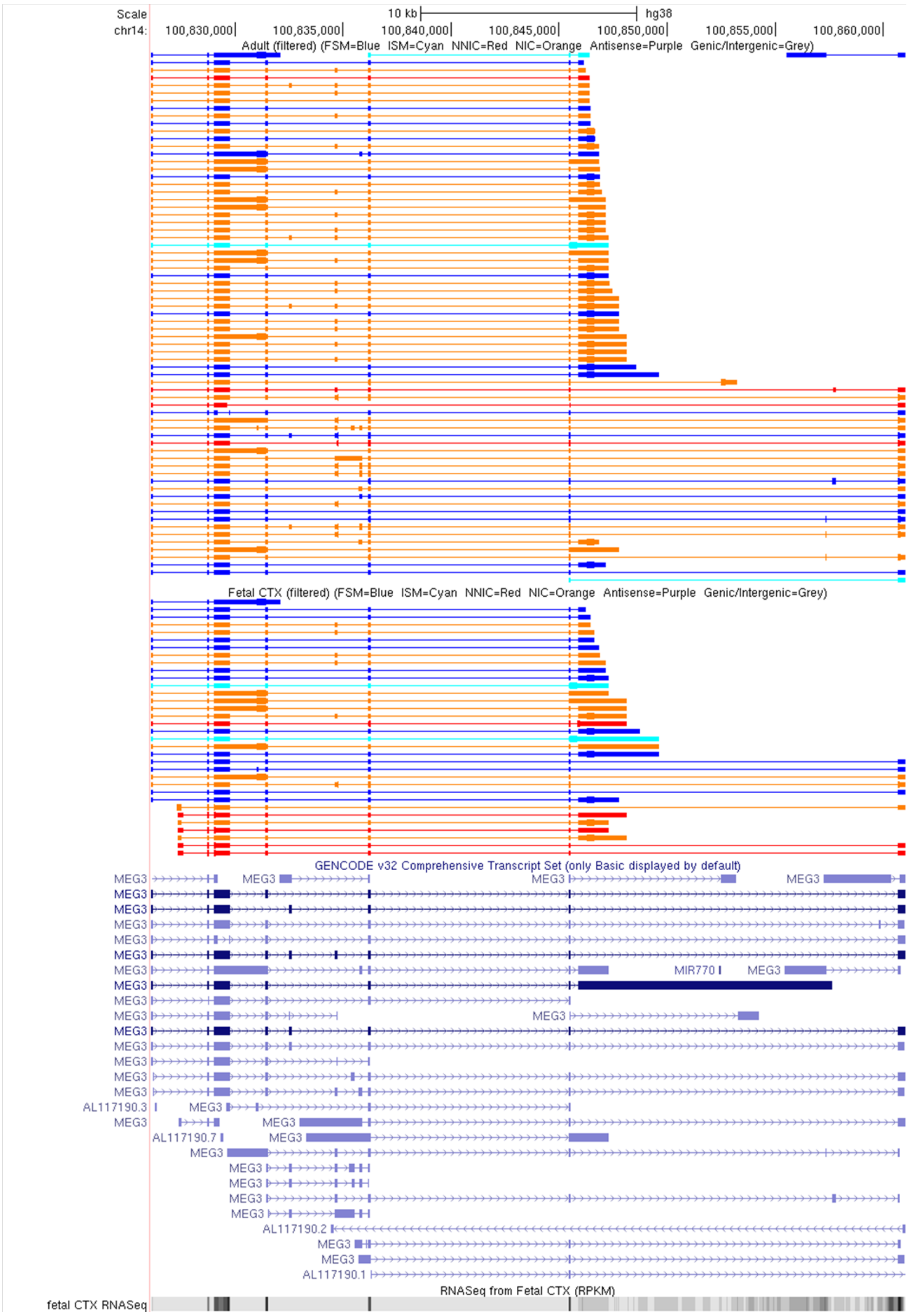

**b**

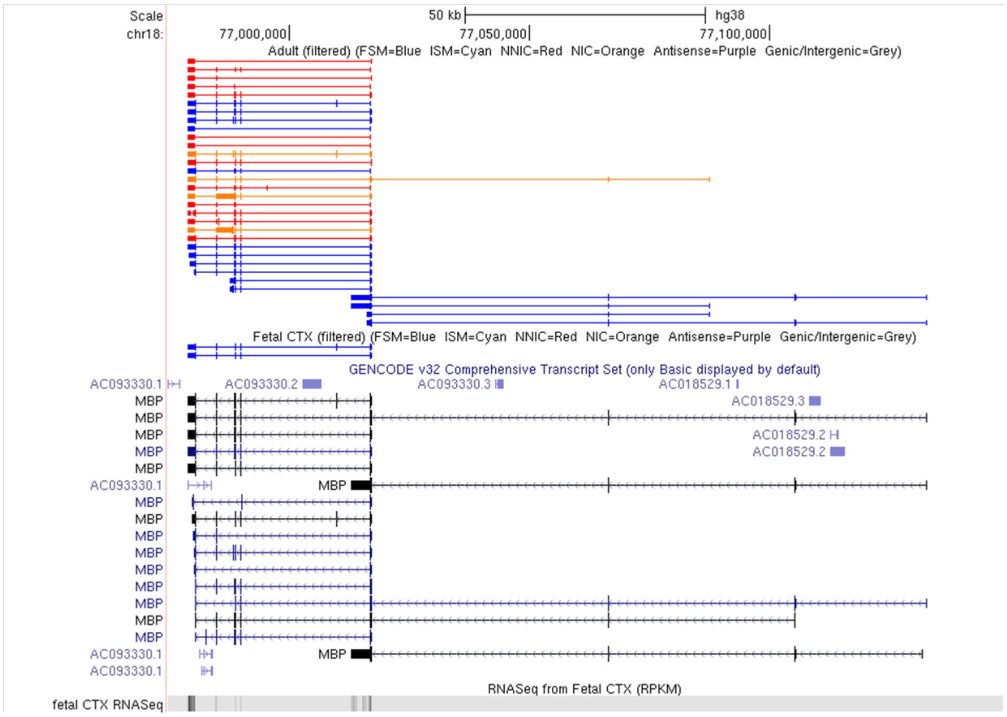
Scale
chr18:
50 kb
hg38
77,000,000
77,050,000
77,100,000
Adult (filtered) (FSM=Blue ISM=Cyan NNIC=Red NIC=Orange Antisense=Purple Genic/Intergenic=Grey)
Fetal CTX (filtered) (FSM=Blue ISM=Cyan NNIC=Red NIC=Orange Antisense=Purple Genic/Intergenic=Grey)
GENCODE v32 Comprehensive Transcript Set (only Basic displayed by default)
AC093330.1
AC093330.2
AC093330.3
AC018529.1
AC018529.3
AC018529.2
MBP
RNASeq from Fetal CTX (RPKM)
fetal CTX RNASeq

**Supplementary Figure 38: Greatest differential transcript usage observed in gene *MAP1B* between human adult and fetal cortex.** Shown is UCSC genome browser track for *MAP1B* for human cortex (n = 7 biologically independent samples) and human reference genome (hg38). *MAP1B* is characterised by largest expression difference in dominant transcript between adult (coloured in blue, n = 4 biologically independent samples) and fetal (coloured in red, n = 3 biologically independent samples) specific isoform. Both isoforms are identical in exonic structure with only difference at 3'UTR (boxed).

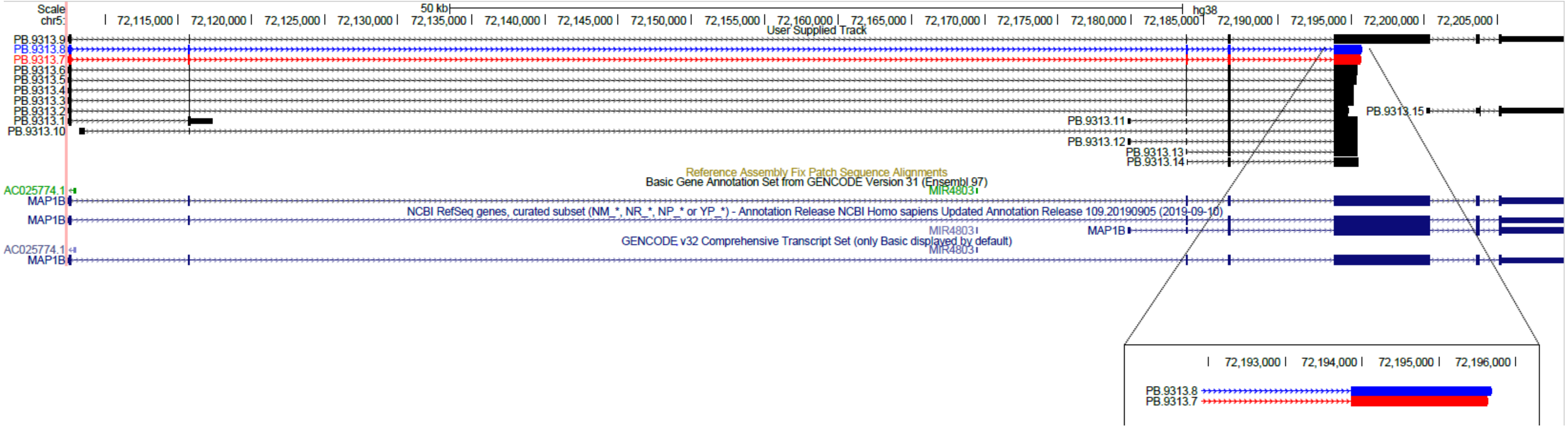

**Supplementary Figure 39: Differential transcript usage observed in gene *SNAP25* between human adult and fetal cortex.** Shown is UCSC genome browser track for *SNAP25* for human cortex (n = 7 biologically independent samples) and human reference genome (hg38). Differential transcript usage is observed in this gene, with one isoform (coloured in red) strongly expressed in fetal cortex while downregulated in adult cortex, and another isoform (coloured in blue) strongly expressed in adult cortex. Both isoforms consist of 8 exons, with the only difference at alternative exon 5 (highlighted in green).

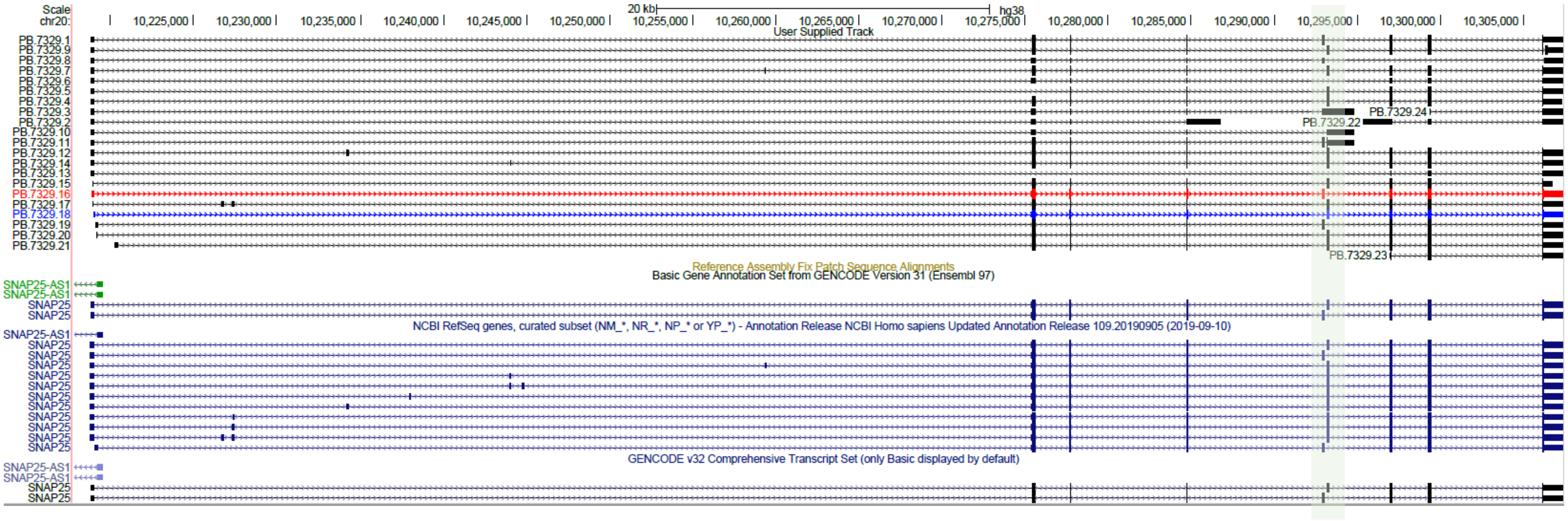

**Supplementary Figure 40: Genes with IR-transcripts were more highly expressed in human fetal cortex than adult cortex.** Shown is the Iso-Seq gene expression of genes with IR-transcripts in human adult (n = 4 biologically independent samples) and fetal (n = 3 biologically independent samples) cortex. IR-genes were more highly expressed in the fetal cortex than the adult cortex (Mann-Whitney-Wilcoxon test, $W = 1.1 \times 10^6$, $P = 2.87 \times 10^{-8}$). IR – Intron retention.

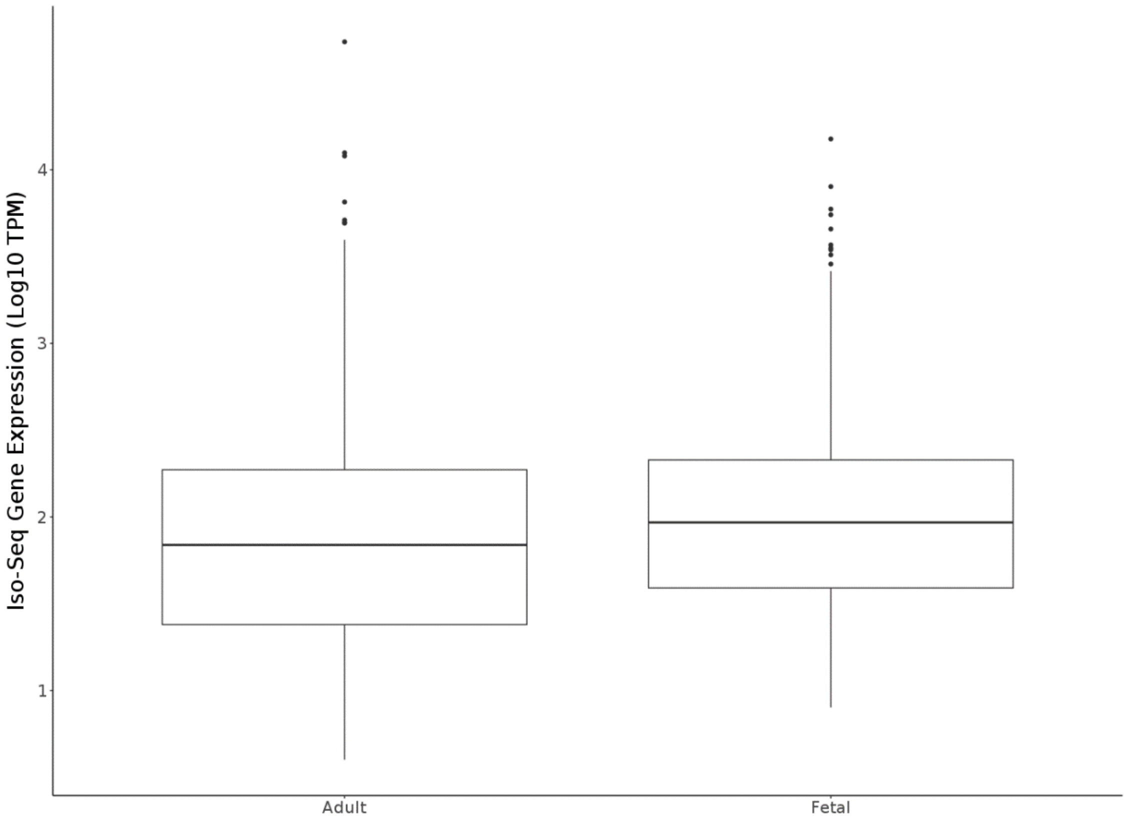

**Supplementary Figure 41: There was considerable overlap in detected genes across the three fetal brain regions, and between fetal and adult cortex.** Venn diagram showing overlap of **a)** fetal brain regions, and **b)** fetal cortex vs adult cortex, transcripts and genes (bracketed) with a TPM > 20. Exclusive expression in a tissue is defined as TPM = 0 in other developmental sample group or other regions. CTX – Cortex, HIP – Hippocampus, STR – Striatum. TPM – Transcripts per Million

**a**

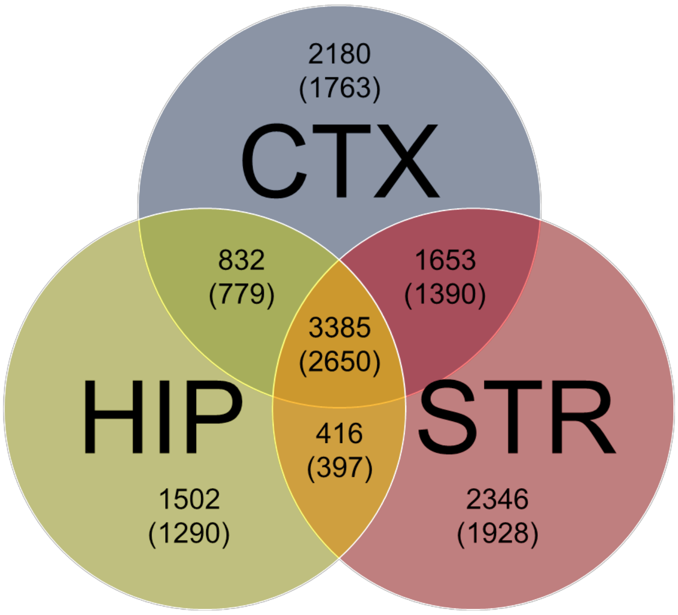


**b**

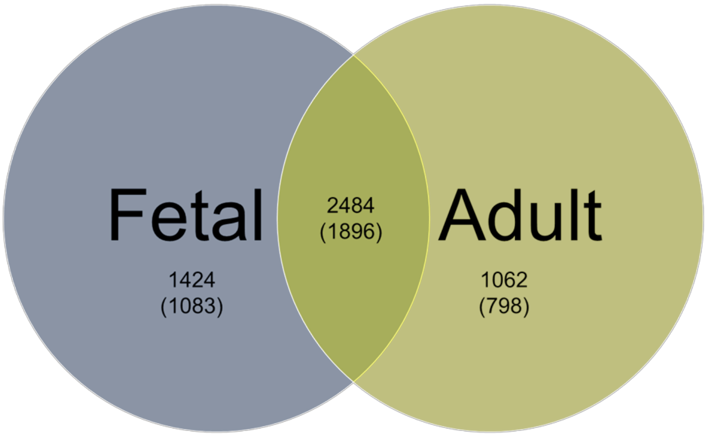

**Supplementary Figure 42: Differential transcript usage observed in gene *MEF2C* between fetal cortex and hippocampus.** Shown is UCSC genome browser track for *MEF2C* for human cortex (n = 7 biologically independent samples) and human reference genome (hg38). Alternative isoforms for this gene were detected in the fetal cortex (coloured in red, n = 3 biologically independent samples) and in the fetal hippocampus (coloured in blue, n = 2 biologically independent samples). Both isoforms consist of 11 exons with exons 1-10 being identical; however, the alternative isoform in hippocampus (~7kb) is significantly longer than the isoform in the cortex (~2kb) due to a shorter exon 11 and significantly shorter 3'UTR (highlighted in green).

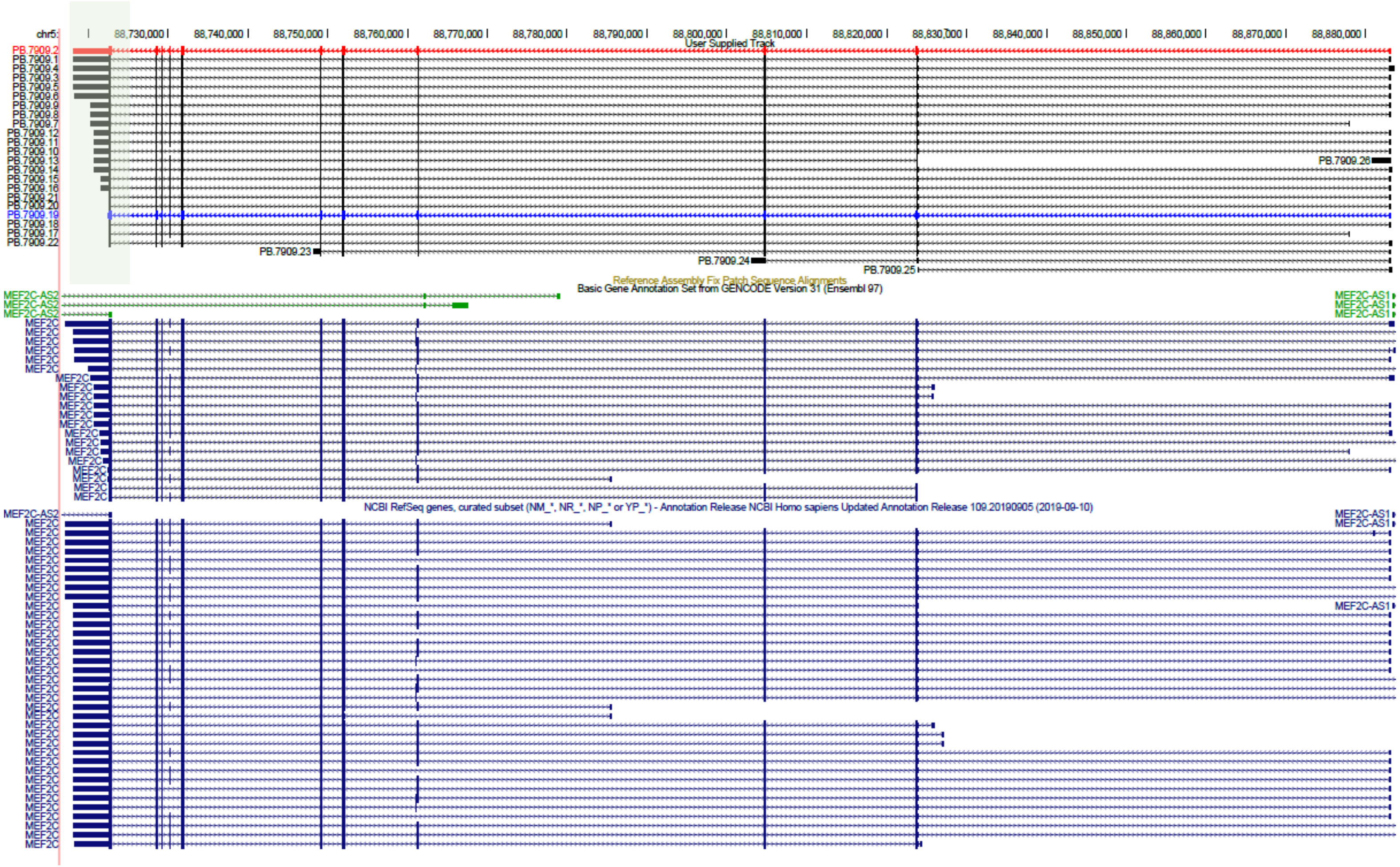

**Supplementary Figure 43: Additional examples of fusion transcripts with shared exons to genes associated with autism and schizophrenia in human cortex**. Shown are UCSC genome browser tracks of fusion transcripts of a disease-associated gene and other adjacent genes. Isoforms are coloured based on *SQANTI2* classification categories (blue = FSM, cyan = ISM, red = NIC, orange = NNC). FSM – Full Splice Match, ISM – Incomplete Splice Match, NIC – Novel In Catalogue, NNC – Novel Not in Catalogue

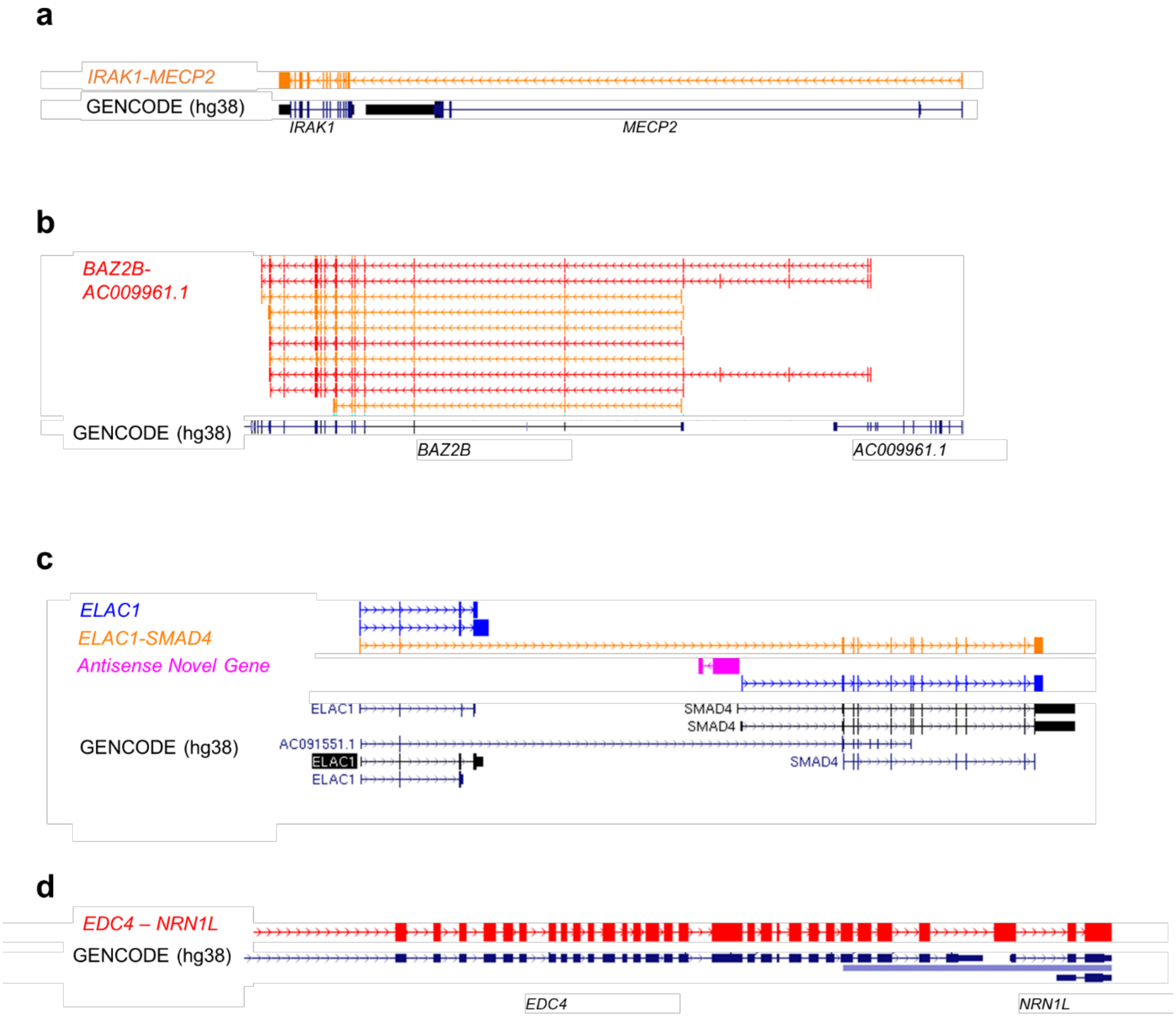
