## Supplementary Tables for "Full-length transcript sequencing of human and mouse identifies widespread isoform diversity and alternative splicing in the cerebral cortex"


| | |
|---|---|
| Supplementary Table 1 | Phenotypic information across samples |
| Supplementary Table 2 | Number and percentage of CCS generated |
| Supplementary Table 3 | SQANTI2 filtered transcripts annotation classification files |
| Supplementary Table 4 | Gene ontology results |
| Supplementary Table 5 | RNA-Seq mapping output |
| Supplementary Table 6 | Number of multi-exonic Isoforms in human and mouse cortex |
| Supplementary Table 7 | Summary table of Iso-Seq human and mouse cortical transcriptome |
| Supplementary Table 8 | Number of isoforms of annotated genes |
| Supplementary Table 9 | Summary information of novel genes |
| Supplementary Table 10 | Blast analysis of novel genes against genome |
| Supplementary Table 11 | Blast analysis of novel genes cross species |
| Supplementary Table 12 | AS events and genes with AS |
| Supplementary Table 13 | List of genes with intron retention |
| Supplementary Table 14 | List of genes with only IR-transcripts |
| Supplementary Table 15 | Number of isoforms in human and mouse cortex |
| Supplementary Table 16 | Differential transcript usage between human and adult fetal cortex |
| Supplementary Table 17 | Summary table of Iso-Seq human fetal hippocampus and striatum |
| Supplementary Table 18 | Differential transcript usage between different fetal brain regions |
| Supplementary Table 19 | Summary of disease-associated genes in human and mouse cortex |
| Supplementary Table 20 | Number of fusion and IR-transcripts of disease-associated genes |
| Supplementary Table 21 | Determining a high gene expression threshold |

**Supplementary Table 1: Phenotypic information across all samples.** Of note, fetal hippocampus and striatum samples were derived from the same donor as the fetal cortex samples. SMRT – Single-molecule real-time, ONT – Oxford Nanopore Technology, RIN – RNA Integrity Number, wpc – weeks post-conception

| Sample ID | Number of SMRT Cells | Species | Tissue | Sex | Age | RIN | RNA-Seq | ONT |
|---|---|---|---|---|---|---|---|---|
| Adult A | 1 | Human | Cortex | F | 89years | 4.1 | No | No |
| Adult B | 1 | Human | Cortex | F | 89years | 6.3 | No | No |
| Adult C | 2 | Human | Cortex | M | 24years | 8 | No | Yes |
| Adult D | 1 | Human | Cortex | M | 45years | 7 | No | No |
| Fetal A | 1 | Human | Cortex | M | 17wpc | 8.3 | Yes | Yes |
| Fetal B | 2 | Human | Cortex | F | 17wpc | 7.2 | Yes | No |
| Fetal C | 2 | Human | Cortex | F | 14wpc | 6.1 | Yes | No |
| Fetal D | 1 | Human | Hippocampus | F | 17wpc | 7.5 | No | No |
| Fetal E | 1 | Human | Hippocampus | F | 14wpc | 7.1 | No | No |
| Fetal F | 1 | Human | Striatum | F | 17wpc | 8.1 | No | No |
| Fetal G | 1 | Human | Striatum | F | 14wpc | 6.9 | No | No |
| Mouse A | 1 | Mouse | Cortex | F | 12months | 8.8 | Yes | No |
| Mouse B | 1 | Mouse | Cortex | F | 6months | 8.8 | Yes | No |
| Mouse C | 1 | Mouse | Cortex | F | 2months | 9.2 | Yes | No |
| Mouse D | 1 | Mouse | Cortex | F | 8months | 9.1 | Yes | No |
| Mouse E | 1 | Mouse | Cortex | F | 2months | 9.2 | Yes | No |
| Mouse F | 1 | Mouse | Cortex | F | 8months | 9 | Yes | No |
| Mouse G | 1 | Mouse | Cortex | F | 2months | 9.2 | Yes | No |
| Mouse H | 1 | Mouse | Cortex | F | 8months | 9.1 | Yes | No |

**Supplementary Table 2: Number and percentage of successful CCS reads generated across samples.** Distribution of CCS read lengths across all cortical samples can be found in **Supplementary Figure 1.** CCS - Circular consensus sequence

| Run ID | Sample ID | Species | Tissue | Number of CCS generated |
|---|---|---|---|---|
| Adult 1 | Adult A | Human | Cortex | 525054 |
| Adult 2 | Adult B | Human | Cortex | 550191 |
| Adult 3 | Adult C | Human | Cortex | 413417 |
| Adult 4 | Adult C | Human | Cortex | 493645 |
| Adult 5 | Adult D | Human | Cortex | 458028 |
| Fetal 1 | Fetal A | Human | Cortex | 266888 |
| Fetal 2 | Fetal B | Human | Cortex | 601083 |
| Fetal 3 | Fetal B | Human | Cortex | 517525 |
| Fetal 4 | Fetal C | Human | Cortex | 765092 |
| Fetal 5 | Fetal C | Human | Cortex | 546524 |
| Fetal 6 | Fetal D | Human | Hippocampus | 247015 |
| Fetal 7 | Fetal E | Human | Hippocampus | 235538 |
| Fetal 8 | Fetal F | Human | Striatum | 319831 |
| Fetal 9 | Fetal G | Human | Striatum | 227667 |
| Mouse 1 | Mouse A | Mouse | Cortex | 593822 |
| Mouse 2 | Mouse B | Mouse | Cortex | 412406 |
| Mouse 3 | Mouse C | Mouse | Cortex | 552414 |
| Mouse 4 | Mouse D | Mouse | Cortex | 569050 |
| Mouse 5 | Mouse E | Mouse | Cortex | 523422 |
| Mouse 6 | Mouse F | Mouse | Cortex | 580940 |
| Mouse 7 | Mouse G | Mouse | Cortex | 511273 |
| Mouse 8 | Mouse H | Mouse | Cortex | 575046 |

**Supplementary Table 3: SQANTI2 annotation of filtered transcripts of human and mouse transcriptome.**

See attached xlsx file SupplementaryTable3_SQANTI2Annotations.xlsx. Each tab refers to Iso-Seq dataset: HumanCTX – Human cortex (n = 7 biologically independent samples), AdultCTX – Adult cortex (n = 4 biologically independent samples), FetalCTX – Fetal cortex (n = 3 biologically independent samples), FetalHIP – Fetal hippocampus (n = 2 biologically independent samples), FetalSTR – Fetal striatum (n = 2 biologically independent samples), MouseCTX – Mouse cortex (n = 8 biologically independent samples).

**Supplementary Table 4: Gene ontology (GO) Results**

See attached xlsx file SupplementaryTable4_GOResults.xlsx. Three sets of GO were performed on human (n = 7 biologically independent samples) and mouse cortex (n = 8 biologically independent samples); i) 500 most abundantly expressed genes (ranked by TPM), finding most significant enrichment in ‘prefrontal cortex’ from Human Gene Atlas database and ‘prefrontal cerebral cortex’ from Mouse Gene Atlas database, ii) 100 most isoformic genes, finding most significant enrichment in ‘RNA binding’ for molecular function for both human cortex and mouse cortex, and enrichment in relevant GWAS datasets for Alzheimer’s disease, autism and schizophrenia, iii) Genes with IR-transcripts, finding most significant enrichment in ‘mRNA splicing’ in human cortex and mouse cortex, and ‘RNA binding’ unique to human fetal cortex for biological process. IR – Intron Retention, TPM – Transcripts per Million.

**Supplementary Table 5: Number of mapped reads from RNA-Seq data**

See attached xlsx file SupplementaryTable5_RNASeqMapping.xlsx. Short-read RNA-Seq data was generated on human fetal cortex (n = 3 biologically independent samples) and mouse cortex (n = 8 biologically independent samples), and then mapped to reference genome (human: hg38, mouse: mm10) using *STAR*.

**Supplementary Table 6: Number of isoforms in human and mouse cortex.**

See attached xlsx file SupplementaryTable6_IsoformNumHumanMouse.xlsx. Isoform diversity from Iso-Seq dataset of human (n = 7 biologically independent samples) and mouse (n = 8 biologically independent samples) is reported in three-fold: i) total number of isoforms identified per gene, normalised to the number of isoforms known in reference genome (human: hg38, mouse: mm10), present in both human and mouse cortex, ii) number of multi-exonic isoforms identified per gene, present in both human and mouse cortex, not accounting and iii) accounting for homology. Homology was considered by converting the mouse gene names to the equivalent homologous human gene names according to mouse genome informatics syntenic gene list (http://www.informatics.jax.org/downloads/reports/HOM_MouseHumanSequence.rpt) (see **Online Methods**).

**Supplementary Table 7: Summary table of Iso-Seq human and mouse cortical transcriptome.**
lncRNA – long non-coding RNA, FSM – Full Splice Match, ISM – Incomplete Splice Match, NIC – Novel In Catalogue, NNC – Novel Not in Catalogue, NMD – Nonsense-mediated mRNA decay

| | Description | Human Cortex | Adult Cortex | Fetal Cortex | Mouse Cortex |
|---|---|---|---|---|---|
| | Unique Genes | 12885 | 10979 | 9667 | 13590 |
| | Annotated Genes | 12832(99.59%) | 10949(99.73%) | 9647(99.79%) | 13450(98.97%) |
| | Novel Genes | 53 (0.41%) | 30 (0.27%) | 20 (0.21%) | 140 (1.03%) |
| Annotated Genes | Transcripts | 42585 | 27842 | 23191 | 51003 |
| | Protein-coding Transcripts | 39352(92.41%) | 25690(92.27%) | 21763(93.84%) | 47757(93.64%) |
| | Genes associated with coding transcripts | 11959(93.17%) | 10234 (93.47%) | 9168 (95.03%) | 12748 (94.78%) |
| | Non-lncRNA Transcripts | 41040 | 26905 | 22507 | 49962 |
| | lncRNA Transcripts | 1545 | 937 | 684 | 1041 |
| | Mono-exonic non lncRNA | 1021 (2.49%) | 722 (2.68%) | 541 (2.4%) | 1325 (2.65%) |
| | Mono-exonic lncRNA Transcripts | 533 (34.5%) | 351 (37.46%) | 250 (36.55%) | 282 (27.09%) |
| | Genes associated with lncRNA Transcripts | 829 | 574 | 381 | 587 |
| | Protein-coding lncRNA Transcripts | 601(38.9%) | 350(37.35%) | 271(39.62%) | 413(39.67%) |
| | Annotated Transcripts (FSM, ISM) | 28654(67.29%) | 20451(73.45%) | 16947(73.08%) | 32664(64.04%) |
| | Novel Transcripts | 13931(32.71%) | 7391(26.55%) | 6244(26.92%) | 18339(35.96%) |
| | FSM | 23629 | 16971 | 14796 | 27475 |
| | ISM | 5025 | 3480 | 2151 | 5189 |
| | NIC | 9724 | 4903 | 4730 | 10649 |
| | NNC | 4013 | 2380 | 1447 | 7414 |
| | Genic Genomic | 41 | 23 | 13 | 57 |
| | Antisense | 0 | 0 | 0 | 0 |
| | Fusion | 153 | 85 | 54 | 219 |
| | Intergenic | 0 | 0 | 0 | 0 |
| | Genic Intron | 0 | 0 | 0 | 0 |
| | Genes associated with Novel Transcripts | 5622 (43.81%) | 3682 (33.63%) | 3219 (33.37%) | 6694 (49.77%) |
| | Genes associated with Annotated Transcripts | 11985 (93.4%) | 10248 (93.6%) | 9026 (93.56%) | 12560 (93.38%) |
| | NMD Transcripts | 5062 (11.89%) | 2690 (9.66%) | 2533 (10.92%) | 4944 (9.69%) |
| | Genes with NMD transcripts | 1483 (13.54%) | 1415 (14.67%) | 2420 (18.86%) | 2264 (16.83%) |

| | | | | |
|---|---|---|---|---|
| Fusion Genes | 114 (0.88%) | 61 (0.56%) | 47 (0.49%) | 160 (1.19%) |
| Transcripts of Fusion Genes | 153 (0.36%) | 85 (0.3%) | 54 (0.23%) | 219 (0.43%) |
| Fusion genes with more than one transcript | 23 (20.18%) | 15 (24.59%) | 5 (10.64%) | 40 (25%) |
| Transcripts with Intron Retention | 5752 (13.49%) | 2557 (9.17%) | 3053 (13.15%) | 4216 (8.24%) |
| Genes with Intron Retention | 2625 (20.37%) | 1435 (13.07%) | 1662 (17.19%) | 2279 (16.77%) |
| Protein-coding, IR-transcripts | 95.31% | 95.19% | 95.48% | 95.56% |
| IR-transcripts with canonical splice junctions | 100% | 100% | 100% | 99.60% |

**Supplementary Table 8: Number of isoforms of annotated genes in human and mouse cortex.**

See attached xlsx file SupplementaryTable8_AnnotatedGenes.xlsx**.** Listed are the number of novel and known multi-exonic isoforms of annotated gees in human cortex (n = 7 biologically independent samples), human adult cortex (n = 4 biologically independent samples), human fetal cortex (n = 3 biologically independent samples), and mouse cortex (n = 8 biologically independent samples).

**Supplementary Table 9: Summary information of novel genes in human and mouse cortex.**

See attached xlsx file SupplementaryTable9_NovelGenes.xlsx. Listed are the novel genes, not previously known in existing reference genome databases (human: hg38, mouse: mm10), for human cortex (n = 7 biologically independent samples), human adult cortex (n = 4 biologically independent samples), human fetal cortex (n = 3 biologically independent samples), and mouse cortex (n = 8 biologically independent samples). Each novel gene is provided with the following information: genomic locus, number of full-length reads (FL) associated, gene name assigned by *SQANTI2*, protein-coding potential, whether RNA-Seq supported, whether located near a CAGE peak defined by *SQANTI2*, and whether identified as novel gene in GTEx consortium (CHESS v2.2 annotation).

**Supplementary Table 10: BLAST analysis of novel genes against genome to identify homology with other genomic regions**

See attached xlsx file SupplementaryTable10_BlastNovelGenes.xlsx. BLAST analysis was performed between novel genes from human cortex (n = 7 biologically independent samples) and human reference genome (hg38), and between novel genes from mouse cortex (n = 8 biologically independent samples) and mouse reference genome (mm10) to identify homology with other genomic regions. Listed are the novel genes that had a BLAST hit after filtering (longer than 500bp, more than 90% identity) and ensuring a different genomic locus.

**Supplementary Table 11: BLAST analysis of human and mouse novel genes identified one common novel gene.** BLAST analysis was performed between mouse novel genes (n = 156 novel transcripts mapping to 131 novel genes) and human novel genes (n = 60 novel transcripts mapping to 49 novel genes), with one BLAST hit referring to a common novel gene overlapping - and antisense to - *E2F3*. Both PB.21939.1 and PB.3680.1 isoform were assigned as "novelGene_E2F3_AS" by *SQANTI2* in human and mouse cortex (**Supplementary Table 3, Supplementary Table 9**). UCSC genome browser track of this common novel gene can be found in **Figure 4**.

| Human PB_ID | Mouse PB_ID | % Identity | Alignment Length | Mismatches | Gap | Human Start | Human End | Mouse Start | Mouse End | E-value | Bit Score |
|---|---|---|---|---|---|---|---|---|---|---|---|
| PB.21939.1 | PB.3680.1 | 94 | 50 | 1 | 2 | 2 | 49 | 565 | 614 | 3.23E-16 | 75 |
| PB.21939.1 | PB.3680.1 | 83.074 | 1926 | 205 | 60 | 2 | 1889 | 565 | 2407 | 0 | 1639 |

**Supplementary Table 12: Alternative splicing events observed in human and mouse cortex**. Tabulated are **a)** number of splicing events and **b)** number of genes observed with those splicing event, in human cortex (n = 7 biologically independent samples), human adult cortex (n = 4 biologically independent samples), human fetal cortex (n = 3 biologically independent samples), and mouse cortex (n = 8 biologically independent samples). Of note, a single gene can be characterised by multiple splicing events, and can thus appear more than once in b). A combination of the *SUPPA2* package and custom analysis scripts (see **Online Methods**) were used to identify transcripts associated with i) exon skipping (SE), ii) mutually exclusive exon use (MX), iii) alternative first (AF) and last (AL) exons, iv) alternative 3' and 5' splice sites, and v) intron retention (IR).

**a)**

| Splicing event | Number and proportion of splicing events | | | |
|---|---|---|---|---|
| | **Human Cortex** | **Human (Adult) Cortex** | **Human (Fetal) Cortex** | **Mouse Cortex** |
| A3 | 2146 (6.35%) | 1240 (6.4%) | 907 (5.89%) | 2798 (7.09%) |
| A5 | 1184 (3.5%) | 678 (3.5%) | 471 (3.06%) | 1672 (4.23%) |
| AF | 11133 (32.95%) | 7265 (37.49%) | 4595 (29.83%) | 14111 (35.74%) |
| AL | 7262 (21.49%) | 4288 (22.13%) | 3529 (22.91%) | 9883 (25.03%) |
| IR | 5752 (17.02%) | 2557 (13.19%) | 3053 (19.82%) | 4216 (10.68%) |
| MX | 355 (1.05%) | 167 (0.86%) | 118 (0.77%) | 415 (1.05%) |
| SE | 5956 (17.63%) | 3184 (16.43%) | 2732 (17.73%) | 6391 (16.19%) |

**b)**

| Splicing event | Number and proportion of genes with splicing events | | | |
|---|---|---|---|---|
| | **Human Cortex** | **Human (Adult) Cortex** | **Human (Fetal) Cortex** | **Mouse Cortex** |
| A3 | 1198 (9.88%) | 757 (7.27%) | 600 (6.51%) | 1487 (11.45%) |
| A5 | 728 (6%) | 461 (4.43%) | 330 (3.58%) | 930 (7.16%) |
| AF | 5307 (43.75%) | 3869 (37.16%) | 2843 (30.84%) | 6089 (46.9%) |
| AL | 4272 (35.22%) | 2828 (27.16%) | 2347 (25.46%) | 5054 (38.93%) |
| IR | 2625 (21.64%) | 1435 (13.78%) | 1662 (18.03%) | 2279 (17.55%) |
| MX | 278 (2.29%) | 136 (1.31%) | 104 (1.13%) | 303 (2.33%) |
| SE | 3482 (28.71%) | 2097 (20.14%) | 1866 (20.24%) | 3662 (28.21%) |

**Supplementary Table 13: Genes with intron retained transcripts in human and mouse cortex.**

See attached xlsx file SupplementaryTable13_IntronRetention.xlsx. Tabulated are the total number of IR-transcripts detected per gene in human cortex (n = 7 biologically independent samples), human adult cortex (n = 4 biologically independent samples), human fetal cortex (n = 3 biologically independent samples), and mouse cortex (n = 8 biologically independent samples). IR – Intron retention.

**Supplementary Table 14: Genes with only intron retained transcripts in human and mouse cortex.**

See attached xlsx file SupplementaryTable14_IRTranscriptsOnly.xlsx. Tabulated are the list of genes found to *only* express transcripts characterised with IR in human cortex (n = 7 biologically independent samples), human adult cortex (n = 4 biologically independent samples), human fetal cortex (n = 3 biologically independent samples), and mouse cortex (n = 8 biologically independent samples). IR – Intron retention.

**Supplementary Table 15: Number of isoforms in human adult and human fetal cortex.**

See attached xlsx file SupplementaryTable15_IsoformNumHumanAdultFetal.xlsx. Isoform diversity from Iso-Seq dataset of human adult (n = 4 biologically independent samples) and human fetal (n = 3 biologically independent samples) is reported in three-fold: i) total number of isoforms identified per gene, normalised to the number of isoforms known in reference genome (human: hg38), identified in human adult and human fetal cortex ii) number of multi-exonic isoforms identified per gene and iii) number of multi-exonic isoforms identified of genes identified in both datasets.

**Supplementary Table 16: Differential transcript usage between human and adult fetal cortex.**

See attached xlsx file SupplementaryTable16_DtuHumanAdultFetal.xlsx

**Supplementary Table 17: Summary table of Iso-Seq human fetal hippocampus and human fetal striatum Iso-Seq dataset.** FSM – Full Splice Match, ISM – Incomplete Splice Match, NIC – Novel In Catalogue, NNC – Novel Not in Catalogue

| Description | Fetal Hippocampus | Fetal Striatum |
|---|---|---|
| Unique Genes | 5606 | 6035 |
| Annotated Genes | 5604 (99.96%) | 6028 (99.88%) |
| Novel Genes | 2 (0.04%) | 7 (0.12%) |
| Transcripts | 8416 | 9678 |
| Annotated Transcripts | 7261 (86.28%) | 8118 (83.88%) |
| Novel Transcripts | 1155 (13.72%) | 1560 (16.12%) |
| FSM | 6720 | 7423 |
| ISM | 541 | 695 |
| NIC | 911 | 1184 |
| NNC | 235 | 357 |
| Genic Genomic | 0 | 3 |
| Antisense | 1 | 3 |
| Fusion | 7 | 9 |
| Intergenic | 1 | 4 |
| Genic Intron | 0 | 0 |

**Supplementary Table 18: Genes identified as showing differential transcript usage across different fetal brain regions.** CTX – Cortex, HIP – Hippocampus, STR - Striatum

| **CTX-HIP** | **CTX-STR** | **HIP-STR** |
|---|---|---|
| CSE1L | AZIN1 | CSE1L |
| ELAVL3 | CNTFR | GNAS |
| H2AFY | COPS3 | H2AFY |
| MEF2C | CSE1L | RPL15 |
| TUBB3 | DUSP26 | SMARCB1 |
| | EFNB1 | TUBA1A |
| | EWSR1 | |
| | H2AFY | |
| | MAGED4B | |
| | MKRN2 | |
| | MSN | |
| | NUSAP1 | |
| | PHF20L1 | |
| | PSIP1 | |
| | R3HCC1 | |
| | RSRC2 | |
| | SLC25A6 | |
| | TCEA1 | |
| | UBAP1 | |

**Supplementary Table 19: Summary table of disease-associated genes in human and mouse cortex.** Isoform diversity was assessed in genes robustly associated with autism (393 genes nominated as being category 1 (high confidence) and category 2 (strong candidate) from the SFARI Gene database https://gene.sfari.org/), Alzheimer's disease (three familial AD genes and 59 genes nominated from the most recent GWAS meta-analysis) and schizophrenia (SZ) (339 genes nominated from the most recent GWAS meta-analysis). AD – Alzheimer's disease, SZ – Schizophrenia. IR – Intron retention, NMD – Nonsense-mediated mRNA decay, FSM – Full Splice Match, ISM – Incomplete Splice Match, NIC – Novel In Catalogue, NNC – Novel Not in Catalogue

| | Human Cortex | | | Mouse Cortex | | |
|---|---|---|---|---|---|---|
| **Description** | **AD** | **SZ** | **Autism** | **AD** | **SZ** | **Autism** |
| Disease-associated genes | 62 | 339 | 393 | 62 | 339 | 393 |
| Detected disease-associated genes ("Detected") | 31 (50%) | 281 (82.89%) | 307 (78.12%) | 39 (62.9%) | 319 (94.1%) | 329 (83.72%) |
| Total Number of Transcripts | 160 | 1200 | 1405 | 223 | 1592 | 2031 |
| Number and % of Annotated Transcripts | 100 (62.5%) | 736 (61.33%) | 935 (66.6%) | 131 (58.7%) | 917 (57.6%) | 1144 (56.3%) |
| Number and % of Novel Transcripts | 60 (37.5%) | 464 (38.67%) | 470 (33.5%) | 92 (41.3%) | 675 (42.4%) | 887(43.7%) |
| FSM | 65 | 569 | 592 | 104 | 727 | 753 |
| ISM | 35 | 167 | 343 | 27 | 190 | 391 |
| NIC | 48 | 341 | 346 | 52 | 418 | 607 |
| NNC | 12 | 123 | 124 | 40 | 257 | 280 |
| Genic Genomic | 0 | 0 | 0 | 0 | 0 | 0 |
| Antisense | 0 | 0 | 0 | 0 | 0 | 0 |
| Fusion | 0 | 0 | 0 | 0 | 0 | 0 |
| Intergenic | 0 | 0 | 0 | 0 | 0 | 0 |
| Genic Intron | 0 | 0 | 0 | 0 | 0 | 0 |
| IR Genes (% of all IR Genes) | 8 (0.3%) | 75 (2.86%) | 68 (2.59%) | 7 (0.31%) | 70 (3.07%) | 64 (2.81%) |
| IR Genes (% of Detected) | 8 (25.8%) | 75 (26.7%) | 68 (22.2%) | 7 (18.0%) | 70 (21.9%) | 64 (19.5%) |
| NMD Genes (% of Detected) | 8 (25.8%) | 58 (20.6%) | 52 (16.9%) | 6 (15.4%) | 61 (19.1%) | 61 (18.5%) |
| IR and NMD genes (% of Detected) | 5 (16.1%) | 26 (9.25%) | 27 (8.79%) | 2 (5.13%) | 30 (9.4%) | 22 (6.69%) |
| Fusion Genes | 1 | 3 | 4 | 0 | 4 | 4 |
| Number of Detected Genes with >1 isoform | 24 (77.4%) | 209 (74.4%) | 239 (77.9%) | 34 (87.2%) | 250 (78.4%) | 277 (84.2%) |

**Supplementary Table 20: Number of fusion and IR-transcripts of disease-associated genes**

See attached xlsx file SupplementaryTable20_FusionIRDisease.xlsx. Tabulated are the *SQANTI2* annotations of transcripts of disease-associated genes in each Iso-Seq dataset: human cortex (n = 7 biologically independent samples), human adult cortex (n = 4 biologically independent samples), human fetal cortex (n = 3 biologically independent samples), and mouse cortex (n = 8 biologically independent samples). The number of transcripts, IR-transcripts, IR-NMD-transcripts and fusion transcripts are detailed for each disease-associated gene across all Iso-Seq cortical datasets. IR – Intron retention, NMD – Nonsense-mediated mRNA decay, IR-NMD – intron retained transcripts predicted for nonsense-mediated mRNA decay.

**Supplementary Table 21: Determining a common high gene expression threshold between human and mouse**

A high gene expression threshold was applied to a few analyses to further understand the relationship between isoform number and gene length (**Supplementary Figure 10**), isoform number and gene exon number (**Supplementary Figure 11**), and to investigate whether there was a difference in intron retention rate between highly-expressed and lowly-expressed genes (**Figure 5**). A gene expression cut-off was sequentially applied to both human and mouse cortex Iso-Seq dataset, and the number of isoforms of the filtered genes were then correlated. Subsequently, the gene expression threshold was determined by the gene expression at which number of isoforms for commonly expressed genes was most correlated between human and mouse – in this case, 2.5 $Log_{10}$TPM. Of note, the genes filtered could have an expression surpassing threshold in mouse but not in human, and vice versa.  TPM – Transcripts per Million

| **Iso-Seq Gene Expression threshold ($Log_{10}$TPM)** | **Human-Mouse correlation of number of isoforms** | **P-value** | **Number of genes surpassing expression threshold** |
|---|---|---|---|
| 0 | 0.521 | 0 | 19646 |
| 0.5 | 0.516 | 0 | 18148 |
| 1 | 0.513 | 0 | 15229 |
| 1.5 | 0.512 | 0 | 10089 |
| 2 | 0.543 | 0 | 4273 |
| **2.5** | **0.606** | **$7.8 \times 10^{-115}$** | **1134** |
| 3 | 0.591 | $2.75 \times 10^{-20}$ | 201 |
| 3.5 | 0.522 | $8.87 \times 10^{-3}$ | 24 |
| 4 | 1 | NA | 2 |
